# ER-liquid condensate contacts sequester FAM134B/C and RhoA to govern cell morphology

**DOI:** 10.1101/2025.11.01.685922

**Authors:** Natalia Jimenez-Moreno, Arianna Karageorgiou, Katie Winnington-Ingram, Jimi Wills, Chinmayi Pednekar, Matthew Pearson, Lukas Gerasimavicius, Ann Wheeler, Joseph A. Marsh, Paul Verkade, Alex von Kriegsheim, Jon D. Lane, Simon Wilkinson

## Abstract

ER-phagy receptors have elusive physiological functions beyond ER remodelling. Here, we use proximity biotinylation to identify their cytoplasmic interactomes. Secondary CRISPR/Cas9 screening reveals regulators of the prototypical FAM134B/C receptors, which include PRKAR1A, canonically known as a subunit of PKA. PRKAR1A directly binds an amphipathic helix in the otherwise disordered cytoplasmic domain of FAM134B. Super-resolution, FRAP and CLEM imaging reveal novel interorganellar contacts between liquid-like condensates of cAMP-bound PRKAR1A and the ER. Condensates promote clustering of ER-embedded FAM134B/C with LC3B, independently of regulation of PKA by PRKAR1A. Proteomics reveal that cytoplasmic RhoA interacts with FAM134B/C clusters and that sequestration of both these molecules occurs within lysosomes embedded within the proximal condensates. This results in reduced actomyosin contractility and ER-condensate interactions thusly determine cell morphology and cancer cell invasion modality. In summary, ER-condensate contacts mediated by FAM134B/C are novel cellular degradation hubs that coordinate ER and cellular remodelling.

## Introduction

The endoplasmic reticulum (ER) is an extensive endomembrane network within the cytoplasm ^1^. While classically functioning as a biosynthetic and calcium storage organelle in virtually all mammalian cell types, the ER is also emerging as a fundamental organising hub for downstream co-ordination of cytoplasmic activities. This is mediated via interorganellar contacts, consistent with its dynamic morphology and large, cytoplasm-facing membrane area. For instance, ER contacts control the behaviour of mitochondria, lysosomes, peroxisomes, and endosomes ^1^. However, our knowledge of these cytoplasmic functions of the ER remains incomplete. Conversely, ER-phagy is emerging as a powerful regulatory pathway acting to remodel the ER ^2–4^. In this process, membrane-embedded, cytoplasm-facing ER-phagy receptors first cluster into complexes with core autophagy proteins, such as ATG8s (LC3- and GABARAP-family members, e.g., LC3B), via linear LC3-interacting regions (LIRs) in their cytoplasmic (extra-luminal) domains ^5–13^. The next step of ER-phagy – and also a key feature of closely related degradation processes that also involve the receptors – is sequestration of ER content into lysosomes, frequently targeting specific ER proteins for turnover ^14–22^. Lysosomal sequestration occurs through encapsulation of ER content into intermediary double-membraned vesicles ^6, 10, 14, 23^ or various other, topologically-distinct events ^15, 22, 24–26^. Insufficiency in ER-phagy and related processes thus results in proteostatic defects and consequent disease phenotypes in model organisms ^6, 16, 27–30^. Consistent with this, ER-phagy is disrupted by inherited mutations in receptors or by other processes in human disease settings, such as type II hereditary sensory and autonomic neuropathy (HSAN), and pancreatic precancer states ^6, 16^. Canonically, ER-phagy receptors are activated in response to ER-centric proteostatic stress, downstream of the unfolded protein response (UPR), or non-specifically in response to nutrient deprivation, which acts upon multiple signalling pathways to modulate most forms of autophagy ^9, 31–38^.

Strikingly, however, analysis of the molecular structure of ER-phagy receptors suggests unexplored roles, beyond proteostasis. Receptors typically feature large cytoplasmic domains of broadly unknown function, frequently characterised by intrinsically-disordered regions (IDRs) ^11, 12, 39, 40^. IDRs have low tendency toward secondary structure but protein-protein interactions can favour particular conformations ^41^. Linear interaction motifs are also prevalent in IDRs ^41^. Half of the mass of the reticulon homology domain (RHD) receptors FAM134B and FAM134C (collectively, FAM134B/C) is an approximately 28 kDa C-terminal cytoplasmic domain characterised by extensive intrinsic disorder ^5, 6, 40, 42^. While this region contains the LIR and may also assist membrane remodelling ^39, 40^, the function of the cytoplasmic domain of FAM134B/C – and receptors in general – remains elusive, particularly with regard to the potential for physiologic integration of ER-phagy with broader cellular remodelling events, in line with the emergent cytoplasmic organisational roles of the ER.

IDRs also frequently facilitate the formation of liquid-like condensates (hereafter referred to as condensates) ^41^. This occurs during the phenomenon of liquid-liquid phase separation (LLPS); condensates are dynamically-assembled organelles exhibiting absent (or incomplete) membrane delimitation of their periphery ^43^. They exhibit rapid, diffusion-limited exchange of constituent proteins with the surrounding cytosol. Condensation can facilitate dynamic, signal-responsive control of cellular responses ^44^. For example, condensation serves to cluster core autophagy proteins and to target cytoplasmic moieties for degradation, particularly ubiquitinated protein aggregates ^45–53^. Relevant to this study, the IDR-containing protein and PKA (protein kinase A) constituent protein PRKAR1A forms condensates upon binding to cAMP ^54, 55^. This occurs concomitantly with PRKAR1A dissociation from PKA catalytic subunits (PKA_CAT_). Notably, cAMP generation occurs during a wide range of basal and induced cell signalling responses, and has profound effects on cellular state, including altered gene expression and cellular morphological changes ^56, 57^. Nevertheless, despite the emerging flexibility of liquid-like condensation as a cytoplasmic organising principle, it has been difficult to envisage involvement of liquid-like condensation in ER-phagy receptor function, given the integration of the latter into the ER membrane.

Here, in order to uncover cytoplasmic pathways that regulate ER-phagy and might potentially coordinate ER-phagy receptor mobilisation with wider cytoplasmic changes, we perform a proximity proteomic screen for extra-luminal interactors of the major receptors. Following on from a secondary, targeted CRISPR-Cas9 screen that identifies functional regulators of FAM134B/C, we demonstrate that dimeric PRKAR1A is a direct interactor of the extensive cytoplasmic region of FAM134B/C. This is facilitated through an amphipathic helix-forming region embedded within the broader C-terminal IDR of FAM134B/C. Mechanistically, cAMP-responsive condensates of PRKAR1A facilitate focal clustering of FAM134B/C with LC3B at novel contact sites with the ER. This generates lysosome-enriched degradation hubs where FAM134B/C is turned over. Importantly, this function of PRKAR1A is shown to be independent of canonical regulation of PKA_CAT_ enzymatic activity. Further proteomic analyses of cAMP-activated FAM134B identify an interaction with cytoplasmic RhoA. This is shown to result in RhoA lysosomal degradation alongside the receptor, leading to a reduction in cell peripheral actomyosin contractility. The physiological importance of this is demonstrated by experimental delineation of an essential requirement for FAM134B/C-PRKAR1A interaction in stereotypical cAMP-induced cell morphological changes and for a switch from collective to single cell invasion of cancer cells in three-dimensional matrices. Overall, we conclude that the novel ER-liquid condensate contact sites scaffolded by FAM134B/C and PRKAR1A are degradative hubs that integrate ER and cellular remodelling.

## Results

### Proximity proteomic screening identifies interactors of ER-phagy receptors

To identify cytoplasmic regulators of ER-phagy, we performed proximity proteomic analyses using five major ER membrane-embedded ER-phagy receptors as bait (FAM134B, FAM134C, SEC62, TEX264, CCPG1), along with the wholly cytoplasmic receptor CALCOCO1 for comparison. We expressed each tagged at cytoplasmic terminus(es) with TurboID (TID) ^58^ and V5 (Fig. 1A) and at levels comparable to endogenous (Fig. S1A). This was done in an established cell culture system we had previously reported as exhibiting basal engagement of FAM134B/C, TEX264 and CCPG1 activities, namely murine PDAC (pancreatic ductal adenocarcinoma) cells ^59^. TID-mediated biotinylation of bait-proximal proteins occurred upon biotin supplementation of live cells (Figs. 1B-D, S1B-C). Reference biotinylation samples were generated with two controls: cytoplasmic TID-NES (nuclear exclusion signal) and ER membrane-anchored ERM-TID (Figs. 1A-D, S1B-C) ^58^. Streptavidin (SA) was used to affinity-purify biotinylated species for each sample followed by mass spectrometric identification of prey proteins enriched versus controls (Figs. 1E, S1D, Table S1). These data revealed a substantive cohort of putative interactors of either unique or multiple groups of ER-phagy receptors (Fig. 1F).

**Fig. 1.**
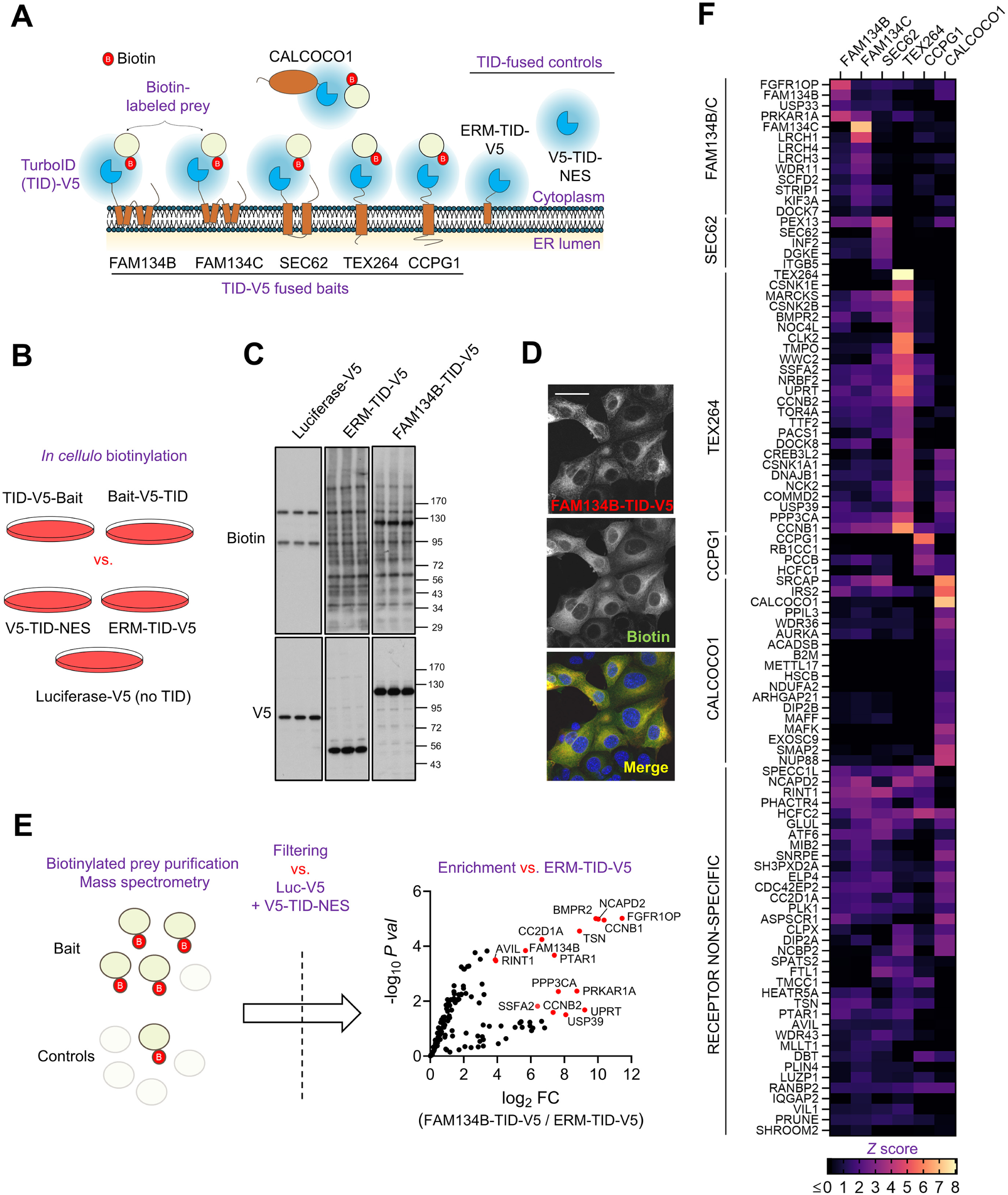
Proximity proteomics identifies novel cytoplasmic interactors of ER-phagy receptors. **(A-B)** ER-phagy receptor baits were fused with the biotin ligase TurboID (TID)-V5 tag at their cytoplasmic terminus(es) (schematic in **A** depicts one terminus each, for clarity). Stable expression in cells (see Fig. S1A) enables biotinylation of cytoplasmic proximal proteins (prey) upon cell supplementation with biotin. **(B)** Criterion for proximal protein status is enriched biotinylation in TID-V5-fused bait cells vs. control cells (ERM-TID-V5, anchored to the ER membrane; V5-TID-NES, cytoplasmic TID; or Luciferase-V5, no TID control). **(C)** Differential biotinylated protein abundance in PDAC cells after biotin treatment (0.5 mM) for 1 h (FAM134B-TID-V5 expressing cells) or 4 h (control cells), detected by HRP-Streptavidin blot (*n* = 3 replicates, see full dataset for all baits in Fig. S1B). MW markers, kDa. **(D)** Representative confocal immunofluorescence images of FAM134B-TID-V5 (V5) and biotinylation signal (Streptavidin-FITC) in PDAC cells treated as in (**C**) (representative of *n* = 2 independent experiments, see full dataset for all baits in Fig. S1C). Scale bar, 50 µm. **(E)** Biotinylated prey proteins obtained by streptavidin affinity-purification were detected by LC-MS/MS mass spectrometry, as described in Methods. Briefly, enrichment vs. controls was assessed by putting identified proteins through a cut-off filter using *P* values calculated vs. Luciferase-V5 and TID- NES (or ERM-TID for cytoplasmic CALCOCO1, see values in Methods). Remaining proteins were assessed for enrichment vs. ERM-TID (or TID-NES for CALCOCO1), as shown in the representative Volcano plot for an exemplar bait, FAM134B-TID-V5 (*n* = 3 replicates, FC = mean fold change, *P* val = *P* value, LIMMA analysis). The most enriched high-confidence hits are highlighted in red (log_2_ FC > 3.5 and -log_10_ *P* val > 1.3). Full Volcano plots for all baits in Fig. S1D. See also full dataset in Table S1. **(F)** Heatmap representation of the proximity prey hits using robust *Z* scores (see Methods). Proteins were manually clustered for apparent receptor specificity (data are combined for sets of replicates for N- and C-terminuses in this representation, see Methods and Table S1).

### cAMP-dependent control of FAM134B/C activity by PRKAR1A

To gain insight into integration of ER events with cytoplasmic physiology, we focused on FAM134B/C as prototypical ER-phagy receptor paralogues. We validated a CRISPR/Cas9 gRNA (short guide RNA) library targeting 21 FAM134B/C interactors (Fig. S2A) and screened for effects on flux of FAM134B/C, i.e., mobilisation within the ER and subsequent lysosomal degradation. This was performed using SRAI (signal-retaining autophagy indicator) ^59, 60^ reporter fusions of FAM134B/C in a semi-automated fluorescence microscopy assay (Fig. 2A). To maximise discovery, we measured effects on both basal flux and the flux induced by pleiotropic nutrient starvation (Earle’s Buffered Saline Solution, EBSS).

**Fig 2.**
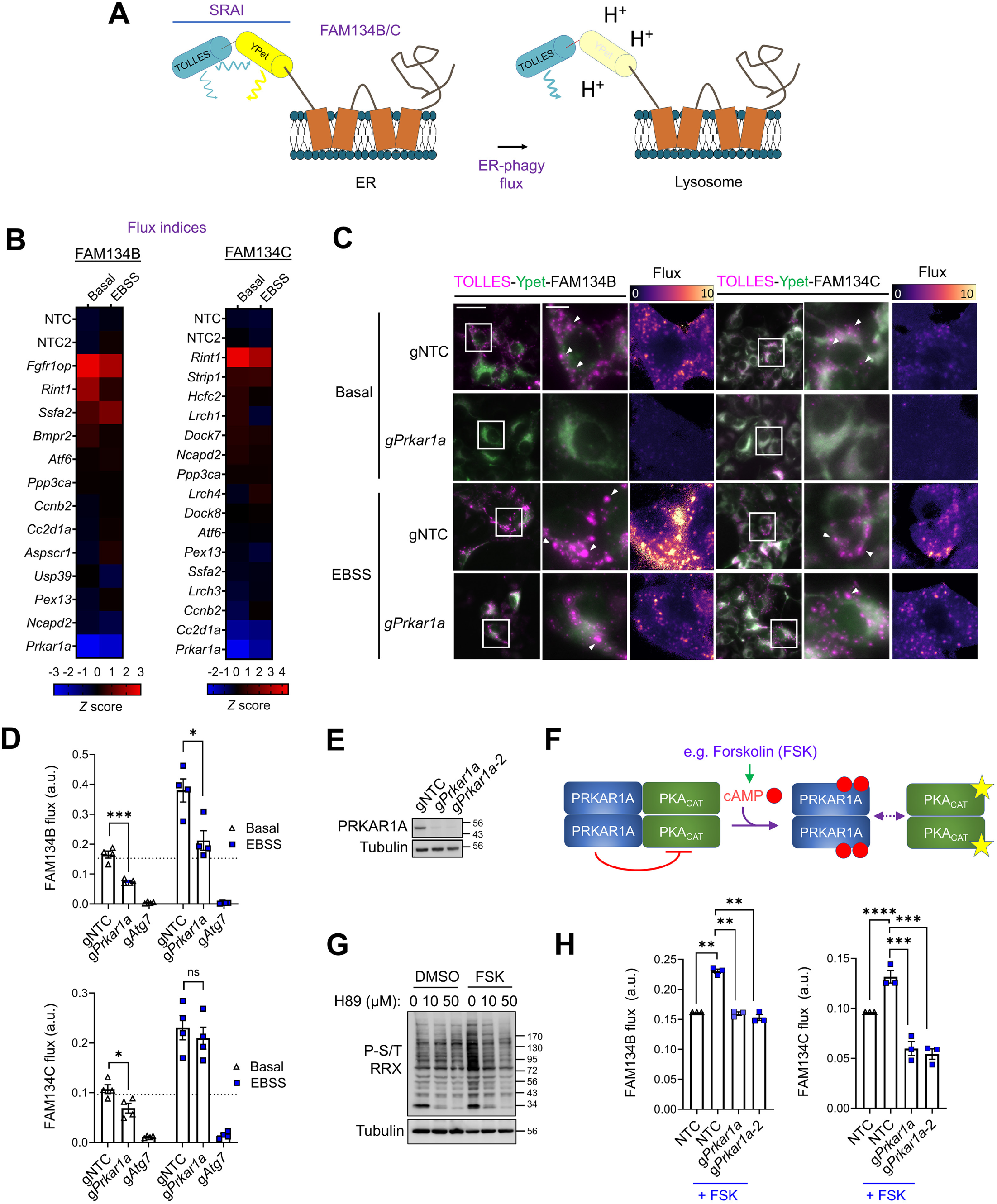
Identification of PRKAR1A as a cAMP-responsive activator of FAM134B/C. **(A)** Schematic of the Signal Retaining Autophagy Indicator reporter (SRAI, a tandem fusion of TOLLES and YPet fluorescent proteins) used to measure transit of fused FAM134B/C to the lysosome (ER-phagy flux). YPet is quenched in the acidic lysosome, concomitantly relieving Fluorescence Resonance Energy Transfer (FRET)-mediated suppression of TOLLES, eliciting a shift in fluorescence ratio. **(B)** Heatmap representing the distribution of fluorescence ratios (FAM134B/C receptor flux indices, see Methods) for PDAC-Cas9 cells stably-expressing SRAI fusions with FAM134B/C and transfected with the indicated gRNAs (short guide RNAs) for 6 days before culture under standard (Basal) or starvation (EBSS) conditions for 4 h (*n* = 4, two independent replicates each of N- and C-terminal SRAI fusions, *Z* score = robust *Z* scores of the means, 5 fields of 20-100 cells per replicate, see also Table S2). NTC = non-targeting control gRNA. **(C)** Widefield fluorescence images (TOLLES, magenta; YPet, green) and cognate fluorescence ratio (flux) heatmaps, derived from PDAC-Cas9 cells stably-expressing SRAI-FAM134B/C, transfected and treated in (**B**) (*n* = 4, representative images shown). Arrowheads indicate TOLLES-only foci (clusters of FAM134B/C that have undergone flux). White boxes delineate zoom panels. Scale bars, 50 µm (main), 10 µm (zoom). **(D)** Individual mean values of flux (TOLLES:YPet index) from secondary screen presented in (**B**), showing the values for gNTC (non-targeting control 1), g*Prkar1a* and additional g*Atg7* controls (targeting core autophagy, Fig. S2C) (*n* = 4, mean ± s.e.m., unpaired t-tests with Holm-Šídák, * = p ≤ 0.05, *** = p < 0.001, ns = p > 0.05). Dotted line indicates mean of gNTC2 (Basal). a.u. = arbitrary units. **(E)** Immunoblot of PRKAR1A levels in PDAC-Cas9 cells transfected with gNTC or 2 sequence unrelated *Prkar1a* gRNAs, g*Prkar1a* is the gRNA in panels (**B**) and (**C**) (*n* = 3 independent replicates, representative blot shown). **(F)** PRKAR1A dimers bind and inhibit PKA catalytic subunits (PKA_CAT_) but dissociate upon cAMP binding, relieving this inhibition (e.g., upon Forskolin [FSK] treatment). **(G)** Immunoblot for PKA-phosphorylated substrates (P-S/T RRX) in PDAC cells stimulated with DMSO vehicle or FSK (10 µM, 1 h), with or without additional PKA_CAT_ inhibitor H89 at indicated doses (*n* = 3 independent replicates, representative blot shown, see also Fig. S2D). **(H)** FAM134B/C flux assays performed with N-terminally fused SRAI-FAM134B/C as in (**B**), but with 2 sequence-unrelated g*Prkar1a* and with stimulation of cytoplasmic cAMP production via FSK treatment (10 µM for 2 h) (*n* = 3 independent replicates, ± s.e.m., gNTC FSK vs. DMSO by 1-sample two-tailed t-test, gNTC vs. *Prkar1a* gRNAs by 1-way ANOVA with indicated Holm-Šídák multiple comparisons, ** = *P* < 0.01, *** = *P* < 0.001, **** = *P* < 0.0001). a.u. = arbitrary units. MW markers, kDa.

Multiple genes (*Fgfr1op, Rint1, Ssfa2, Cc2d1a*) were found to encode regulators of FAM134B and/or FAM134C under basal and/or EBSS-stimulated conditions (Fig. 2B, Table S2). However, the strongest promoter of FAM134B and FAM134C flux was *Prkar1a* (Fig. 2B). Accordingly, we further validated PRKAR1A for proximity to FAM134B/C using low-throughput TID-mediated biotinylation and affinity purification (Fig. S2B). Notably, *Prkar1a* knockout substantially inhibited the *Atg7*-dependent basal flux of FAM134B/C (Figs. 2C-D, Fig. S2C). The knockout had a lesser effect on EBSS-induced flux, particularly for FAM134C (Figs. 2C-D). Overall, these data suggest the presence of signals for FAM134B/C activation that act through PRKAR1A, and that these signals are engaged to some extent under nutrient-replete, basal conditions.

The mechanism of PRKAR1A-mediated activation of FAM134B/C could hypothetically have been its canonical function of PKA_CAT_ suppression ^57^. However, we performed an SRAI reporter experiment employing two sequence-independent *Prkar1a* gRNAs (Fig. 2E) and forskolin (FSK) ^61^, which elevates intracellular cAMP and increases PRKAR1A dissociation from PKA_CAT_, activating the kinase activity of the latter (Figs. 2F-G, S2D). We observed that FSK provoked *Prkar1a*-dependent flux of FAM134B/C (Fig. 2H). As such, PRKAR1A-mediated suppression of PKA_CAT_ kinase was excluded as the activating mechanism for FAM134B/C activity. This is because, firstly, in the case of this rejected hypothesis, FSK- mediated activation of PKA_CAT_ (Fig. 2G) should lead to suppression of FAM134B/C activity; experimentally, the opposite effect was seen (Fig. 2H). Furthermore, if *Prkar1a* knockout-mediated PKA_CAT_ activation was responsible for stimulating FAM134B/C then FSK and *Prkar1a* knockout should have an additive effect in FAM134B/C activation or, alternatively, simply mask each other’s activating effect; experimentally, however, *Prkar1a* knockout opposes the activation of FAM134B/C by FSK (Fig. 2H). Thus, these data establish cAMP as the signal mediating FAM134B/C activity through PRKAR1A, likely via unknown, PKA_CAT_-suppression independent mechanism(s).

The above screening data, taken together, identify PRKAR1A as a cytoplasmic neighbour and kinase- independent, cAMP-stimulated activator of FAM134B/C. Given the ubiquity of cAMP in regulation of cellular state, this PRKAR1A-FAM134B/C association was prioritised for further molecular and functional inquiry.

### The cytoplasmic C-terminus of FAM134B binds a hydrophobic pocket in PRKAR1A

We next investigated the basis of the proximity interaction of FAM134B/C with PRKAR1A. Firstly, we found that ectopically-expressed FAM134B-V5 successfully co-immunoprecipitated (coIP’d) PRKAR1A (Fig. 3A), as also did endogenous protein (Figs. 3B, S3A) and FAM134C (Fig. S3B). Cross-linking using dithiobis-succinimidyl propionate (DSP) ^62^ was required to detect the FAM134B interaction robustly (Fig. 3A). Taking this data together, we inferred a transient or weak protein-protein interaction with PRKAR1A. Next, we tested mutants of PRKAR1A for coIP with FAM134B, in order to delineate the key domain(s) required for binding (Fig. 3C). In summary, the N-terminal 93 amino acids (NT) of PRKAR1A were found to be sufficient and necessary for interaction (Fig. 3D, NT and ΔNT mutants, respectively). Contrastingly, residues in the pseudo-substrate (PS) region that bind PKA_CAT_ were not required (mtPS mutant ^63^, Figs. 3C-D, S3C). Furthermore, AKAP11 — a LIR-containing protein that is a known binding partner of PRKAR1A NT ^64, 65^ — was also not required for PRKAR1A-FAM134B interaction (Fig. S3D-E). Thus, the NT domain of PRKAR1A binds FAM134B independently of PKA_CAT_ (and AKAP11), in line with the independence of FAM134B/C regulation from PKA_CAT_ suppression.

**Fig. 3.**
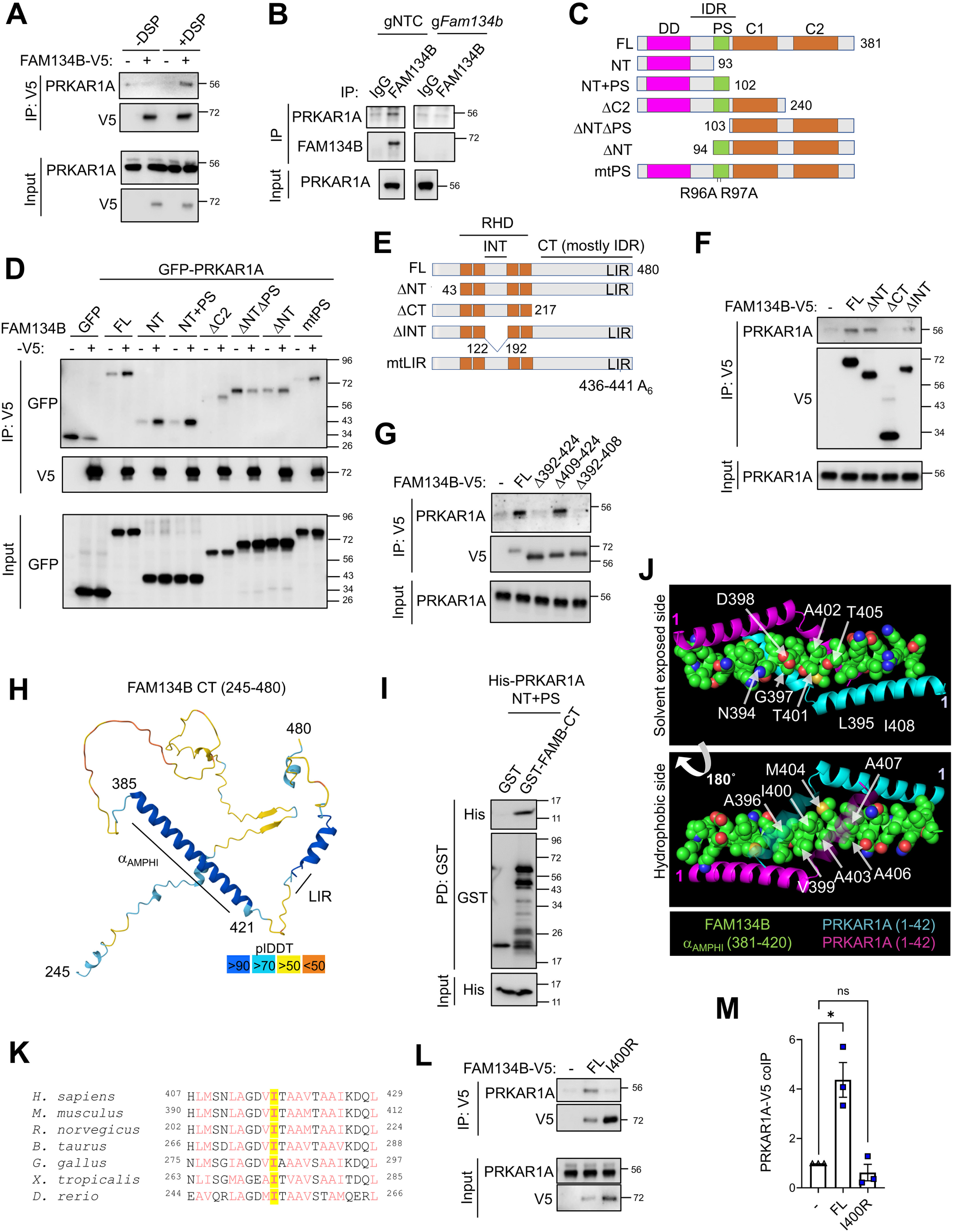
A hydrophobic pocket in PRKAR1A binds a helix within the C-terminus of FAM134B. **(A)** Immunoprecipitation (IP) of FAM134B-V5 from stably-expressing PDAC cells (+) or empty vector controls (-), with or without DSP-mediated crosslinking, IP and 1/50 input immunoblotted for endogenous PRKAR1A and V5 (*n* = 3 independent replicates, representative blot shown). **(B)** IP of endogenous FAM134B from PDAC-Cas9 cells transfected with NTC (non-targeting control) or *Fam134b* gRNA, IP and 1/50 input blotted for endogenous PRKAR1A and FAM134B (*n* = 4 independent replicates, representative blot shown, IgG = rabbit IgG). **(C)** Variants of PRKAR1A examined herein, compared with wild-type full-length (FL); NT = N-terminal region, PS = pseudo-substrate region, C1 and C2 = cAMP-binding domains, mtPS = PKA-binding deficient point mutant, IDR = intrinsically disordered region, DD = dimerization domain. **(D)** Cross-linking IP of FAM134B-V5 from stably-expressing PDAC cells (+) or empty vector controls (-), after transfection with GFP or N-terminally GFP-tagged PRKAR1A protein variants, IP and 1/50 input immunoblotted (*n* = 3 independent replicates, representative blot shown). **(E)** Variants of FAM134B examined herein (murine numbering), compared with wild-type full-length (FL); CT = C-terminal region, IDR = intrinsically disordered region, INT = internal cytoplasmic region, mtLIR = mutant LIR, NT = N-terminal region, RHD = reticulon homology domain). **(F-G)** Cross-linking IP of FAM134B-V5 from stably-reconstituted Δ*Fam134b/c* PDAC cells (FL vs. indicated point mutations or deletions) or empty vector control (-), IP and 1/50 input immunoblotted for endogenous PRKAR1A and V5 (*n* = 3 independent replicates, representative blot shown). **(H)** AlphaFold 3 prediction of the bulk of FAM134B CT (murine FAM134B amino acids [a.a.] 245-480). α_AMPHI_ = long amphipathic helix. LIR = LC3-interacting region. pIDDT = confidence score. **(I)** Immunoblot of *in vitro* glutathione-sepharose pull-down (PD) of GST or GST-FAM134B CT with bacterially-purified N-terminally His-tagged PRKAR1A NT+PS and 1/50 input (*n* = 2 independent replicates, representative blot shown). **(J)** Alphafold 3 prediction of FAM134B CT (murine a.a. 381-420) complexed with a dimer of PRKAR1A NT (murine PRKAR1A a.a. 1-42) (see also Fig. S3J), showing hydrophobic residues on FAM134B α_AMPHI_ in a groove generated by two helices from each monomer of PRKAR1A in antiparallel orientation. Red and blue atoms indicate oxygen and nitrogen within charged or otherwise polar residues. (K) Cross-species alignment of FAM134B α_AMPHI_ residues. Aliphatic residues in red. I400 (mouse numbering) highlighted in yellow. **(L-M)** Cross-linking IP of FAM134B-V5 from stably-reconstituted Δ*Fam134b/c* PDAC cells (FL vs. I400R mutant) or empty vector control (-), IP and 1/50 input immunoblotted for endogenous PRKAR1A and V5. Representative blot in (**L**), quantification in (**M**) for ratio of PRKAR1A IP to input, normalised to FAM134B-V5 IP (*n* = 3 independent replicates, mean ± s.e.m., normalised to empty vector control, 1- sample 2-tailed t-tests, * = *P* ≤ 0.05, ns = *P* > 0.05). MW markers = kDa.

Within FAM134B/C, candidate cytoplasmic interaction sites for PRKAR1A included the predominantly intrinsically-disordered C-terminal domain (CT), the N-terminal region (NT) and the region between the membrane-embedded hairpins of the RHD (INT) (Fig. S3F). To identify the site, we reconstituted Δ*Fam134b/c* double-knockout (DKO) cells (Fig. S3G) with FAM13B-V5, either wild-type (full-length, FL) or mutant (Fig. 3E), and performed coIP for PRKAR1A. These experiments revealed that the LIR in FAM134B was dispensable for PRKAR1A binding (Fig. S3H), as were the NT and INT regions (Fig. 3F). However, the CT region was required for binding (ΔCT mutant, Fig. 3F). Binding was further mapped to within amino acids 392-408 of FAM134B (Figs. 3G, S3I, murine numbering). Strikingly, within the predominantly disordered C-terminal domain, this sequence of amino acids stood out as having predicted amphipathic α-helix (α_AMPHI_) forming capacity (amino acids 385 to 421, Fig. 3H). Two shorter amphipathic helices have previously been reported adjacent to the hairpins and to interact with phospholipid membrane ^31, 38, 66^. A short amphiphilic helix adjacent to the LIR also promotes ATG8 binding ^67^ (Fig. 3H). However, the longer α_AMPHI_ has not been characterised. We noted that the energetic favourability of α_AMPHI_ was likely to be diminished in aqueous solution of the monomer, implying a potential hydrophobic interaction with PRKAR1A. Furthermore, we concluded that such an interaction would be direct in nature, as bacterially-expressed, N-terminal regions of PRKAR1A (NT+PS) bound to recombinant FAM134B CT *in vitro* (Fig. 3I). In line with a direct hydrophobic interaction, modelling of the PRKAR1A NT and FAM134B CT complex showed that the hydrophobic face of α_AMPHI_ could be shielded within a pocket created by two subunits of the dimerization domain (DD) subregion of PRKAR1A NT (Figs. 3J, S3J). Cross-species conservation of aliphatic residues in this region (Fig. 3K) aided us in identifying an experimental point mutation within FAM134B α_AMPHI_, I400R (I417 in human FAM134B), which was predicted to disrupt the trimeric complex (FoldX analysis of mutational difference in the free energy change [ΔΔ*G*] = 1.69 for FAM134B CT-PRKAR1A NT trimeric structure; ΔΔG = -0.19 for FAM134B CT secondary structure, including α_AMPHI_). This was confirmed by coIP (Fig. 3L-M), both corroborating the structural interaction between FAM134B and PRKAR1A, and generating a point mutation for further interrogation of the role of this interaction.

In summary, a helix within the predominantly disordered C-terminal domain of FAM134B binds directly to PRKAR1A, independently of LC3B, revealing a molecular connection between the ER membrane and a cytoplasmic, signal-responsive scaffold protein.

#### Liquid-like condensates of PRKAR1A associate with FAM134B clusters at ER contacts

Given the PKA_CAT_ independence of FAM134B/C activation in response to cAMP, we hypothesised instead a causal role for the cAMP-responsive liquid-like condensation property of PRKAR1A ^54, 55^. In line with this, FSK treatment induced cytoplasmic foci of PRKAR1A that exhibited colocalization with mApple-FAM134B and LC3B (Fig. 4A-B). Endogenous FAM134B and tagged FAM134C similarly formed foci with PRKAR1A (Fig. S4A-B). To test whether these PRKAR1A-colocalised structures were truly condensates, we used cells co-expressing PRKAR1A and FAM134B fused with mApple and EGFP fluorescent proteins, respectively. After FSK treatment to induce foci positive for both fluorophores, we performed two series of FRAP (fluorescence recovery after photobleaching) experiments ^68^— one for each fluorescent protein resident at the foci (Fig. 4C-D). Notably, focal fluorescence of the PRKAR1A-fused mApple recovered rapidly after bleaching, similarly to diffuse cytosolic mApple (Fig. 4C-D, Movie S1). Contrastingly, FAM134B-fused EGFP showed very limited recovery from bleaching (Fig. 4C-D, Movie S2). Thus, at sites of cAMP-induced FAM134B colocalization, PRKAR1A displays liquid-like properties, whereas the FAM134B within the focus is immobilised, consistent with embedment within ER membrane and clustering therein. In line with this, PRKAR1A-FAM134B foci were detected at the ER network – marked by RTN4 – using confocal light microscopy (Figs. S4C-D). Next, we set out to obtain greater insight into the spatial relationship between the FAM134B-positive ER and PRKAR1A condensates using super-resolution and ultrastructural analyses. Firstly, imaging by SoRA (super resolution imaging via optical photon reassignment) revealed cAMP-induced PRKAR1A foci to be irregularly-shaped globules that were partially enwrapped by layers of FAM134B and LC3B, which themselves showed substantial but incomplete overlap (Figs. 4E, S4E, Movie S3). Furthermore, correlative-light electron microscopy (CLEM) of PRKAR1A and FAM134B double-positive foci revealed electron-dense, irregularly-shaped structures that exhibited extensive interorganellar contacts with adjacent ER membrane (Figs. 4F, S4F). ER tubules appeared to delimit these structures in part but non-membrane delimited regions of the periphery in contact with the surrounding cytoplasm were readily apparent (Figs. 4F, S4F). We thus interpreted these structures as PRKAR1A condensates forming extensive interorganellar contacts with FAM134B-enriched regions of ER. Furthermore, partially- embedded membrane-delimited vesicles, most prominently lysosomal structures, were identified at the periphery of these condensates (Figs. 4F, S4F). This observation was corroborated by SoRA imaging of LAMP1-positive lysosomes within PRKAR1A foci (Fig. 4G, Movie S4).

**Fig. 4.**
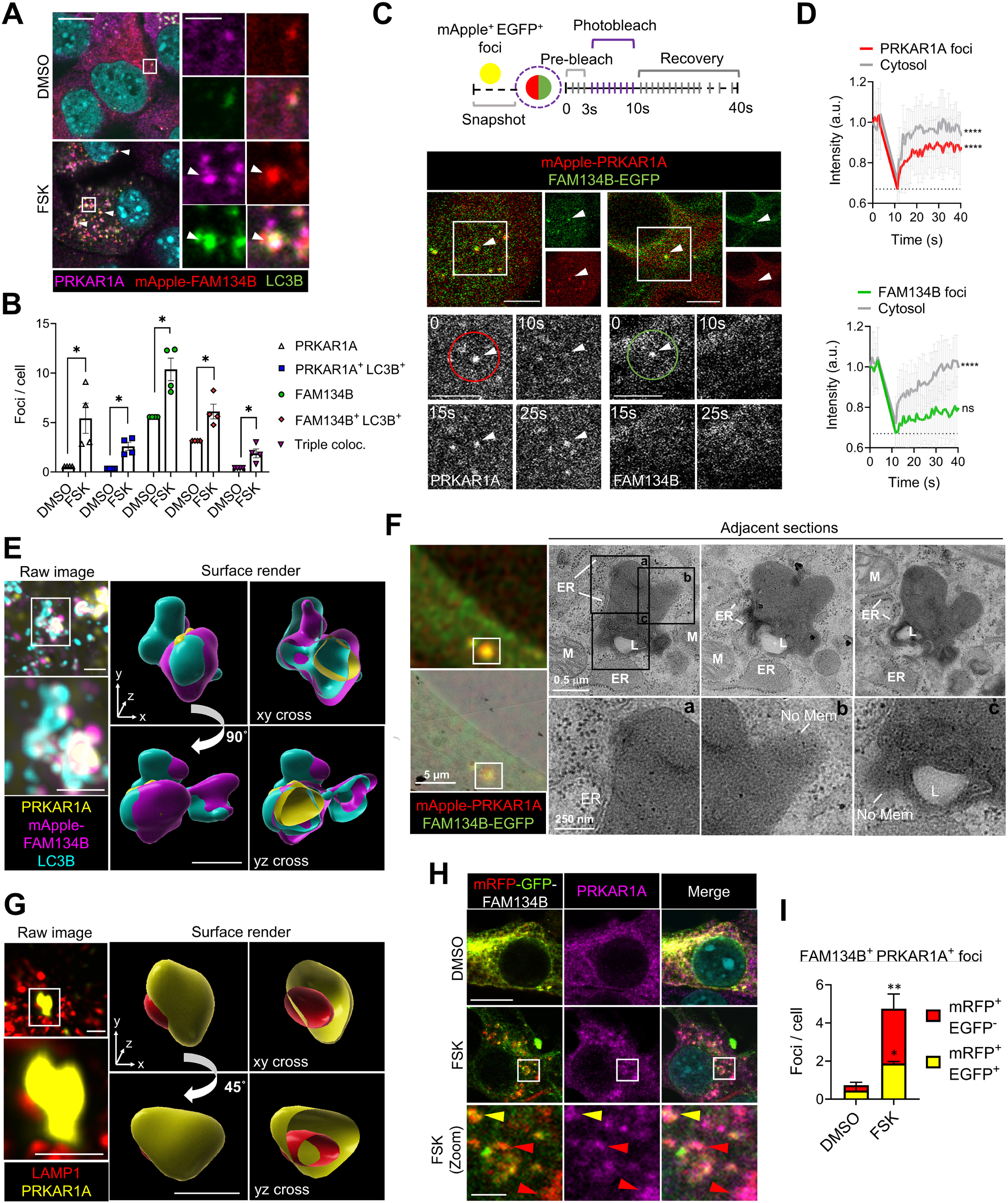
PRKAR1A liquid-like condensates form ER contacts at sites of FAM134B/C clustering. **(A-B)** Confocal immunofluorescence microscopy of PDAC cells stably expressing mApple-FAM134B (red) treated with DMSO or forskolin, FSK (40 µM, 4 h), co-stained for LC3B (green) and PRKAR1A (magenta). Representative images in (**A**), white boxes delineate zoom panels. Scale bars, 10 µm (main), 2 µm (zoom). Arrowheads indicate triple colocalization. Quantified in (**B**) for indicated focus abundance (*n* = 4 independent experiments, > 100 cells/condition, mean ± s.e.m., DMSO normalised to mean value, 1-sample two-tailed t-tests, DMSO vs. FSK, * = *P* ≤ 0.05). **(C-D)** FRAP of mApple-PRKAR1A and FAM134B-EGFP co-localising foci in FSK-treated PDAC cells (40 µM, 2 h). Two experiments were performed, one for each fluorophore. Schematic and representative images in (**C**) illustrate experimental design and typical results. See also Movies S1-2. Arrowheads in colour image indicate dual colour foci prior to bleaching, white boxes delineate zoom images. Zooms indicate photobleaching and recovery at time points corresponding to the schematic (bleached regions-of-interest (ROIs) delineated by circles). Bleached foci are indicated by white arrows. Cytoplasmic areas with no foci were bleached as controls. Quantifications of normalised fluorescence intensity over time are shown in (**D**) (*n* = 23 [mApple] or 21 [EGFP] bleached foci ROI; *n* = 14 [mApple] or 16 [EGFP], bleached no-foci cytoplasm ROI, ± s.d., 1-sample t-test of area under curves from 10 s to 20 s vs. basal [dotted line], **** = p < 0.0001, ns = p > 0.05). a.u. = arbitrary units. Scale bars, 5 µm. **(E)** SoRa super-resolution immunofluorescence microscopy analysis of PDAC cells stably-expressing mApple-FAM134B (magenta) treated with FSK (40 µM, 4 h), co-stained for LC3B (cyan) and PRKAR1A (yellow). See also Movie S3. Left panel, representative raw image and zoom (delineated by white box). Right panel, representative Imaris surface rendering of the zoomed image and cross-sections thereof (*n* = 2 independent replicates, see also volume rendered example in Fig. S4E). Scale bars, 1 μm. **(F)** Correlative light electron microscopy (CLEM) of mApple-PRKAR1A and FAM134B-EGFP co-localised foci after FSK treatment (40 µM, 2 h) (see also example in Fig. S4F). Left panels, representative fluorescence overlay. Right panels, representative electron microscopy (EM). Boxes indicate zoom panels. Scale bars, 5 μm (fluorescence), 0.5 μm (EM) and 250 nm (EM zoom). ER = endoplasmic reticulum; L = lysosome; M = mitochondrion; No mem = non-membrane delimited. **(G)** SoRa super-resolution immunofluorescence microscopy of PDAC cells treated with FSK (40 µM, 30 min), co-stained for LAMP1 (red) and PRKAR1A (yellow). See also Movie S4. Left panel, representative raw image and zoom (delineated by white box). Right panel, representative Imaris surface rendering of the zoomed image and cross-sections thereof (*n* = 2 independent replicates). Scale bars, 1 μm. **(H-I)** Fluorescence confocal microscopy of PDAC cells stably-expressing mRFP-GFP-FAM134B treated with DMSO or FSK (40 µm, 30 min), co-stained for PRKAR1A (magenta). Representative images in (**H**), white boxes indicate zoom panels, and quantification in (**I**) of lysosomal/acidified FAM134B^+^ PRKAR1A^+^ foci (red arrowheads in (**H**), mRFP^+^ GFP^-^) and non-lysosomal FAM134B^+^ PRKAR1A^+^ foci (yellow arrowheads in (**H**), mRFP^+^ GFP^+^) (*n* = 3 independent replicates, > 100 cells/condition, mean ± s.e.m., 2- way ANOVA and Holm-Šídák multiple comparisons, DMSO vs. FSK, * = *P* ≤ 0.05, ** = *P* ≤ 0.01). Scale bars, 10 μm (main), 2 µm (zoom).

Finally, in line with the incorporation of lysosomes, immunofluorescence analysis of tandem-tagged flux reporters – mRFP-GFP fused to FAM134B ^6^ and co-stained with PRKAR1A – demonstrated that cAMP-induced FAM134B-PRKAR1A foci were indeed sites where FAM134B traffic into lysosomes occurred (Fig. 4H-I).

Overall, the above data show that cAMP-mediated liquid-like condensates of PRKAR1A interact with clustered FAM134B/C at condensate-ER contact sites and that this correlates spatially with FAM134B flux, framing these novel structures as degradation hubs.

### Liquid-like condensates facilitate FAM134B clustering and interaction with LC3B

We next hypothesised that PRKAR1A liquid-like condensation was causal for FAM134B/C-LC3B clustering, thus facilitating flux. In line with this hypothesis, we first blocked PRKAR1A condensate formation in response to FSK by treatment with the solvent 1,6-hexanediol (Hex) ^69^, and found that this also ablated formation of FAM134B-LC3B foci (Fig. 5A-B). In contrast, the PKA_CAT_ inhibitor H89 ^61^ (Fig. 2G) did not greatly impact either of these phenomena (Fig. 5A-B). These findings were corroborated by genetic observation. First, the formation of FAM134B-LC3B foci in response to FSK was ablated in *Prkar1a* knockout cells (Fig. 5C-D). However, this was rescued by reconstitution with wild-type, condensate-forming PRKAR1A (full-length, FL) but not by condensate-deficient PRKAR1A ΔIDR ^54^ (Figs. 5E-G, S5A-B), despite this mutant retaining intrinsic ability to bind to FAM134B (Fig. S5C). Notably, PKA_CAT_-binding deficient PRKAR1A (mtPS) also successfully formed condensates and rescued FAM134B-LC3B foci (Figs. 5E-G, S5A-B). These genetic observations thus consolidated the hypothesis that liquid-like condensation of PRKAR1A – and not PKA_CAT_ interaction – was required for FAM134B- LC3B clustering.

**Fig. 5.**
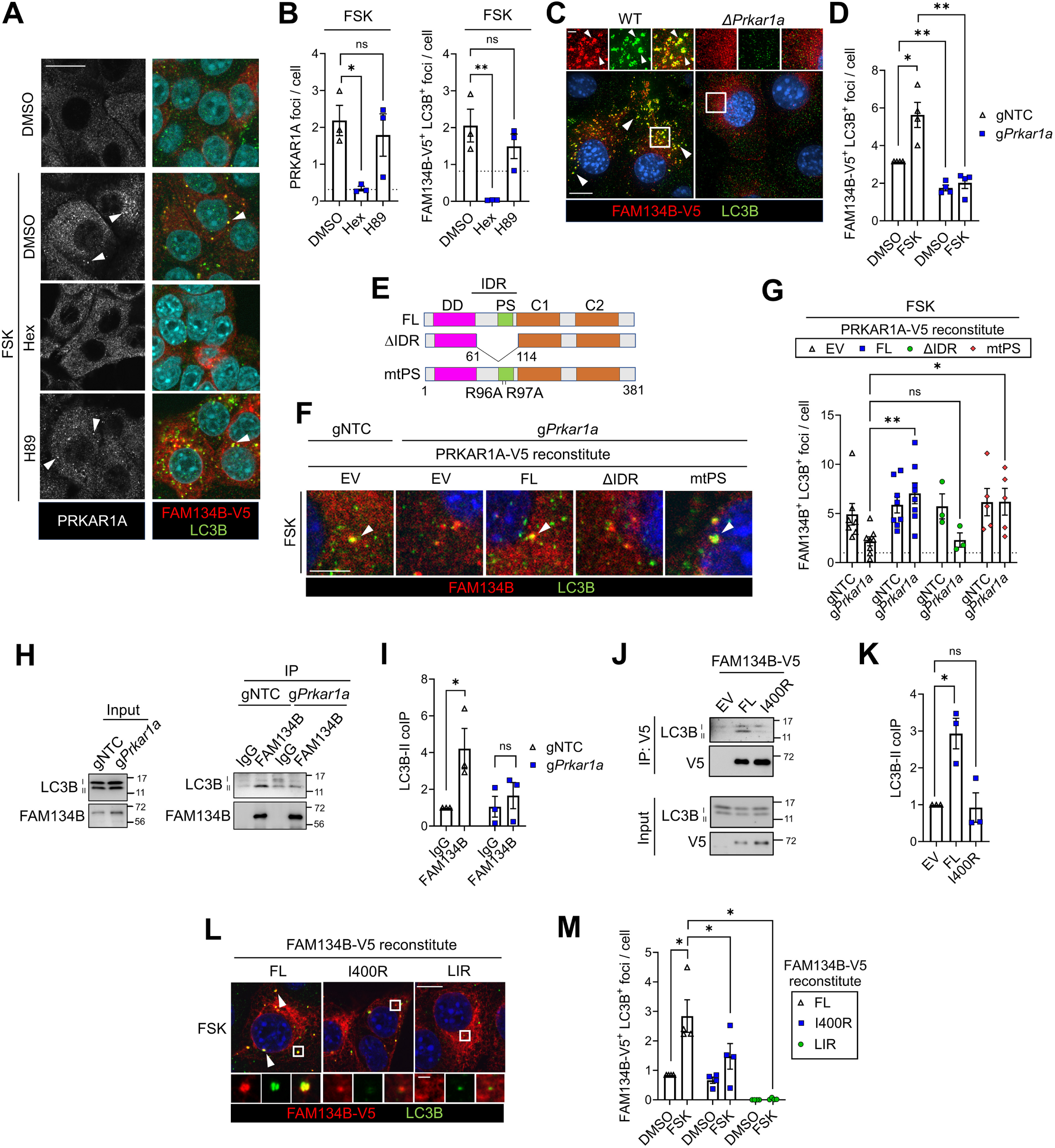
Liquid-like condensates cluster FAM134B enabling interaction with LC3B. **(A-B)** Confocal immunofluorescence microscopy of PDAC cells stably-expressing FAM134B-V5 and treated with DMSO or forskolin, FSK (40 μM, 2 h), +/- 1,6-hexanediol (Hex, 2.5 %) or PKA_CAT_ inhibitor H89 (20 μM), stained with PRKAR1A or co-stained for V5 and LC3B. Representative confocal channel images in (**A**) of PRKAR1A (white, left panels) or V5 and LC3B (red and green, right panels). White arrowheads indicate PRKAR1A foci (left panels) or FAM134B-V5^+^ LC3B^+^ foci (right panels). Scale bar, 20 μm. Quantifications presented in (**B**) for PRKAR1A foci and FAM134B-V5^+^ LC3B^+^ foci in FSK-treated cells, normalised to no FSK control (dotted line) (*n* = 3 independent replicates, > 100 cells/condition, mean ± s.e.m., 1-way ANOVA and Holm-Šídák multiple comparisons, ** = *P* < 0.01, ns = *P* > 0.05). **(C)** SoRa super-resolution immunofluorescence microscopy of PDAC WT or *ΔPrkar1a* cells stably-expressing FAM134B-V5 and treated with FSK (40 µM, 4 h), co-stained for V5 (red) and LC3B (green). Zoom panels are delineated by white boxes. Arrowheads indicate FAM134B-V5^+^ LC3B^+^ foci. (*n* = 2 independent replicates, representative image). Scale bars, 10 μm (main), 2 µm (zoom). **(D)** Quantification of FAM134B-V5^+^ LC3B^+^ foci from confocal immunofluorescence microscopy of PDAC-Cas9 cells stably-expressing FAM134B-V5 and transfected with indicated gRNAs (NTC = non-targeting control) and treated with DMSO or FSK (40 µM, 4 h), co-stained for V5 and LC3B (*n* = 4 independent replicates, > 100 cells/condition, ± s.e.m., normalised to mean of gNTC + DMSO, 1-sample 2-tailed t-tests vs. gNTC + DMSO, 2-tailed t-test between FSK treated, * = *P* ≤ 0.05, ** = *P* < 0.01). **(E)** Variants of PRKAR1A examined in (**F**-**G**), compared with wild-type, full-length (FL); mtPS = PKA-binding deficient point mutant, ΔIDR = non-condensing intrinsically disordered region deletion. **(F-G)** PDAC-Cas9 were transduced with gRNA-insensitive PRKAR1A-V5 variant (or empty vector control, EV) then transfected for 7 days with either NTC gRNA or gRNA for endogenous *Prkar1a*. Cells were then treated with DMSO or FSK (40 µM, 4 h). Representative confocal immunofluorescence images of FSK-treated cells shown in (**F**), co-stained for endogenous FAM134B (red), LC3B (green) and V5 (see Fig. S5A). Arrowheads indicate co-localising foci. Scale bar, 5 μm. Quantification in (**G**) of co- localising FAM134B^+^ LC3B^+^ foci in FSK-treated, V5-positive cells. Numbers normalised to basal (dotted line, gNTC + DMSO + EV reconstituted) (*n* ≥ 3 independent replicates, >100 cells/condition, mean ± s.e.m., 2-way ANOVA and Holm-Šídák multiple comparisons, ** = *P* < 0.01, * = *P* ≤ 0.05, ns = *P* > 0.05). **(H-I)** Immunoprecipitation (IP) of endogenous FAM134B from PDAC-Cas9 cells transfected with indicated NTC or *Prkar1a* gRNA, IP and 1/50 input immunoblotted for endogenous LC3B. Representative blot shown in (**H**), quantification of LC3B-II (lipidated LC3B) ratio in IP to input, normalised to control (gNTC with IgG) in (**I**) (*n* = 3 independent replicates, ± s.e.m., 1-sample 2-tailed t-test for gNTC, 2-tailed student t-test for g*Prkar1a*, * = *P* ≤ 0.05, ns = *P* > 0.05). **(J-K)** IP of FAM134B-V5 from stably-reconstituted PDAC Δ*Fam134b/c* cells (wild-type, full-length [FL] vs. I400R), IP and 1/50 input immunoblotted for LC3B. Representative blot in (**J**), quantification in (**K**) of ratio of LC3B-II (lipidated LC3B) in IP to input, normalised to EV (empty vector, non-reconstituted) control (*n* = 3 independent replicates, ± s.e.m., 1-sample two-tailed t-tests, * = *P* ≤ 0.05, ns = *P* > 0.05). **(L-M)** Stably-reconstituted PDAC Δ*Fam134b/c* cells (FL, I400R or LIR mutant) treated with DMSO or FSK (40 µM, 4 h), co-stained with V5 (red) and LC3B (green). (**L**) Confocal images of FSK-treated cells. Arrowheads, FAM134B-V5^+^ LC3B^+^ co-localising foci. White boxes, zooms. Scale bars, 10 µm (main), 1 µm (zoom). (**M**) FAM134B-V5^+^ LC3B^+^ foci numbers, normalised to FL + DMSO (*n* = 4, > 100 cells/condition, ± s.e.m., 1-sample t-test FSK FL vs. DMSO FL, or Holm-Šídák t-tests, * = p ≤ 0.05).

Biochemical data also underscored that FAM134B/C binding with LC3B was dependent upon PRKAR1A interaction. Firstly, coIP of LC3B with FAM134B or FAM134C was ablated by *Prkar1a* knockout (Figs. 5H-I, S5D-E). Secondly, whereas FL FAM134B protein binds LC3B upon reconstitution of Δ*Fam134b/c* DKO cells (Fig. 5J-K), correlating with formation of FAM134B-LC3B foci (Fig. 5L-M), the I400R PRKAR1A binding-deficient mutant is defective in both LC3B binding and co-localisation (Fig. 5J-M).

The above data, taken together, demonstrate that cAMP-stimulated condensates of PRKAR1A are required to promote clustering and association of FAM134B/C with LC3B.

### FAM134B at liquid-like condensates sequesters and inhibits RhoA

An important function of condensates is spatiotemporal co-ordination of cytoplasmic signalling ^44^. Here, we have shown that condensates act upstream of ER-phagy at ER contact sites, clustering FAM134B/C proteins in response to cAMP and leading to their degradation. However, it remains unresolved whether FAM134B/C clustered in this manner can exert downstream cytoplasmic function in co-ordinating responses to cAMP. We thus performed a proximity proteomics-led investigation of FAM134B cytoplasmic interactors after FSK treatment (Figs. 6A, S6A-C, Table S3). Reactome analysis^70^ of the interactome revealed that the most enriched signatures upon FSK treatment were of signalling by Rho-family small GTPases, most prominently RhoA (Figs. 6B, S6D). Indeed, the dataset revealed proximity of FAM134B with a wide network of RhoA interacting proteins, including known upstream regulators and downstream effectors (Figs. 6A, S6E-F). While RhoA levels in the proteomic dataset were not confidently quantifiable, it was nonetheless striking that RhoA was the common physical and functional interactor of this most significant category of FAM134B proximal proteins.

**Fig. 6.**
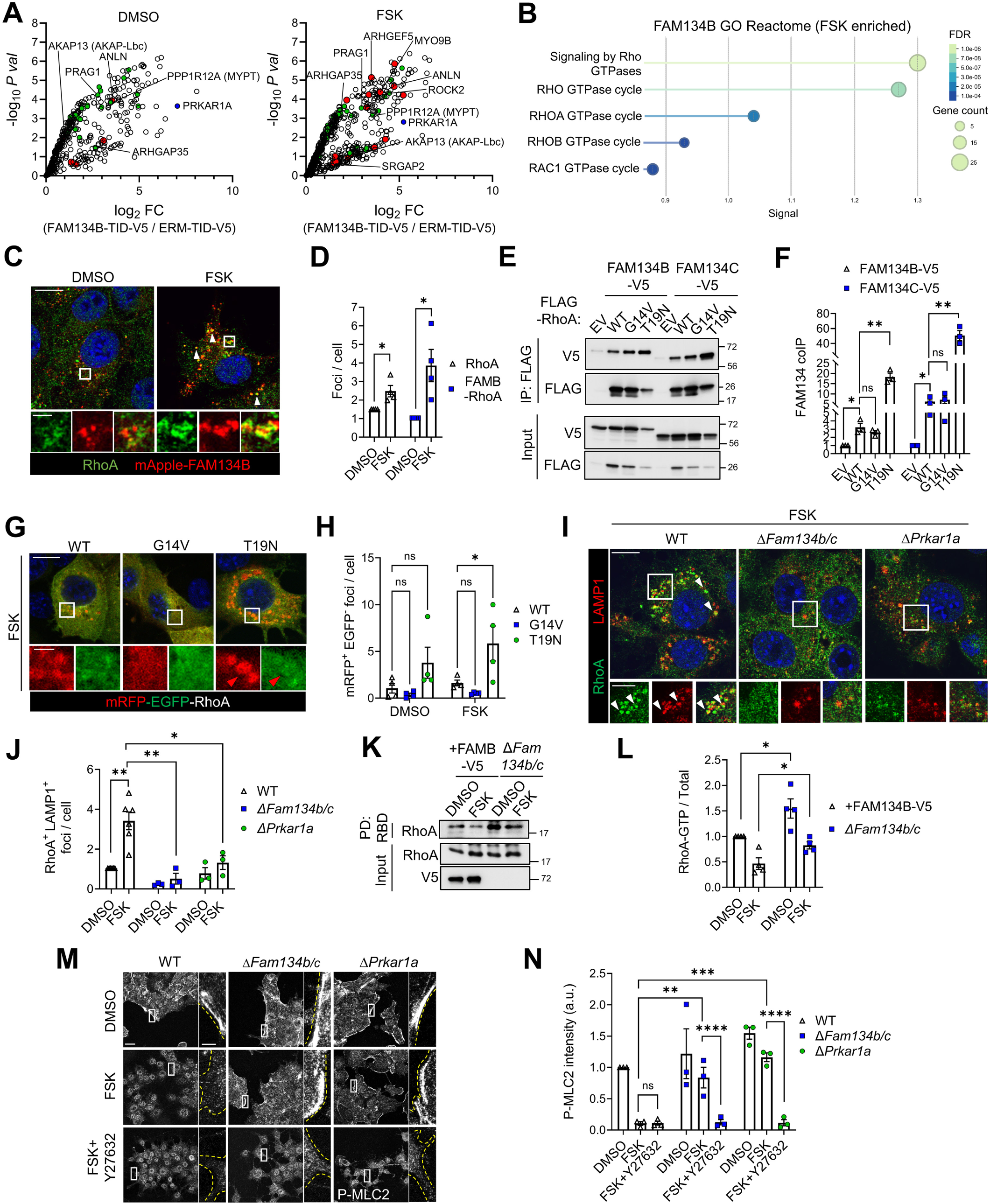
FAM134B at condensates sequesters and inhibits cytoplasmic RhoA. **(A)** Volcano plots for enriched biotinylated proteins after proximity proteomics for FAM134B-TID-V5. Shown are proteins remaining after *P*-value cut-off versus TID-NES controls, plotted for enrichment vs. ERM-TID (*n* = 3 independent replicates, FC = mean fold change, *P* val = *P* value, LIMMA analysis). See also Table S3. Red highlights FSK-enriched prey proteins from within enriched Reactomes (Fig. 6B). Green highlights additional proteins from within these Reactomes present in analysis of all conditions (+/- FSK, Fig. S6E). Example labels are RhoGEFs, RhoGAPs, and RhoA effectors/downstream substrates. **(B)** Reactome Pathway enrichment analysis of prey hits enriched in FSK-stimulated FAM134B-TID-V5 proximity proteomics versus DMSO see also Table S3 and Fig. S6A-F). FDR = false discovery rate. **(C-D)** Immunofluorescence confocal microscopy of PDAC cells stably-expressing mApple-FAM134B (red) treated with DMSO or FSK (40 µM, 4 h), co-stained for RhoA (green). Representative image in (**C**). White boxes indicate zooms. Scale bars, 10 μm (main), 2 µm (zoom). Arrowheads indicate FAM134B^+^ RhoA^+^ co-localising foci. Quantifications in (**D**), RhoA foci and co-localising foci, normalised to DMSO (*n* = 4, > 100 cells/condition, mean ± s.e.m., 1-sample 2-tailed t-tests, * = *P* ≤ 0.05). **(E-F)** FLAG co-IP (immunoprecipitation) from HEK293FT cells co-transfected with FAM134B/C-V5 and FLAG-RhoA variants (WT, wild-type; G14V, active; T19N, inactive), or empty vector control (EV) and input 1/50. Representative immunoblot shown in (**E**), quantified as ratio of IP to input FAM134B/C- V5, normalised to IP FLAG-RhoA, in (**F**) (*n* = 3, mean ± s.e.m., normalised to EV, 1-sample two-tailed t-test WT vs. EV and Holm-Šídák corrected t-tests, ** = *P* < 0.01, * = *P* ≤ 0.05, ns = *P* > 0.05). **(G-H)** Confocal fluorescence microscopy of PDAC expressing mRFP-EGFP-RhoA variants. Representative FSK-treated cells in (**G**). White boxes, zoom panels. Scale bars, 10 μm (main), 2 µm (zoom). (**H**) Quantifications of lysosomal (mRFP^+^ EGFP^-^) foci (arrowheads) (*n* = 4, >100 cells/condition, mean ± s.e.m., 2 way-ANOVA with Holm-Šídák multiple comparisons, * = *P* ≤ 0.05, ns = *P* > 0.05). **(I-J)** Confocal immunofluorescence microscopy of PDAC parental (WT, wild-type) or indicated knockout cells (Δ) treated with DMSO or FSK (40 µM, 4 h), co-stained for RhoA (green) and LAMP1 (red). Representative images of FSK-treated cells in (**I**). White boxes, zooms. Arrowheads indicate co- localising foci. Scale bars, 10 μm (main), 5 µm (zoom). Quantification of colocalising foci in (**J**), normalised to mean of DMSO + WT (*n* ≥ 3, > 100 cells/condition, mean ± s.e.m., 1-sample, 2-tailed t-test for DMSO vs. FSK in WT, and Holm-Šídák multiple t-tests, ** = *P* < 0.01, * = *P* ≤ 0.05). **(K-L)** Analysis of active RhoA in PDAC Δ*Fam134b/c* cells (right lanes) or reconstituted (rescue) cells expressing FAM134B-V5 (left lanes) after treatment with DMSO or FSK (40 µM, 4 h), using Rhotekin-Rho-binding domain (RBD) pull-down (PD) for RhoA-GTP (see reconstitution control, Fig. S6I). Representative blot in (**K**), quantified in (**L**) for ratio of RhoA PD to input, normalised to Reconstituted + DMSO (*n* = 4, mean ± s.e.m., 2-way ANOVA with Holm-Šídák multiple comparisons, * = *P* ≤ 0.05). **(M-N)** Confocal microscopy of PDAC (WT, wild-type) or indicated knockouts (Δ) treated with DMSO or FSK (40 µM, 2 h) (+/- 30 min pre-treatment ROCK inhibitor [Y27632], 20 µM), stained for phospho- MLC2. Representative images in (**M**). Yellow lines, periphery of cells. Scale bars, 20 μm (main), 5 µm (zoom). (**N**) Quantifications of peripheral extranuclear P-MLC2, normalised to WT + DMSO, a.u. = arbitrary units (*n* = 3, > 200 cells/condition, mean ± s.e.m., 2-way ANOVA and Holm-Šídák comparisons, **** = *P* < 0.0001, *** = *P* < 0.001, ** = *P* < 0.01, ns = *P* > 0.05).

RhoA GTPase is an enzyme recruited to the plasma membrane from the cytoplasm during activation^71^. In the active, GTP-bound form, membrane-localised RhoA recruits downstream effectors, e.g., the ROCK1/2 kinases, hereafter ROCK (notably, ROCK2 is also detected in the FSK-induced FAM134B proximity proteome, Figs. 6A, S6F, Table S3). ROCK drives cell peripheral phosphorylation of myosin light chain (phospho-MLC2, P-MLC2) ^72^.

Stimulated by these observations, we investigated the relationship of RhoA with condensate-associated FAM134B/C. First, we found that endogenous RhoA colocalises with FAM134B and PRKAR1A foci, dependent upon FSK (Figs. 6C-D, S6G-H). Thus, RhoA is concentrated at ER-adjacent condensates. Secondly, coIP demonstrated that FAM134B/C could bind RhoA – notably with strong selectivity for the inactive over the active form (T19N GTP-binding deficient mutant versus WT or G14V, a constitutively GTP-bound mutant ^73^) (Fig. 6E-F). Consistent with this binding selectivity, tandem tagging of RhoA with mRFP-GFP demonstrated that the T19N form is selectively trafficked to lysosomes in an FSK-dependent manner (Fig. 6G-H). Taking the above data together, we conclude that cAMP mediates sequestration of RhoA in lysosomes, dependent upon binding with FAM134B/C at PRKAR1A condensate contact sites. Corroborating this, we observed that FSK-induced endogenous RhoA recruitment to lysosomal foci (LAMP1) was dependent upon both *Fam134b/c* and *Prkar1a* (Fig. 6I-J).

Finally, we addressed whether inhibition of RhoA signalling, a known output of cAMP ^57^, was dependent upon FAM134B/C and PRKAR1A. Firstly, RhoA-GTP pull-down assays ^74^ demonstrated that FSK-stimulated suppression of RhoA-GTP levels was indeed mitigated in Δ*Fam134b/c* DKO cells (Figs. 6K-L, S6I). Secondly, FSK-mediated abrogation of peripheral P-MLC2 – a complementary measure of RhoA-ROCK output – was dependent upon *Fam134b/c* and *Prkar1a* (Fig. 6M-N). Furthermore, direct RhoA-ROCK pathway inhibition by treatment with the ROCK kinase inhibitor Y27632 ^75^ compensated for *Fam134b/c* and *Prkar1a* deficiency (Fig. 6M-N). Contrastingly, H89 treatment did not reverse FSK-mediated suppression of P-MLC2, further underscoring the PKA_CAT_ activity-independent role of PRKAR1A condensates in co-ordinating FAM134B/C function (Fig. S6J-K).

These data show that PRKAR1A condensates facilitate sequestration of cytoplasmic RhoA through interaction of the latter with clustered FAM134B/C, resulting in lysosomal targeting and inhibition of downstream signalling.

### Physiological function of the PRKAR1A-FAM134B/C axis in cell morphology and invasion mode

The preceding data show that cytoplasmic condensates of PRKAR1A contact the ER and bind FAM134B/C, thus co-ordinating ER-phagy receptor and RhoA sequestration into lysosomes. However, the physiological role of this remains unclear. Notably, a readily apparent effect of cAMP – associated with suppression of RhoA and P-MLC2 – is reduction in peripheral contractile filamentous actin (F-actin), in turn driving morphological events such as formation of lamellipodia or tubulin-rich, neurite-like extensions ^56, 76^. Indeed, tubulin-rich protrusions were observed in PDAC cells upon FSK treatment; strikingly, this was dependent upon *Prkar1a* and *Fam134b/c*, firmly implicating ER-condensate contact sites in this morphological response to cAMP (Fig. 7A-B). Furthermore, reconstitution of Δ*Fam134b/c* DKO cells with FAM134B mutants demonstrated that FSK-driven morphological changes were dependent upon both the I400 residue required for PRKAR1A binding and the LIR motif (Figs. 7A-B, S7A). In line with the suppression of RhoA-ROCK activity by PRKAR1A and FAM134B/C, the insensitivity of knockout cells to morphological changes was reversed by Y27632 (Fig. S7B-C).

**Fig. 7.**
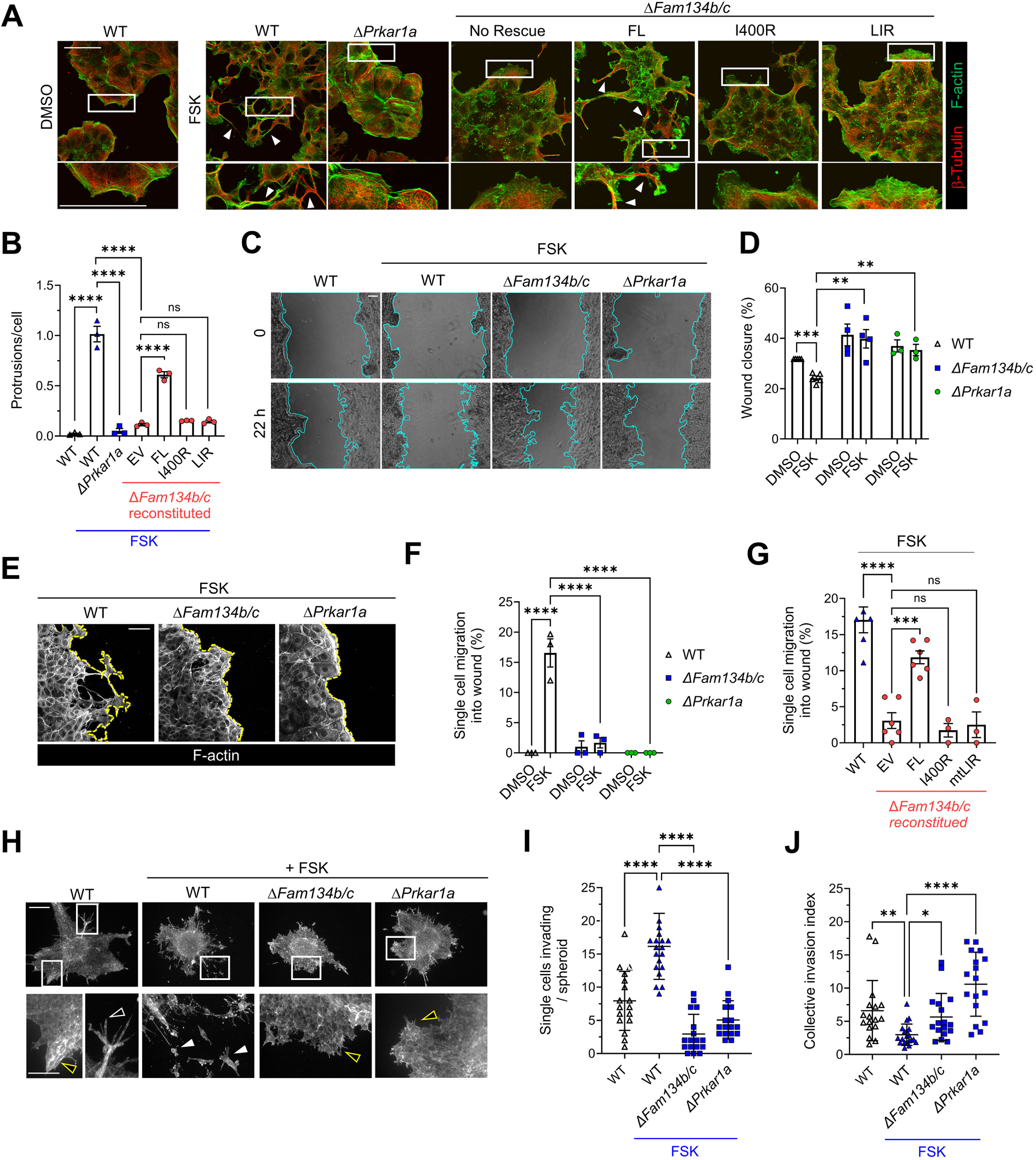
PRKAR1A and FAM134B regulate physiological changes in cell shape and invasion. **(A-B)** Confocal immunofluorescence microscopy of PDAC parental (WT, wild-type) or *ΔPrkar1a* cells or *ΔFam134b/c* cells, in the latter instance reconstituted with empty vector (No Rescue) or indicated variants of FAM134B (FL, full-length wild-type; I400R, PRKAR1A-binding deficient mutant; LIR, LC3- interacting region mutant), treated with DMSO or forskolin (FSK, 40 µM, 2 h), co-stained for F-actin (phalloidin, green) and β-Tubulin (red). Representative images in (**A**), white boxes delineate zoom panels, arrowheads indicate peripheral protrusions. Scale bars, 50 μm. Protrusions per cell quantified in (**B**) (*n* = 3 independent replicates, > 100 cells/condition, mean ± s.e.m., 1-way ANOVA and indicated Holm-Šídák multiple comparisons, **** = *P* < 0.0001, ns = *P* > 0.05). **(C-F)** Scratch wound healing assay on PDAC parental cells (WT) or Δ*Prkar1a* or Δ*Fam134b/c* treated with DMSO or FSK (40 µM). See also Movies S5-8. Representative phase contrast images in (**C**). Cyan line indicates wound front. Scale bar, 100 μm. Quantification of % wound closure after 22 h is shown in (**D**), values normalized to mean of WT DMSO control (*n* ≥ 3 independent replicates, mean ± s.e.m., 1-sample t-test FSK FL vs. DMSO FL, otherwise Holm-Šídák corrected multiple t-tests as indicated, *** = p < 0.001, ** = p < 0.01). Representative confocal fluorescence images after 7 h FSK-treatment (phalloidin staining for F-actin) in (**E**). Yellow line indicates wound edge, coherent in Δ*Prkar1a* or Δ*Fam134b/c*. Scale bar, 50 μm. Quantification of single cells migrating into the wound (without collective invasion of neighbours, see Methods) at 7 h DMSO or FSK treatment shown in (**F**) (*n* = 3 independent replicates, mean ± s.e.m., 2-way ANOVA and indicated Holm-Šídák multiple comparisons, **** = *P* < 0.0001). **(G)** PDAC WT or Δ*Fam134b/c* cells, the latter reconstituted with empty vector (No Rescue) or indicated variants of FAM134B were subjected to a scratch wound healing assay for 7 h with FSK treatment (40 µM) as described in (**C**-**F**). Quantification of single cells migrating into wound (n ≥ 3 independent replicates, mean ± s.e.m., 1-way ANOVA and indicated Holm-Šídák multiple comparisons. **** = *P* < 0.0001, *** = *P* < 0.001, ns = *P >* 0.05). **(H-J)** 3D spheroid collagen matrix invasion assay (48 h) with PDAC parental cells (WT), Δ*Prkar1a* or Δ*Fam134b/c,* treated with DMSO or FSK (40 µM, final 24 h). See also Movies S9-S12. Representative widefield fluorescence images of spheroids stained for F-actin (Phalloidin) shown in (**H**). White boxes indicate zoom panels. Black arrowheads indicate collective invasion, protrusive (yellow border) or multicellular streaming (white border). Solid white arrowheads indicate invasive single cells. Scale bars, 200 μm (main), 100 µm (zooms). Single cell invasion is quantified in (**I**), collective cell invasion is quantified in (**J**), as described in Materials and Methods (*n* = ≥ 17 spheroids from 3 independent replicates, mean ± s.e.m., 1-way ANOVA and indicated Holm-Šídák multiple comparisons. **** = *P* < 0.0001, ** = *P* < 0.01, * = *P* ≤ 0.05)

Behaviourally, cAMP-dependent morphological changes stimulated by FSK increased the rate at which single cells broke free of their neighbours at wound edges in two-dimensional scratch assays, resulting in reduced collective migration and wound closure (Fig. 7C-F, Movies S5-6). FSK-stimulated reduction in wound closure and increased single cell migration was absent in Δ*Fam134b/c* and Δ*Prkar1a* knockout cells (Fig. 7C-F, Movies S7-8). While single cell migration could be rescued in Δ*Fam134b/c* cells by FAM134B or FAM134C reconstitution (Fig. S7D-E), rescue was dependent on both I400 and the LIR of FAM134B (Fig. 7G), further underscoring the role of PRKAR1A-mediated clustering of FAM134B and LC3B. Single cell migration could also be rescued upon reconstitution of Δ*Prkar1a* cells with either FL or PKA_CAT_-binding deficient PRKAR1A (mtPS) (Fig. S7F-G). Finally, consolidating the role of PRKAR1A-FAM134B condensates in physiologic cell behaviour, we assayed their influence on cancer cell invasion into matrices, a process characterised by plasticity between modes of collective and solitary locomotion ^77^. Here, we observed a *Fam134b/c*- and *Prkar1a*-dependent FSK-stimulated switch from collective growth of cancer spheroids into the matrix to breakaway invasion of single cells (Fig. 7H-J, Movies S9-S12). Notably, this *Fam134b/c*-dependent switch in invasion modality was also seen when stimulating cytosolic cAMP via a physiologic G-protein coupled receptor agonist, adrenaline (epinephrine) (Fig. S7H-I).

These data, taken together, reveal that the physiological relevance of PRKAR1A condensates in regulating FAM134B/C mobilisation and RhoA activity is control of cell morphological plasticity in response to cAMP.

## Discussion

Screening identified a cohort of proteins associated with the cytoplasmic regions of ER-phagy receptors. For FAM134B/C, we established that a subset of such interactors regulates receptor activation and ER-phagy flux. In particular, through detailed molecular investigation of an exemplary interactor, PRKAR1A, we established that the cytoplasmic region of FAM134B/C, because of direct interaction via an amphipathic α-helix (α_AMPHI_), can control both ER and cytoplasmic remodelling as part of a coherent cell signalling response (Fig. 8). In this scenario, FAM134B/C activation is dependent upon a non-canonical, PKA_CAT_-independent function of cAMP-bound PRKAR1A. Liquid condensates of PRKAR1A form ER contact sites and facilitate local clustering of FAM134B with LC3B. These ER contact sites can be considered degradation hubs. Firstly, these sites are where the FAM134B/C ER-phagy receptors are sequestered into local lysosomes. Furthermore, FAM134B/C also interacts with the small GTPase RhoA, leading to its lysosomal co-degradation within these hubs. The physiologic result of condensate-mediated degradation of FAM134B/C and RhoA is inhibition of cytoplasmic signalling to the cytoskeleton, which in turn is required for cAMP-driven peripheral membrane dynamics and switching of cell behaviour from collective to single cell migration and invasion.

**Fig 8.**
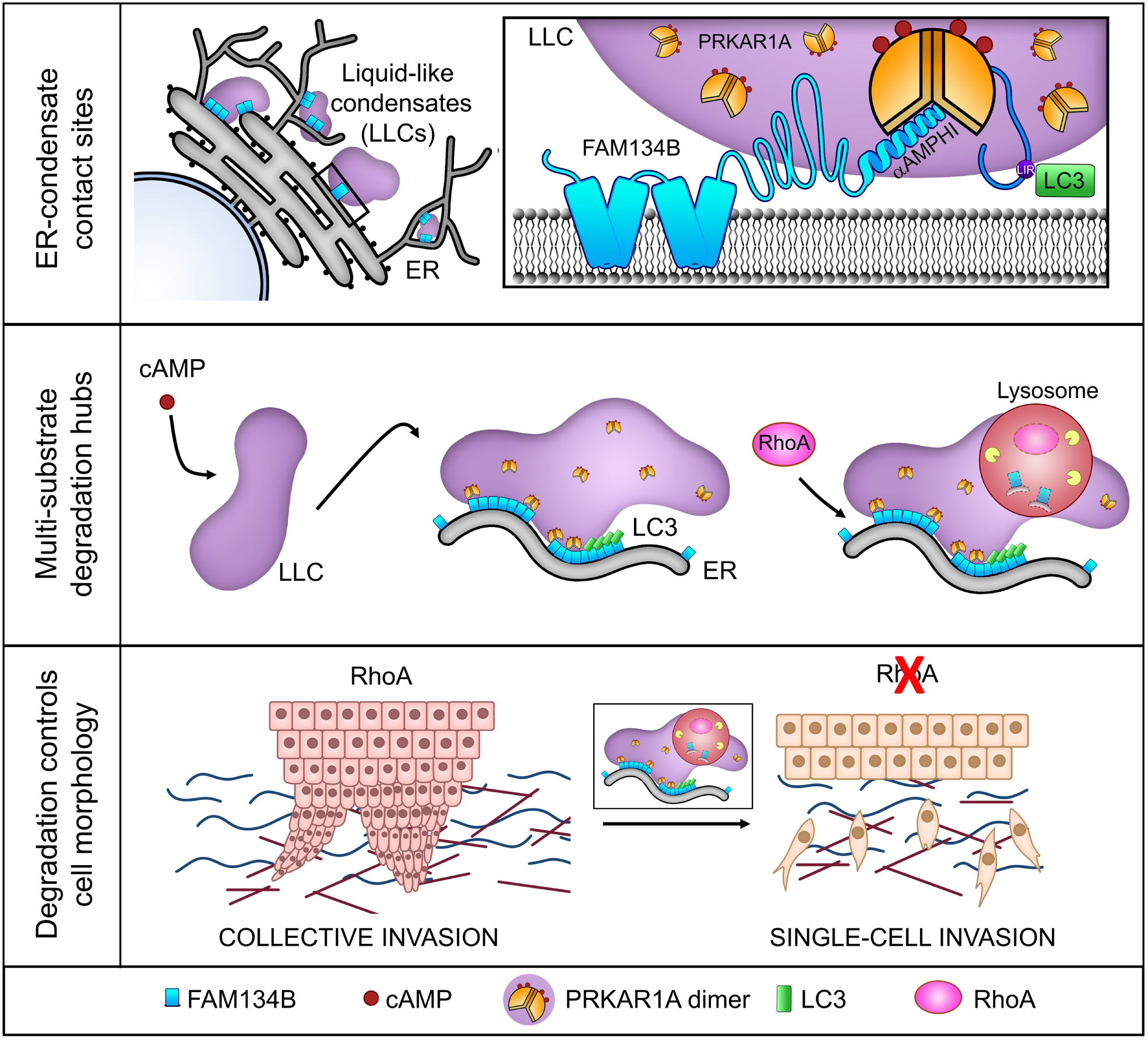
Schematic illustration of the proposed model. Liquid-like condensates (LLCs) of cAMP-bound PRKAR1A form close contacts with the ER. At this site, PRKAR1A dimers interact with the ER integral FAM134B/C via a direct interaction of the dimerization domain in the N-terminus (NT) of PRKAR1A with the amphipathic helix (α_AMPHI_) in FAM134B (top panel). Mechanistically, LLCs locally cluster immobile FAM134B/C in the ER with LC3B. These organellar contact sites between ER and LLCs are a hub for lysosomal degradation of the FAM134B/C-enriched ER and co-degradation of the master regulator of cell morphology and FAM134B/C interactor, RhoA (middle panel). Physiologically, RhoA degradation via FAM134B/C regulates peripheral membrane dynamics and switches the invasion modality of cancer cells in 3D, e.g., from collective invasion to single cell invasion (bottom panel).

Our findings first broaden the cellular roles of FAM134B/C ER-phagy receptors beyond ER remodelling and proteostasis. Indeed, the ER is increasingly appreciated as a hub for co-ordinating cellular events via interorganellar and plasma membrane contacts ^1^. For example, complementary to our findings, wherein FAM134B/C-rich contact sites with condensates regulate cell morphology, other recent reports have suggested that ER dynamics could regulate overall cell morphology, for instance via microtubules or ER-plasma membrane contacts ^78–80^. We speculate that broader effects of ER structural remodelling by ER-phagy, impinging on such other contact sites, could co-operate with regulation of RhoA in control of cell morphology.

Secondly, this study also establishes a role for liquid-like condensates in regulation of ER-phagy and cell morphology, in both established cell culture and in three-dimensional cancer cell invasion settings. However, it is notable that analogous morphological transitions occur in orthogonal settings, such as neuronal differentiation; we note that substantial changes in FAM134B/C flux or PRKAR1A degradation are evidenced therein within several proteomic datasets ^81–83^. This could suggest that the principle of coordinated ER and cell morphology control by ER receptors and PRKAR1A condensates is applicable in other, diverse biological contexts.

Thirdly, at a molecular level, this study also reveals insight into the roles of the extensive cytoplasmic domains of FAM134B/C. The phospholipid- and cholesterol-binding properties of the small amphipathic helical regions associated with the RHD are known to regulate ER-phagy ^66, 84^. However, for the first time we reveal that not only does FAM134B/C bind to cytoplasmic RhoA, but that an amphipathic helix (α_AMPHI_) within the extensive IDR of FAM134B/C favours docking with the hydrophobic pocket created by cytoplasmic PRKAR1A dimers. Intriguingly, it is possible that, in the absence of PRKAR1A, that this helix could also be stabilised by hydrophobic interaction with other as-yet-unknown proteins or even lipids. Indeed, some other LIR-independent interactions of the C-terminus of FAM134B have recently been suggested: with mATG9 (part of the core autophagy machinery); and with the ER membrane protein STIM1 ^85, 86^. The latter interaction was corroborated by its presence in the proximity proteomics presented herein (Table S3), underscoring a potentially promiscuous nature of FAM134B/C cytoplasmic interactions and involvement in cell signalling responses.

While IDR regions are often involved in liquid-like condensate formation, FAM134B/C molecules within clusters are immobile. This is a sharp contrast with the emerging role of liquid-like condensation of cargo receptors in selective autophagy pathways such as aggrephagy and mitophagy ^45, 51–53, 87^.

However, PRKAR1A, as a cytoplasmic IDR protein, can be viewed as an ancillary, liquid-like co-receptor for ER-phagy through its formation of condensates at ER contact sites, which serve to cluster FAM134B/C with LC3B. This finding thus extends the role of liquid-like condensation in control of selective autophagy receptor function to ER-phagy. Additionally, it reveals how FAM134B/C receptors are able to exploit the rapid, signal-responsive nature of condensate formation to respond to afferent cytoplasmic signals to coordinate downstream degradation pathways.

Fourthly, this work ascribes a new function to the recently-described, liquid-condensed form of PRKAR1A. While PRKAR1A condensation has been suggested to regulate PKA_CAT_ signalling by cAMP sequestration ^54, 55, 88, 89^, the implications of this phenomenon beyond – and independent of – kinase regulation are not elucidated. Here, we reveal a PKA_CAT_-independent role for liquid-condensed PRKAR1A in coordinated regulation of ER-phagy and RhoA, in conjunction with FAM134B/C. Indeed, this work reveals a major new function for PRKAR1A as an apical, signal-responsive facilitator of selective autophagy. However, this finding is not exclusive with suggestions that PRKAR1A can regulate mTOR ^90^ or can be a downstream target for autophagic degradation, likely via interaction with the LIR-motif protein AKAP11 (not required for FAM134B interaction, as shown herein) ^64, 65, 82, 91^. Indeed, recruitment of PRKAR1A to nascent autophagosomes and subsequent degradation could potentially modulate PKA_CAT_ kinase outputs during cAMP responses ^65, 91^; this is not mutually exclusive with the lack of PKA_CAT_ requirement for ER-phagy and RhoA regulation.

Finally, the physiological relevance of liquid-like condensate-mediated regulation of FAM134B/C is clear in the control of RhoA-dependent morphological plasticity, including the ability to transition from collective to single cell motility and invasion in two- and three-dimensional settings. Interestingly, the soluble autophagy receptor p62 was reported to bind to active, GTP-bound RhoA ^92^. Although we find that FAM134B/C binds selectively to inactive RhoA, the observation of enhanced activity of RhoA in

*Atg5* null cells in this study is not at odds with our data. In any instance, the PRKAR1A-FAM134B/C axis should in future be investigated for its role in cancer metastasis. Furthermore, the functions of FAM134B/C described herein will be corrupted in other diseases, such as hereditary sensory neuropathies associated with truncations in the C-terminal domain of FAM134B ^6^, potentially extending their mechanistic aetiologies beyond defective ER proteostasis.

## Supporting information

Raw immunoblots

## Acknowledgements

We thank Carla Salomo-Coll for her help with the schematic illustration; Laura Murphy and Martin Lee from the Advanced Imaging Resource, Institute of Genetics and Cancer, University of Edinburgh and Lorna Hodgson from Wolfson Bioimaging Facility, University of Bristol, for imaging technical support. We thank Neil O. Carragher from Institute of Genetics and Cancer, University of Edinburgh for IncuCyte S3 analyses. We thank Stephen Brown and Jeffrey Joseph from MRC Human Genetics Unit, Institute of Genetics and Cancer, University of Edinburgh for DNA sequencing. This work was funded by a CRUK Senior Fellowship to SW (C20685/A29576).

## Extended Methods

### Key Resource Table

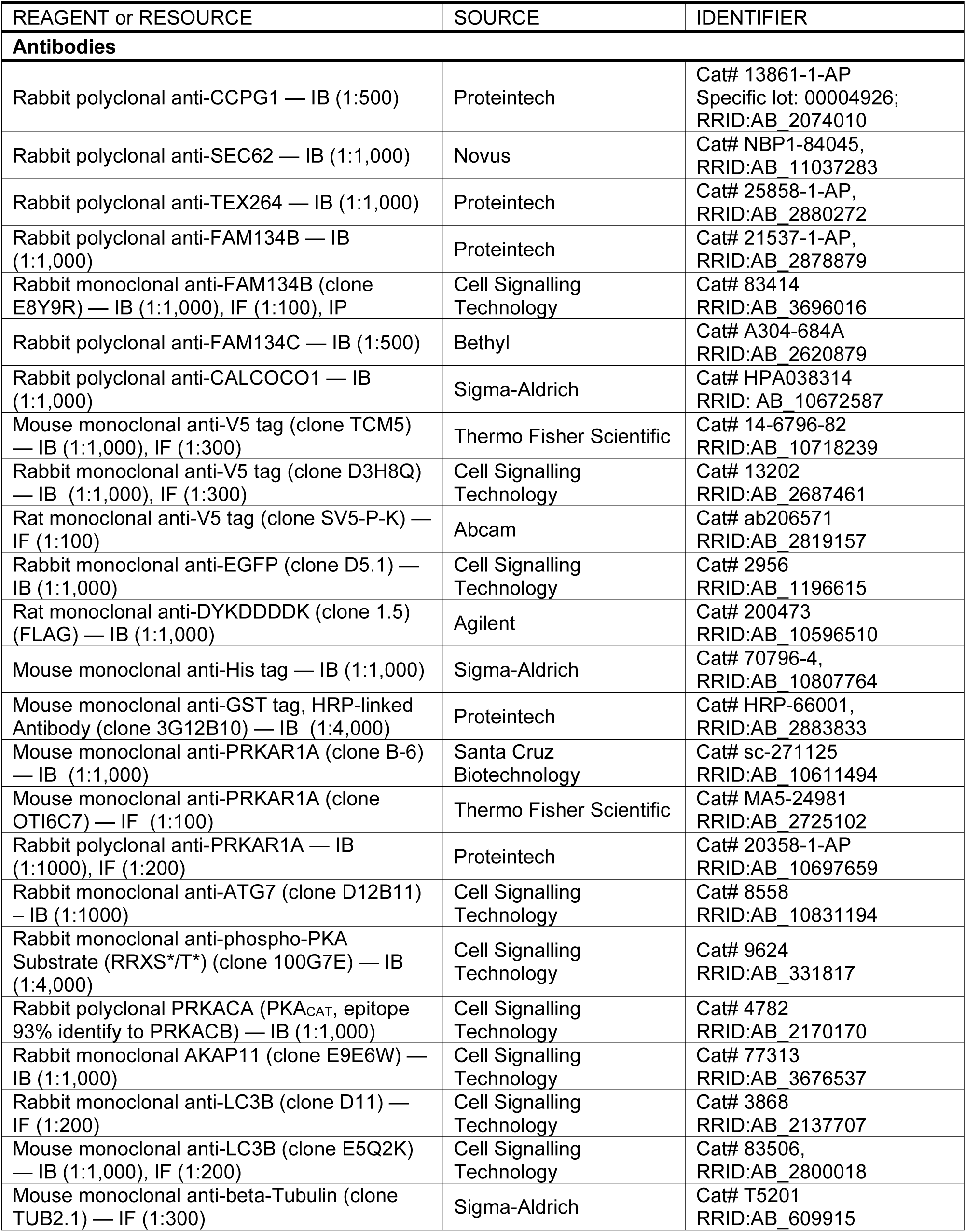
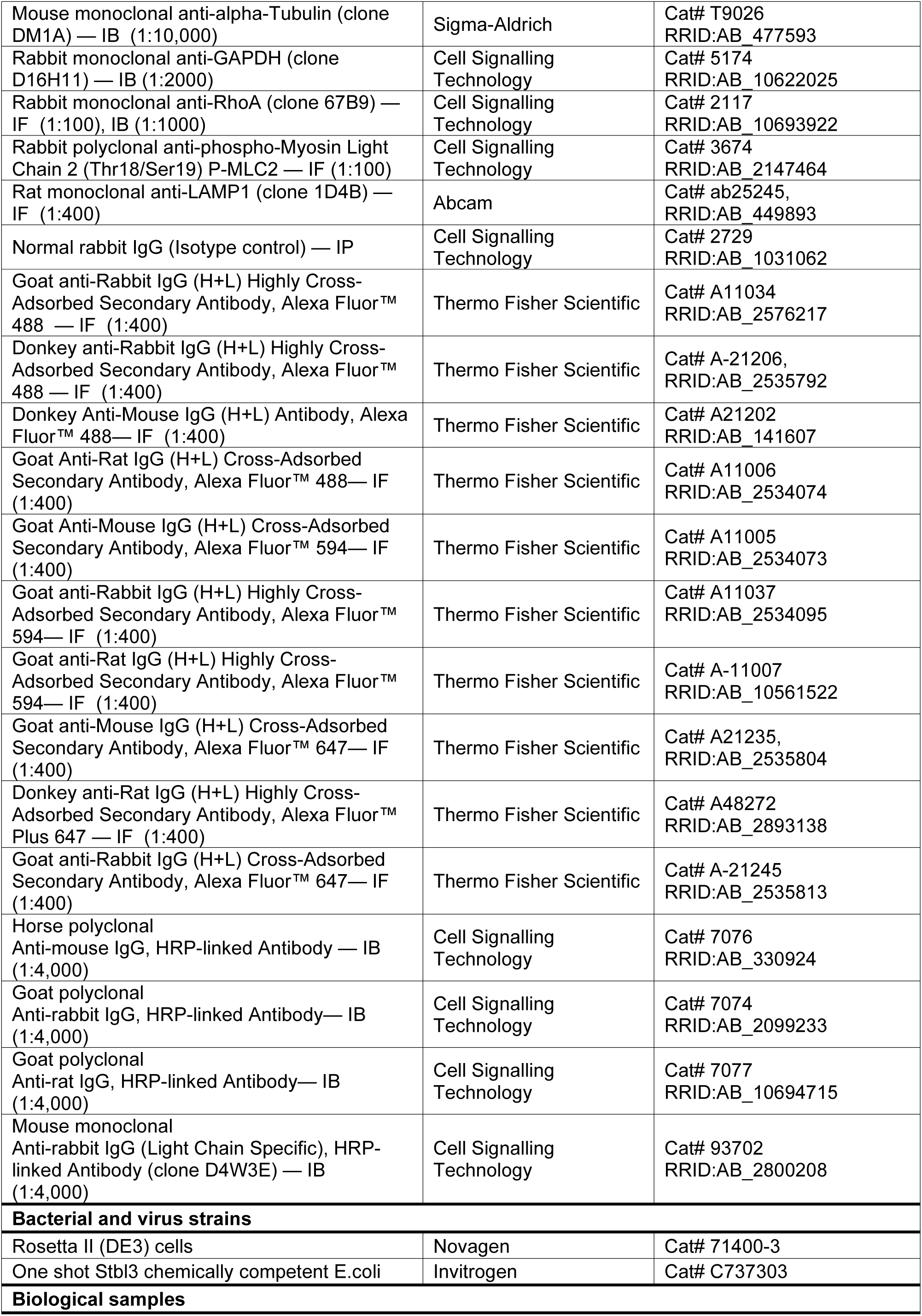
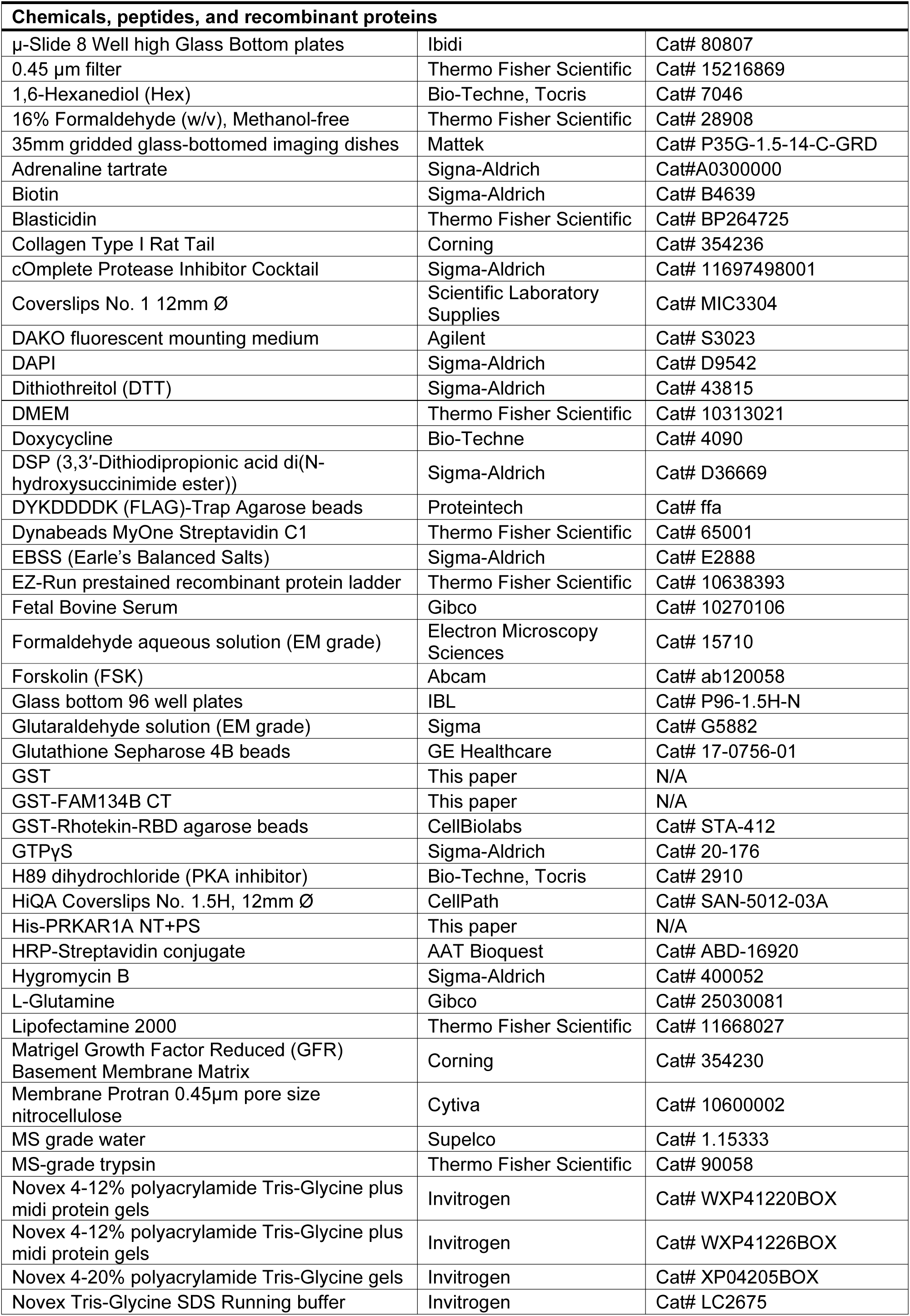
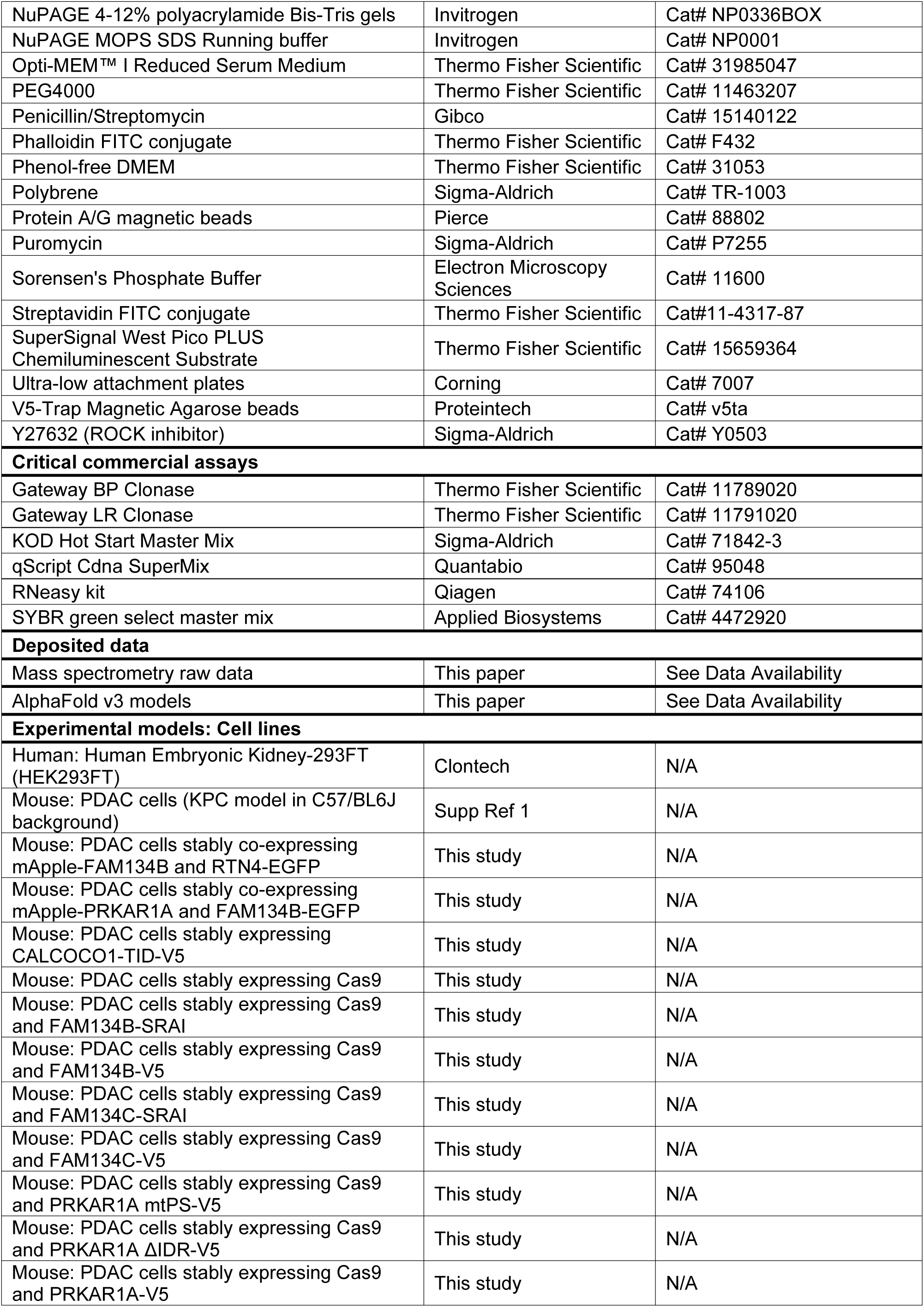
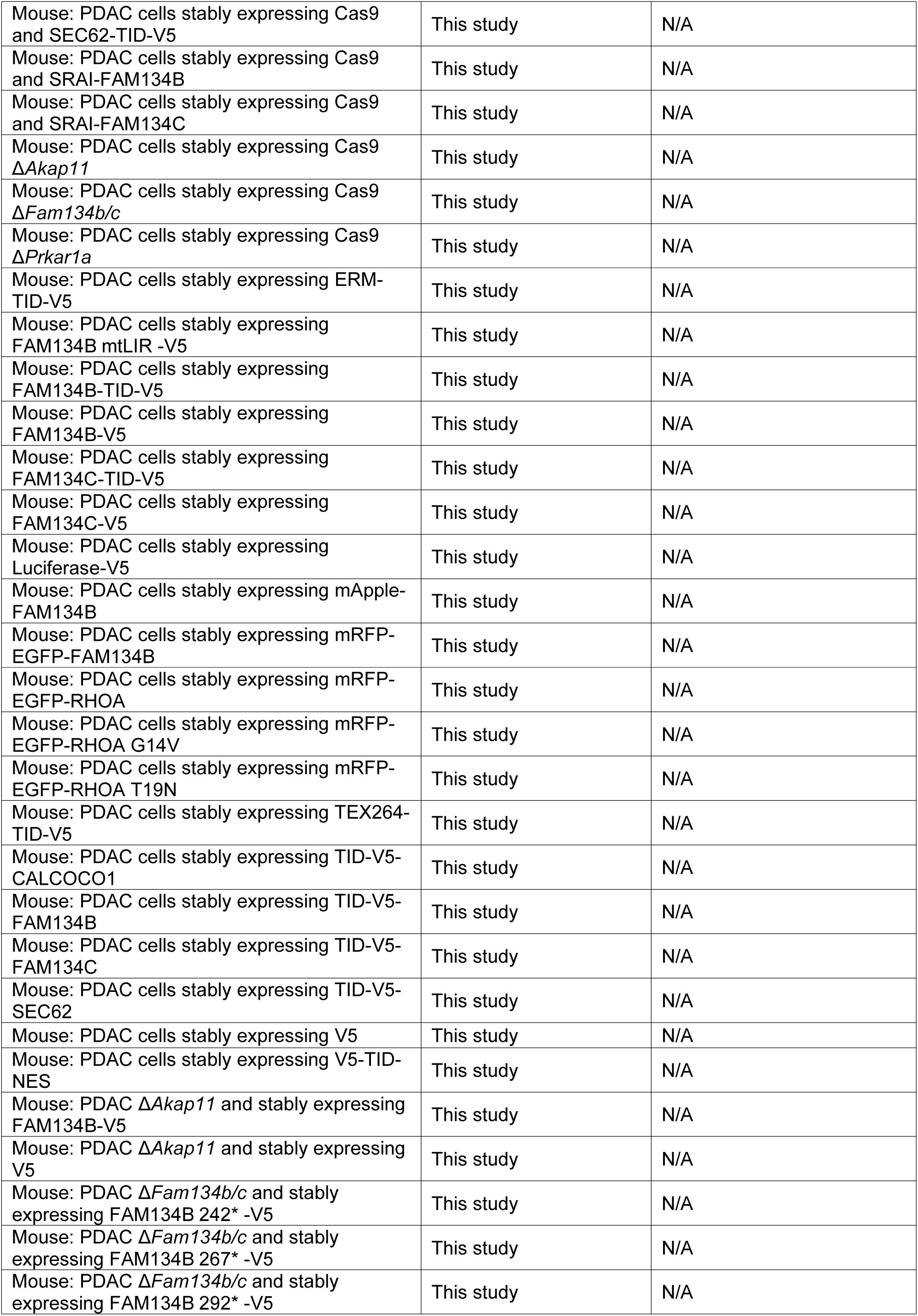
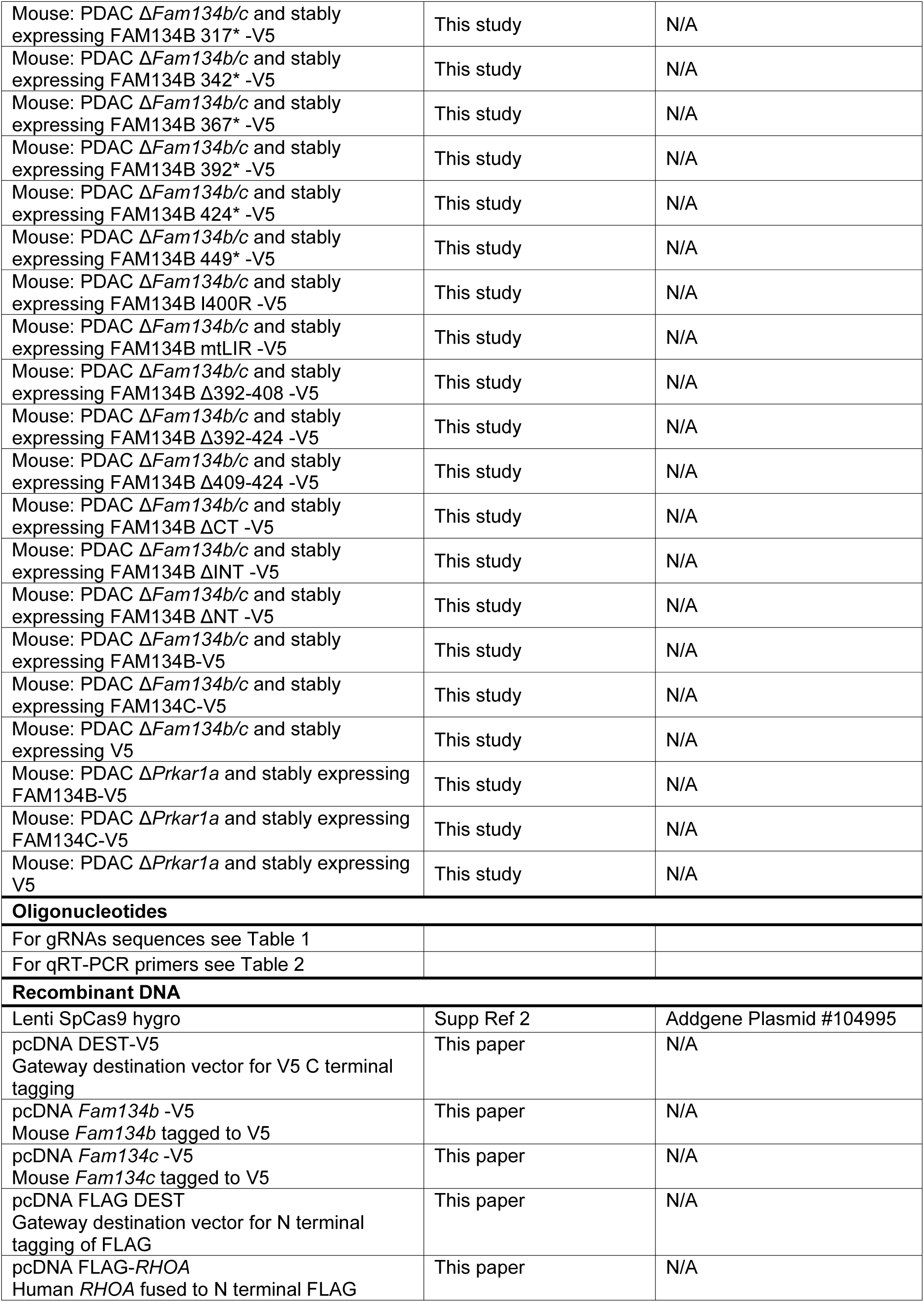
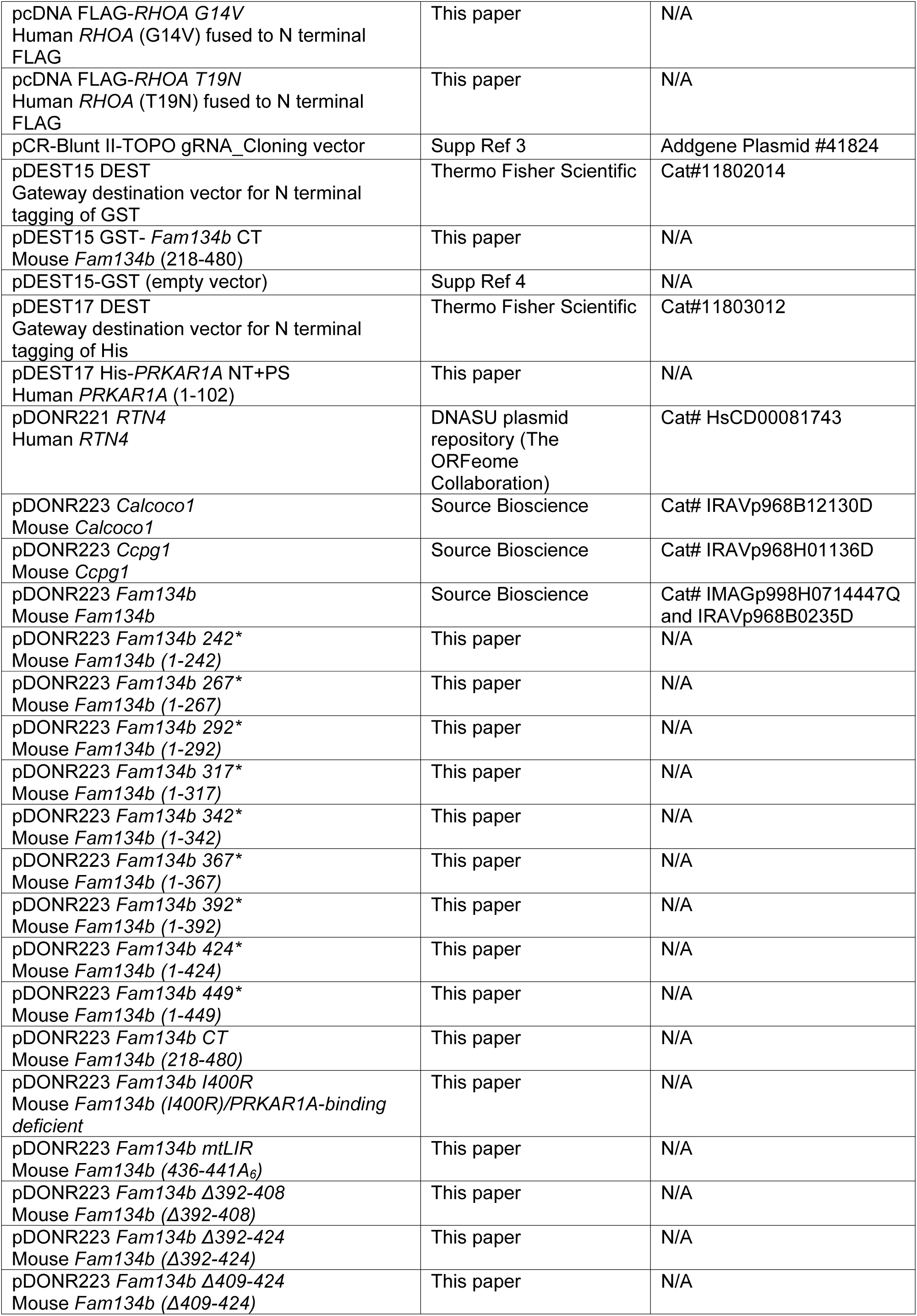
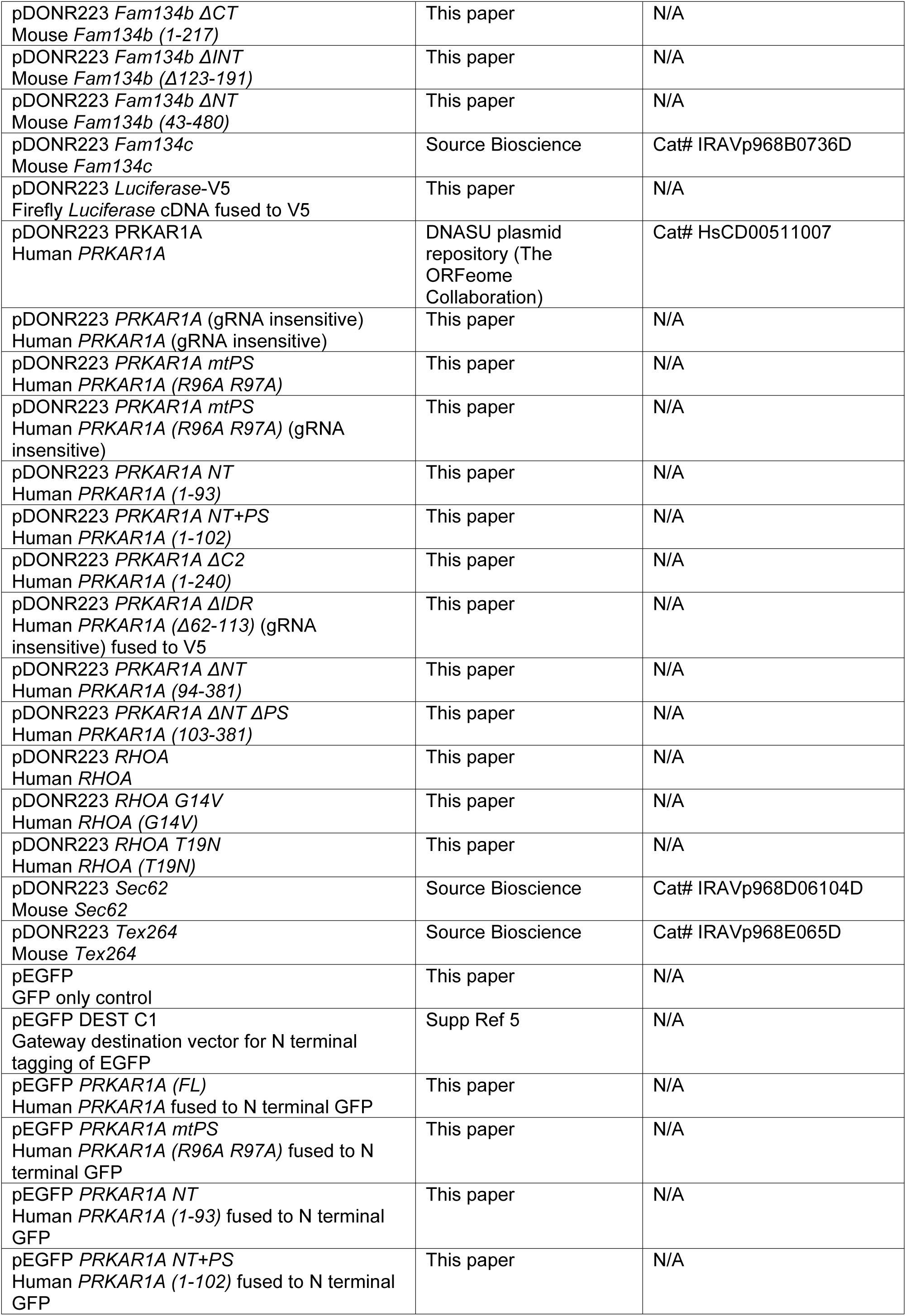
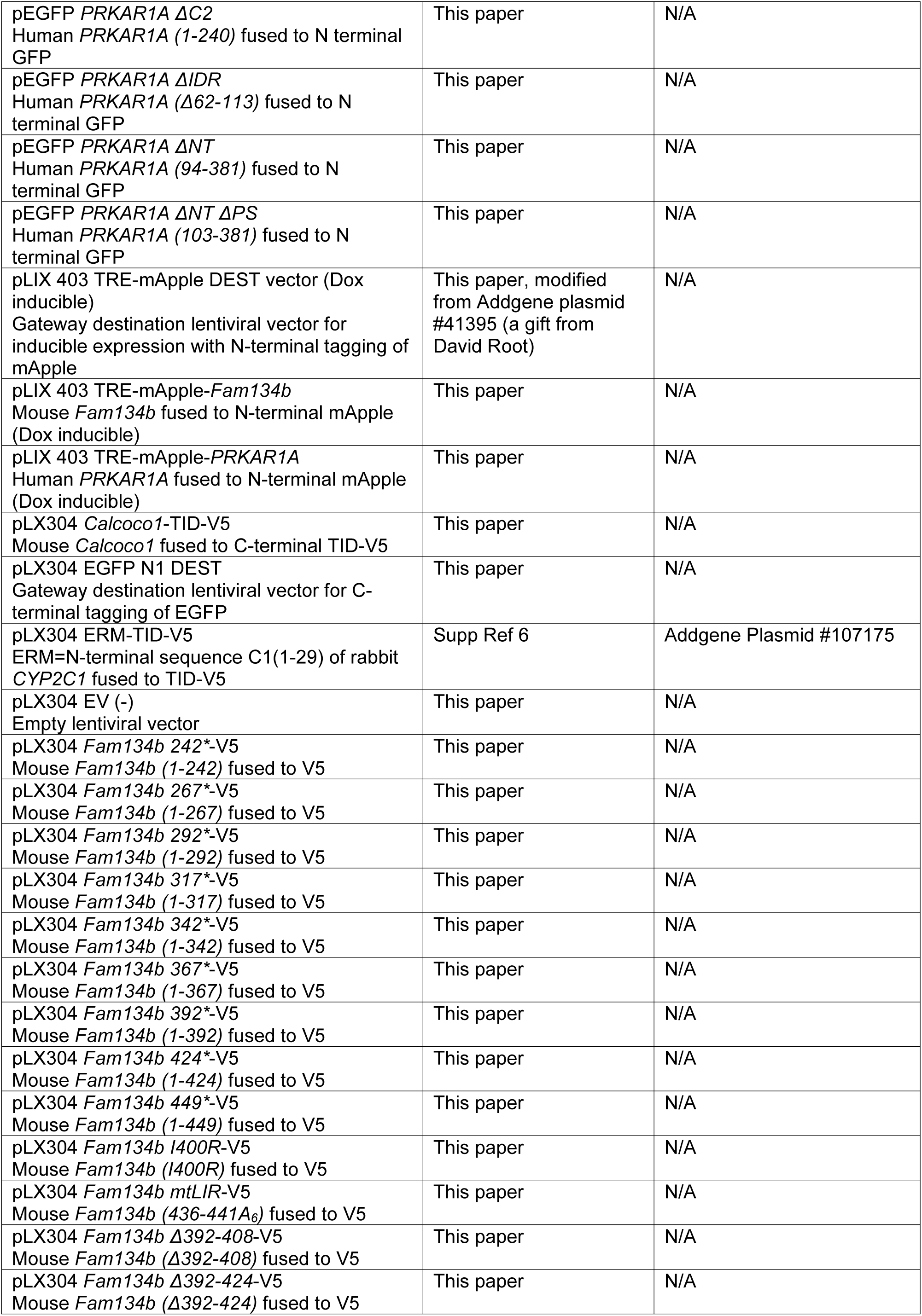
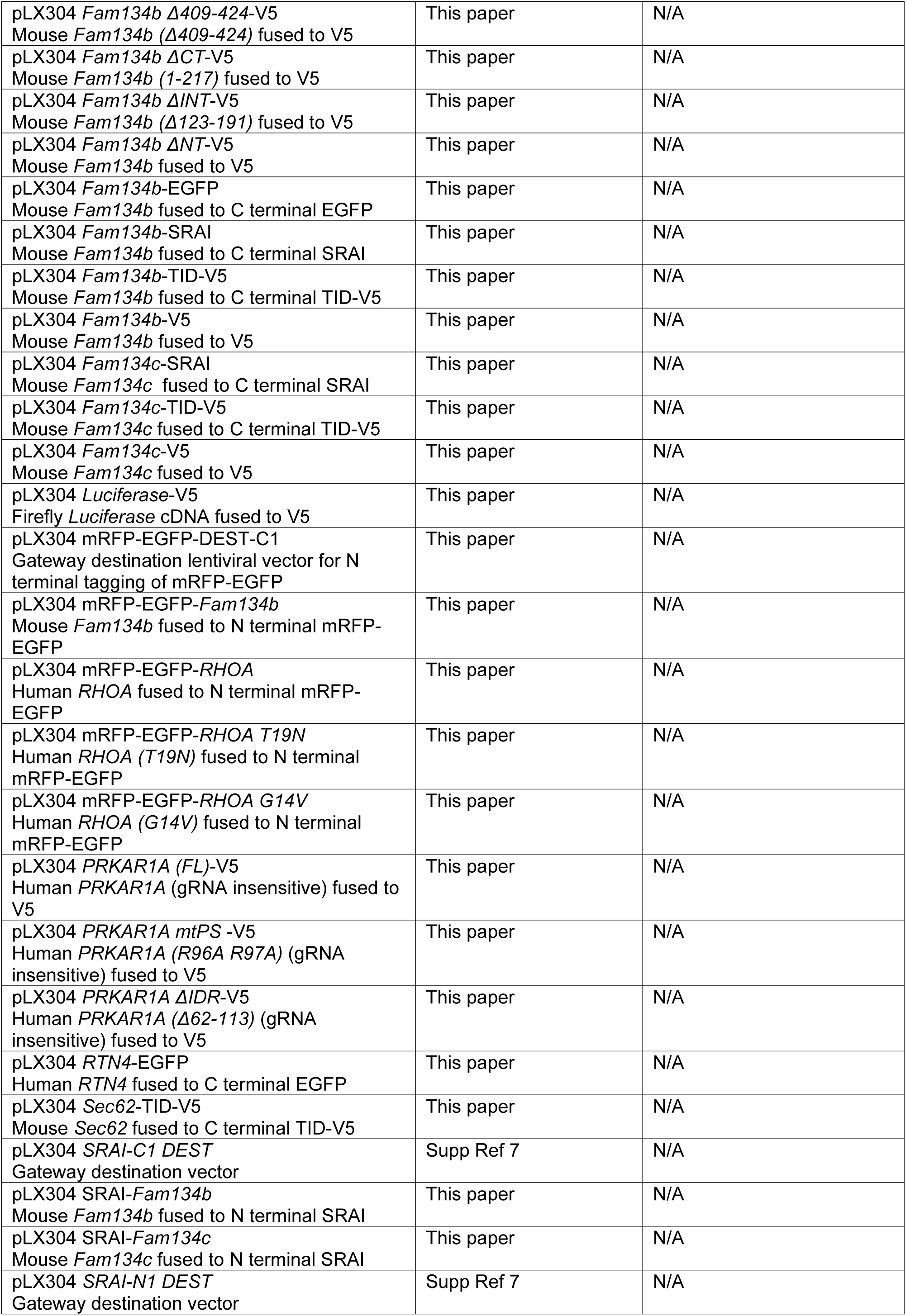
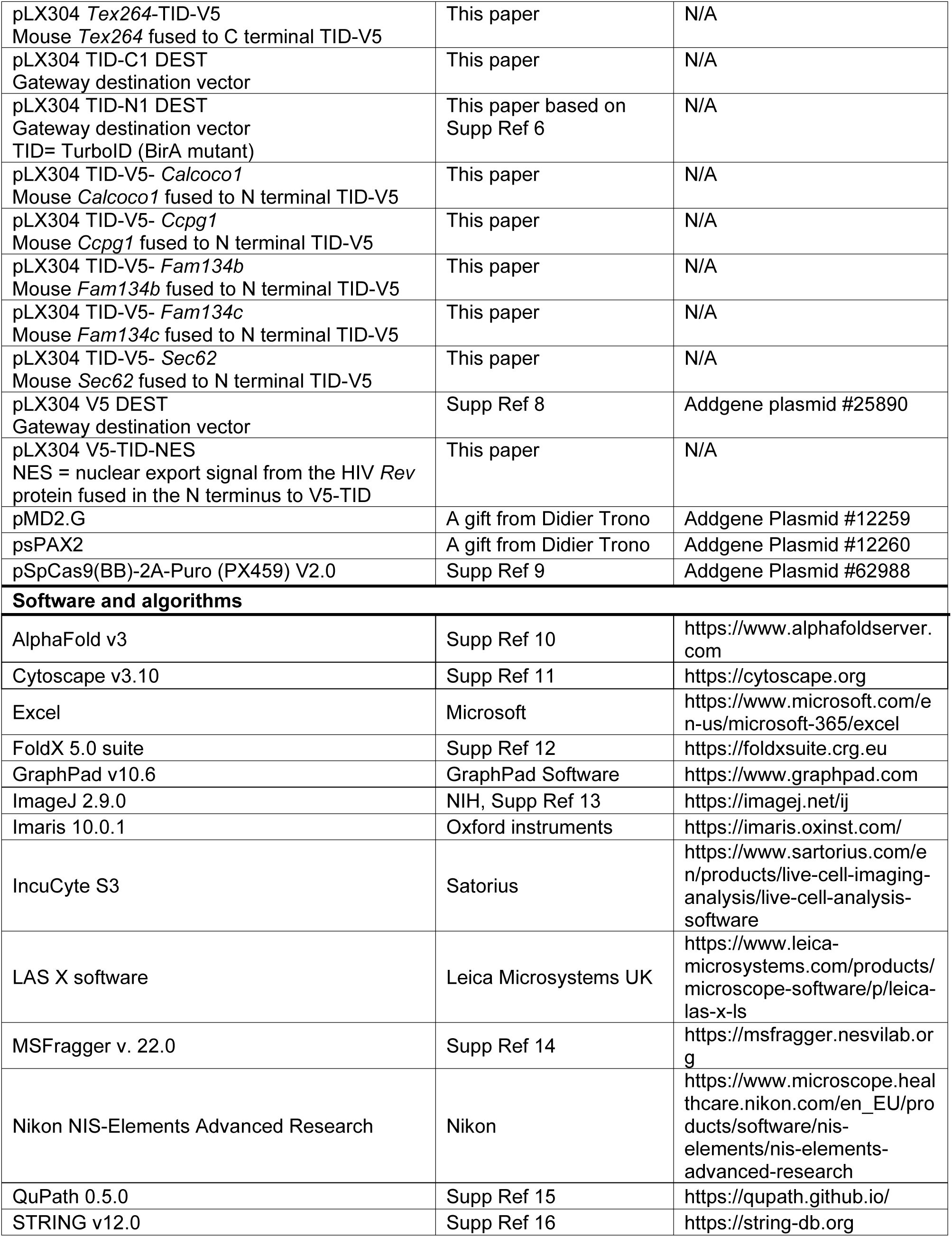

| REAGENT or RESOURCE | SOURCE | IDENTIFIER |
| --- | --- | --- |
| **Antibodies** |  |  |
| Rabbit polyclonal anti-CCPG1 — IB (1:500) | Proteintech | Cat# 13861-1-AP<br>Specific lot: 00004926;<br>RRID:AB_2074010 |
| Rabbit polyclonal anti-SEC62 — IB (1:1,000) | Novus | Cat# NBP1-84045,<br>RRID:AB_11037283 |
| Rabbit polyclonal anti-TEX264 — IB (1:1,000) | Proteintech | Cat# 25858-1-AP,<br>RRID:AB_2880272 |
| Rabbit polyclonal anti-FAM134B — IB (1:1,000) | Proteintech | Cat# 21537-1-AP,<br>RRID:AB_2878879 |
| Rabbit monoclonal anti-FAM134B (clone E8Y9R) — IB (1:1,000), IF (1:100), IP | Cell Signalling Technology | Cat# 83414<br>RRID:AB_3696016 |
| Rabbit polyclonal anti-FAM134C — IB (1:500) | Bethyl | Cat# A304-684A<br>RRID:AB_2620879 |
| Rabbit polyclonal anti-CALCOCO1 — IB (1:1,000) | Sigma-Aldrich | Cat# HPA038314<br>RRID:AB_10672587 |
| Mouse monoclonal anti-V5 tag (clone TCM5) — IB (1:1,000), IF (1:300) | Thermo Fisher Scientific | Cat# 14-6796-82<br>RRID:AB_10718239 |
| Rabbit monoclonal anti-V5 tag (clone D3H8Q) — IB (1:1,000), IF (1:300) | Cell Signalling Technology | Cat# 13202<br>RRID:AB_2687461 |
| Rat monoclonal anti-V5 tag (clone SV5-P-K) — IF (1:100) | Abcam | Cat# ab206571<br>RRID:AB_2819157 |
| Rabbit monoclonal anti-EGFP (clone D5.1) — IB (1:1,000) | Cell Signalling Technology | Cat# 2956<br>RRID:AB_1196615 |
| Rat monoclonal anti-DYKDDDDK (clone 1.5) (FLAG) — IB (1:1,000) | Agilent | Cat# 200473<br>RRID:AB_10596510 |
| Mouse monoclonal anti-His tag — IB (1:1,000) | Sigma-Aldrich | Cat# 70796-4,<br>RRID:AB_10807764 |
| Mouse monoclonal anti-GST tag, HRP-linked Antibody (clone 3G12B10) — IB (1:4,000) | Proteintech | Cat# HRP-66001,<br>RRID:AB_2883833 |
| Mouse monoclonal anti-PRKAR1A (clone B-6) — IB (1:1,000) | Santa Cruz Biotechnology | Cat# sc-271125<br>RRID:AB_10611494 |
| Mouse monoclonal anti-PRKAR1A (clone OTI6C7) — IF (1:100) | Thermo Fisher Scientific | Cat# MA5-24981<br>RRID:AB_2725102 |
| Rabbit polyclonal anti-PRKAR1A — IB (1:1000), IF (1:200) | Proteintech | Cat# 20358-1-AP<br>RRID:AB_10697659 |
| Rabbit monoclonal anti-ATG7 (clone D12B11) — IB (1:1000) | Cell Signalling Technology | Cat# 8558<br>RRID:AB_10831194 |
| Rabbit monoclonal anti-phospho-PKA Substrate (RRXS*/T*) (clone 100G7E) — IB (1:4,000) | Cell Signalling Technology | Cat# 9624<br>RRID:AB_331817 |
| Rabbit polyclonal PRKACA (PKA<sub>CAT</sub>, epitope 93% identify to PRKACB) — IB (1:1,000) | Cell Signalling Technology | Cat# 4782<br>RRID:AB_2170170 |
| Rabbit monoclonal AKAP11 (clone E9E6W) — IB (1:1,000) | Cell Signalling Technology | Cat# 77313<br>RRID:AB_3676537 |
| Rabbit monoclonal anti-LC3B (clone D11) — IF (1:200) | Cell Signalling Technology | Cat# 3868<br>RRID:AB_2137707 |
| Mouse monoclonal anti-LC3B (clone E5Q2K) — IB (1:1,000), IF (1:200) | Cell Signalling Technology | Cat# 83506,<br>RRID:AB_2800018 |
| Mouse monoclonal anti-beta-Tubulin (clone TUB2.1) — IF (1:300) | Sigma-Aldrich | Cat# T5201<br>RRID:AB_609915 |
| Mouse monoclonal anti-alpha-Tubulin (clone DM1A) — IB (1:10,000) | Sigma-Aldrich | Cat# T9026<br>RRID:AB_477593 |
| Rabbit monoclonal anti-GAPDH (clone D16H11) — IB (1:2000) | Cell Signalling Technology | Cat# 5174<br>RRID:AB_10622025 |
| Rabbit monoclonal anti-RhoA (clone 67B9) — IF (1:100), IB (1:1000) | Cell Signalling Technology | Cat# 2117<br>RRID:AB_10693922 |
| Rabbit polyclonal anti-phospho-Myosin Light Chain 2 (Thr18/Ser19) P-MLC2 — IF (1:100) | Cell Signalling Technology | Cat# 3674<br>RRID:AB_2147464 |
| Rat monoclonal anti-LAMP1 (clone 1D4B) — IF (1:400) | Abcam | Cat# ab25245,<br>RRID:AB_449893 |
| Normal rabbit IgG (Isotype control) — IP | Cell Signalling Technology | Cat# 2729<br>RRID:AB_1031062 |
| Goat anti-Rabbit IgG (H+L) Highly Cross-Adsorbed Secondary Antibody, Alexa Fluor™ 488 — IF (1:400) | Thermo Fisher Scientific | Cat# A11034<br>RRID:AB_2576217 |
| Donkey anti-Rabbit IgG (H+L) Highly Cross-Adsorbed Secondary Antibody, Alexa Fluor™ 488 — IF (1:400) | Thermo Fisher Scientific | Cat# A-21206,<br>RRID:AB_2535792 |
| Donkey Anti-Mouse IgG (H+L) Antibody, Alexa Fluor™ 488 — IF (1:400) | Thermo Fisher Scientific | Cat# A21202<br>RRID:AB_141607 |
| Goat Anti-Rat IgG (H+L) Cross-Adsorbed Secondary Antibody, Alexa Fluor™ 488 — IF (1:400) | Thermo Fisher Scientific | Cat# A11006<br>RRID:AB_2534074 |
| Goat Anti-Mouse IgG (H+L) Cross-Adsorbed Secondary Antibody, Alexa Fluor™ 594 — IF (1:400) | Thermo Fisher Scientific | Cat# A11005<br>RRID:AB_2534073 |
| Goat anti-Rabbit IgG (H+L) Highly Cross-Adsorbed Secondary Antibody, Alexa Fluor™ 594 — IF (1:400) | Thermo Fisher Scientific | Cat# A11037<br>RRID:AB_2534095 |
| Goat anti-Rat IgG (H+L) Highly Cross-Adsorbed Secondary Antibody, Alexa Fluor™ 594 — IF (1:400) | Thermo Fisher Scientific | Cat# A-11007<br>RRID:AB_10561522 |
| Goat anti-Mouse IgG (H+L) Cross-Adsorbed Secondary Antibody, Alexa Fluor™ 647 — IF (1:400) | Thermo Fisher Scientific | Cat# A21235,<br>RRID:AB_2535804 |
| Donkey anti-Rat IgG (H+L) Highly Cross-Adsorbed Secondary Antibody, Alexa Fluor™ Plus 647 — IF (1:400) | Thermo Fisher Scientific | Cat# A48272<br>RRID:AB_2893138 |
| Goat anti-Rabbit IgG (H+L) Cross-Adsorbed Secondary Antibody, Alexa Fluor™ 647 — IF (1:400) | Thermo Fisher Scientific | Cat# A-21245<br>RRID:AB_2535813 |
| Horse polyclonal Anti-mouse IgG, HRP-linked Antibody — IB (1:4,000) | Cell Signalling Technology | Cat# 7076<br>RRID:AB_330924 |
| Goat polyclonal Anti-rabbit IgG, HRP-linked Antibody — IB (1:4,000) | Cell Signalling Technology | Cat# 7074<br>RRID:AB_2099233 |
| Goat polyclonal Anti-rat IgG, HRP-linked Antibody — IB (1:4,000) | Cell Signalling Technology | Cat# 7077<br>RRID:AB_10694715 |
| Mouse monoclonal Anti-rabbit IgG (Light Chain Specific), HRP-linked Antibody (clone D4W3E) — IB (1:4,000) | Cell Signalling Technology | Cat# 93702<br>RRID:AB_2800208 |
| **Bacterial and virus strains** |  |  |
| Rosetta II (DE3) cells | Novagen | Cat# 71400-3 |
| One shot Stbl3 chemically competent E.coli | Invitrogen | Cat# C737303 |
| **Biological samples** |  |  |

| **Chemicals, peptides, and recombinant proteins** |  |  |
| --- | --- | --- |
| μ-Slide 8 Well high Glass Bottom plates | Ibidi | Cat# 80807 |
| 0.45 μm filter | Thermo Fisher Scientific | Cat# 15216869 |
| 1,6-Hexanediol (Hex) | Bio-Techne, Tocris | Cat# 7046 |
| 16% Formaldehyde (w/v), Methanol-free | Thermo Fisher Scientific | Cat# 28908 |
| 35mm gridded glass-bottomed imaging dishes | Mattek | Cat# P35G-1.5-14-C-GRD |
| Adrenaline tartrate | Sigma-Aldrich | Cat#A0300000 |
| Biotin | Sigma-Aldrich | Cat# B4639 |
| Blasticidin | Thermo Fisher Scientific | Cat# BP264725 |
| Collagen Type I Rat Tail | Corning | Cat# 354236 |
| cOmplete Protease Inhibitor Cocktail | Sigma-Aldrich | Cat# 11697498001 |
| Coverslips No. 1 12mm Ø | Scientific Laboratory Supplies | Cat# MIC3304 |
| DAKO fluorescent mounting medium | Agilent | Cat# S3023 |
| DAPI | Sigma-Aldrich | Cat# D9542 |
| Dithiothreitol (DTT) | Sigma-Aldrich | Cat# 43815 |
| DMEM | Thermo Fisher Scientific | Cat# 10313021 |
| Doxycycline | Bio-Techne | Cat# 4090 |
| DSP (3,3'-Dithiodipropionic acid di(N-hydroxysuccinimide ester)) | Sigma-Aldrich | Cat# D36669 |
| DYKDDDDK (FLAG)-Trap Agarose beads | Proteintech | Cat# ffa |
| Dynabeads MyOne Streptavidin C1 | Thermo Fisher Scientific | Cat# 65001 |
| EBSS (Earle's Balanced Salts) | Sigma-Aldrich | Cat# E2888 |
| EZ-Run prestained recombinant protein ladder | Thermo Fisher Scientific | Cat# 10638393 |
| Fetal Bovine Serum | Gibco | Cat# 10270106 |
| Formaldehyde aqueous solution (EM grade) | Electron Microscopy Sciences | Cat# 15710 |
| Forskolin (FSK) | Abcam | Cat# ab120058 |
| Glass bottom 96 well plates | IBL | Cat# P96-1.5H-N |
| Glutaraldehyde solution (EM grade) | Sigma | Cat# G5882 |
| Glutathione Sepharose 4B beads | GE Healthcare | Cat# 17-0756-01 |
| GST | This paper | N/A |
| GST-FAM134B CT | This paper | N/A |
| GST-Rhotekin-RBD agarose beads | CellBiolabs | Cat# STA-412 |
| GTPγS | Sigma-Aldrich | Cat# 20-176 |
| H89 dihydrochloride (PKA inhibitor) | Bio-Techne, Tocris | Cat# 2910 |
| HiQA Coverslips No. 1.5H, 12mm Ø | CellPath | Cat# SAN-5012-03A |
| His-PRKAR1A NT+PS | This paper | N/A |
| HRP-Streptavidin conjugate | AAT Bioquest | Cat# ABD-16920 |
| Hygromycin B | Sigma-Aldrich | Cat# 400052 |
| L-Glutamine | Gibco | Cat# 25030081 |
| Lipofectamine 2000 | Thermo Fisher Scientific | Cat# 11668027 |
| Matrigel Growth Factor Reduced (GFR) Basement Membrane Matrix | Corning | Cat# 354230 |
| Membrane Protran 0.45μm pore size nitrocellulose | Cytiva | Cat# 10600002 |
| MS grade water | Supelco | Cat# 1.15333 |
| MS-grade trypsin | Thermo Fisher Scientific | Cat# 90058 |
| Novex 4-12% polyacrylamide Tris-Glycine plus midi protein gels | Invitrogen | Cat# WXP41220BOX |
| Novex 4-12% polyacrylamide Tris-Glycine plus midi protein gels | Invitrogen | Cat# WXP41226BOX |
| Novex 4-20% polyacrylamide Tris-Glycine gels | Invitrogen | Cat# XP04205BOX |
| Novex Tris-Glycine SDS Running buffer | Invitrogen | Cat# LC2675 |
| NuPAGE 4-12% polyacrylamide Bis-Tris gels | Invitrogen | Cat# NP0336BOX |
| NuPAGE MOPS SDS Running buffer | Invitrogen | Cat# NP0001 |
| Opti-MEM™ I Reduced Serum Medium | Thermo Fisher Scientific | Cat# 31985047 |
| PEG4000 | Thermo Fisher Scientific | Cat# 11463207 |
| Penicillin/Streptomycin | Gibco | Cat# 15140122 |
| Phalloidin FITC conjugate | Thermo Fisher Scientific | Cat# F432 |
| Phenol-free DMEM | Thermo Fisher Scientific | Cat# 31053 |
| Polybrene | Sigma-Aldrich | Cat# TR-1003 |
| Protein A/G magnetic beads | Pierce | Cat# 88802 |
| Puromycin | Sigma-Aldrich | Cat# P7255 |
| Sorensen's Phosphate Buffer | Electron Microscopy Sciences | Cat# 11600 |
| Streptavidin FITC conjugate | Thermo Fisher Scientific | Cat#11-4317-87 |
| SuperSignal West Pico PLUS Chemiluminescent Substrate | Thermo Fisher Scientific | Cat# 15659364 |
| Ultra-low attachment plates | Corning | Cat# 7007 |
| V5-Trap Magnetic Agarose beads | Proteintech | Cat# v5ta |
| Y27632 (ROCK inhibitor) | Sigma-Aldrich | Cat# Y0503 |
| **Critical commercial assays** |  |  |
| Gateway BP Clonase | Thermo Fisher Scientific | Cat# 11789020 |
| Gateway LR Clonase | Thermo Fisher Scientific | Cat# 11791020 |
| KOD Hot Start Master Mix | Sigma-Aldrich | Cat# 71842-3 |
| qScript Cdna SuperMix | Quantabio | Cat# 95048 |
| RNeasy kit | Qiagen | Cat# 74106 |
| SYBR green select master mix | Applied Biosystems | Cat# 4472920 |
| **Deposited data** |  |  |
| Mass spectrometry raw data | This paper | See Data Availability |
| AlphaFold v3 models | This paper | See Data Availability |
| **Experimental models: Cell lines** |  |  |
| Human: Human Embryonic Kidney-293FT (HEK293FT) | Clontech | N/A |
| Mouse: PDAC cells (KPC model in C57/BL6J background) | Supp Ref 1 | N/A |
| Mouse: PDAC cells stably co-expressing mApple-FAM134B and RTN4-EGFP | This study | N/A |
| Mouse: PDAC cells stably co-expressing mApple-PRKAR1A and FAM134B-EGFP | This study | N/A |
| Mouse: PDAC cells stably expressing CALCOCO1-TID-V5 | This study | N/A |
| Mouse: PDAC cells stably expressing Cas9 | This study | N/A |
| Mouse: PDAC cells stably expressing Cas9 and FAM134B-SRAI | This study | N/A |
| Mouse: PDAC cells stably expressing Cas9 and FAM134B-V5 | This study | N/A |
| Mouse: PDAC cells stably expressing Cas9 and FAM134C-SRAI | This study | N/A |
| Mouse: PDAC cells stably expressing Cas9 and FAM134C-V5 | This study | N/A |
| Mouse: PDAC cells stably expressing Cas9 and PRKAR1A mtPS-V5 | This study | N/A |
| Mouse: PDAC cells stably expressing Cas9 and PRKAR1A ΔIDR-V5 | This study | N/A |
| Mouse: PDAC cells stably expressing Cas9 and PRKAR1A-V5 | This study | N/A |
| Mouse: PDAC cells stably expressing Cas9 and SEC62-TID-V5 | This study | N/A |
| Mouse: PDAC cells stably expressing Cas9 and SRAI-FAM134B | This study | N/A |
| Mouse: PDAC cells stably expressing Cas9 and SRAI-FAM134C | This study | N/A |
| Mouse: PDAC cells stably expressing Cas9 $\Delta Akap11$ | This study | N/A |
| Mouse: PDAC cells stably expressing Cas9 $\Delta Fam134b/c$ | This study | N/A |
| Mouse: PDAC cells stably expressing Cas9 $\Delta Prkar1a$ | This study | N/A |
| Mouse: PDAC cells stably expressing ERM-TID-V5 | This study | N/A |
| Mouse: PDAC cells stably expressing FAM134B mtLIR -V5 | This study | N/A |
| Mouse: PDAC cells stably expressing FAM134B-TID-V5 | This study | N/A |
| Mouse: PDAC cells stably expressing FAM134B-V5 | This study | N/A |
| Mouse: PDAC cells stably expressing FAM134C-TID-V5 | This study | N/A |
| Mouse: PDAC cells stably expressing FAM134C-V5 | This study | N/A |
| Mouse: PDAC cells stably expressing Luciferase-V5 | This study | N/A |
| Mouse: PDAC cells stably expressing mApple-FAM134B | This study | N/A |
| Mouse: PDAC cells stably expressing mRFP-EGFP-FAM134B | This study | N/A |
| Mouse: PDAC cells stably expressing mRFP-EGFP-RHOA | This study | N/A |
| Mouse: PDAC cells stably expressing mRFP-EGFP-RHOA G14V | This study | N/A |
| Mouse: PDAC cells stably expressing mRFP-EGFP-RHOA T19N | This study | N/A |
| Mouse: PDAC cells stably expressing TEX264-TID-V5 | This study | N/A |
| Mouse: PDAC cells stably expressing TID-V5-CALCOCO1 | This study | N/A |
| Mouse: PDAC cells stably expressing TID-V5-FAM134B | This study | N/A |
| Mouse: PDAC cells stably expressing TID-V5-FAM134C | This study | N/A |
| Mouse: PDAC cells stably expressing TID-V5-SEC62 | This study | N/A |
| Mouse: PDAC cells stably expressing V5 | This study | N/A |
| Mouse: PDAC cells stably expressing V5-TID-NES | This study | N/A |
| Mouse: PDAC $\Delta Akap11$ and stably expressing FAM134B-V5 | This study | N/A |
| Mouse: PDAC $\Delta Akap11$ and stably expressing V5 | This study | N/A |
| Mouse: PDAC $\Delta Fam134b/c$ and stably expressing FAM134B 242* -V5 | This study | N/A |
| Mouse: PDAC $\Delta Fam134b/c$ and stably expressing FAM134B 267* -V5 | This study | N/A |
| Mouse: PDAC $\Delta Fam134b/c$ and stably expressing FAM134B 292* -V5 | This study | N/A |
| Mouse: PDAC ΔFam134b/c and stably expressing FAM134B 317* -V5 | This study | N/A |
| Mouse: PDAC ΔFam134b/c and stably expressing FAM134B 342* -V5 | This study | N/A |
| Mouse: PDAC ΔFam134b/c and stably expressing FAM134B 367* -V5 | This study | N/A |
| Mouse: PDAC ΔFam134b/c and stably expressing FAM134B 392* -V5 | This study | N/A |
| Mouse: PDAC ΔFam134b/c and stably expressing FAM134B 424* -V5 | This study | N/A |
| Mouse: PDAC ΔFam134b/c and stably expressing FAM134B 449* -V5 | This study | N/A |
| Mouse: PDAC ΔFam134b/c and stably expressing FAM134B I400R -V5 | This study | N/A |
| Mouse: PDAC ΔFam134b/c and stably expressing FAM134B mtLIR -V5 | This study | N/A |
| Mouse: PDAC ΔFam134b/c and stably expressing FAM134B Δ392-408 -V5 | This study | N/A |
| Mouse: PDAC ΔFam134b/c and stably expressing FAM134B Δ392-424 -V5 | This study | N/A |
| Mouse: PDAC ΔFam134b/c and stably expressing FAM134B Δ409-424 -V5 | This study | N/A |
| Mouse: PDAC ΔFam134b/c and stably expressing FAM134B ΔCT -V5 | This study | N/A |
| Mouse: PDAC ΔFam134b/c and stably expressing FAM134B ΔINT -V5 | This study | N/A |
| Mouse: PDAC ΔFam134b/c and stably expressing FAM134B ΔNT -V5 | This study | N/A |
| Mouse: PDAC ΔFam134b/c and stably expressing FAM134B-V5 | This study | N/A |
| Mouse: PDAC ΔFam134b/c and stably expressing FAM134C-V5 | This study | N/A |
| Mouse: PDAC ΔFam134b/c and stably expressing V5 | This study | N/A |
| Mouse: PDAC ΔPrkar1a and stably expressing FAM134B-V5 | This study | N/A |
| Mouse: PDAC ΔPrkar1a and stably expressing FAM134C-V5 | This study | N/A |
| Mouse: PDAC ΔPrkar1a and stably expressing V5 | This study | N/A |
| **Oligonucleotides** |  |  |
| For gRNAs sequences see Table 1 |  |  |
| For qRT-PCR primers see Table 2 |  |  |
| **Recombinant DNA** |  |  |
| Lenti SpCas9 hygro | Supp Ref 2 | Addgene Plasmid #104995 |
| pcDNA DEST-V5<br>Gateway destination vector for V5 C terminal tagging | This paper | N/A |
| pcDNA *Fam134b* -V5<br>Mouse *Fam134b* tagged to V5 | This paper | N/A |
| pcDNA *Fam134c* -V5<br>Mouse *Fam134c* tagged to V5 | This paper | N/A |
| pcDNA FLAG DEST<br>Gateway destination vector for N terminal tagging of FLAG | This paper | N/A |
| pcDNA FLAG-*RHOA*<br>Human *RHOA* fused to N terminal FLAG | This paper | N/A |
| pcDNA FLAG-*RHOA* G14V<br>Human *RHOA* (G14V) fused to N terminal<br>FLAG | This paper | N/A |
| pcDNA FLAG-*RHOA* T19N<br>Human *RHOA* (T19N) fused to N terminal<br>FLAG | This paper | N/A |
| pCR-Blunt II-TOPO gRNA_Cloning vector | Supp Ref 3 | Addgene Plasmid #41824 |
| pDEST15 DEST<br>Gateway destination vector for N terminal<br>tagging of GST | Thermo Fisher Scientific | Cat#11802014 |
| pDEST15 GST- *Fam134b* CT<br>Mouse *Fam134b* (218-480) | This paper | N/A |
| pDEST15-GST (empty vector) | Supp Ref 4 | N/A |
| pDEST17 DEST<br>Gateway destination vector for N terminal<br>tagging of His | Thermo Fisher Scientific | Cat#11803012 |
| pDEST17 His-*PRKAR1A* NT+PS<br>Human *PRKAR1A* (1-102) | This paper | N/A |
| pDONR221 *RTN4*<br>Human *RTN4* | DNASU plasmid<br>repository (The<br>ORFeome<br>Collaboration) | Cat# HsCD00081743 |
| pDONR223 *Calcoco1*<br>Mouse *Calcoco1* | Source Bioscience | Cat# IRAVp968B12130D |
| pDONR223 *Ccpg1*<br>Mouse *Ccpg1* | Source Bioscience | Cat# IRAVp968H01136D |
| pDONR223 *Fam134b*<br>Mouse *Fam134b* | Source Bioscience | Cat# IMAGp998H0714447Q<br>and IRAVp968B0235D |
| pDONR223 *Fam134b* 242*<br>Mouse *Fam134b* (1-242) | This paper | N/A |
| pDONR223 *Fam134b* 267*<br>Mouse *Fam134b* (1-267) | This paper | N/A |
| pDONR223 *Fam134b* 292*<br>Mouse *Fam134b* (1-292) | This paper | N/A |
| pDONR223 *Fam134b* 317*<br>Mouse *Fam134b* (1-317) | This paper | N/A |
| pDONR223 *Fam134b* 342*<br>Mouse *Fam134b* (1-342) | This paper | N/A |
| pDONR223 *Fam134b* 367*<br>Mouse *Fam134b* (1-367) | This paper | N/A |
| pDONR223 *Fam134b* 392*<br>Mouse *Fam134b* (1-392) | This paper | N/A |
| pDONR223 *Fam134b* 424*<br>Mouse *Fam134b* (1-424) | This paper | N/A |
| pDONR223 *Fam134b* 449*<br>Mouse *Fam134b* (1-449) | This paper | N/A |
| pDONR223 *Fam134b* CT<br>Mouse *Fam134b* (218-480) | This paper | N/A |
| pDONR223 *Fam134b* I400R<br>Mouse *Fam134b* (I400R)/*PRKAR1A*-binding<br>deficient | This paper | N/A |
| pDONR223 *Fam134b* mtLIR<br>Mouse *Fam134b* (436-441A<sub>6</sub>) | This paper | N/A |
| pDONR223 *Fam134b* Δ392-408<br>Mouse *Fam134b* (Δ392-408) | This paper | N/A |
| pDONR223 *Fam134b* Δ392-424<br>Mouse *Fam134b* (Δ392-424) | This paper | N/A |
| pDONR223 *Fam134b* Δ409-424<br>Mouse *Fam134b* (Δ409-424) | This paper | N/A |
| pDONR223 *Fam134b* ΔCT<br>Mouse *Fam134b* (1-217) | This paper | N/A |
| pDONR223 *Fam134b* ΔINT<br>Mouse *Fam134b* (Δ123-191) | This paper | N/A |
| pDONR223 *Fam134b* ΔNT<br>Mouse *Fam134b* (43-480) | This paper | N/A |
| pDONR223 *Fam134c*<br>Mouse *Fam134c* | Source Bioscience | Cat# IRAVp968B0736D |
| pDONR223 *Luciferase*-V5<br>Firefly *Luciferase* cDNA fused to V5 | This paper | N/A |
| pDONR223 *PRKAR1A*<br>Human *PRKAR1A* | DNASU plasmid repository (The ORFeome Collaboration) | Cat# HsCD00511007 |
| pDONR223 *PRKAR1A* (gRNA insensitive)<br>Human *PRKAR1A* (gRNA insensitive) | This paper | N/A |
| pDONR223 *PRKAR1A* *mtPS*<br>Human *PRKAR1A* (R96A R97A) | This paper | N/A |
| pDONR223 *PRKAR1A* *mtPS*<br>Human *PRKAR1A* (R96A R97A) (gRNA insensitive) | This paper | N/A |
| pDONR223 *PRKAR1A* NT<br>Human *PRKAR1A* (1-93) | This paper | N/A |
| pDONR223 *PRKAR1A* NT+PS<br>Human *PRKAR1A* (1-102) | This paper | N/A |
| pDONR223 *PRKAR1A* ΔC2<br>Human *PRKAR1A* (1-240) | This paper | N/A |
| pDONR223 *PRKAR1A* ΔIDR<br>Human *PRKAR1A* (Δ62-113) (gRNA insensitive) fused to V5 | This paper | N/A |
| pDONR223 *PRKAR1A* ΔNT<br>Human *PRKAR1A* (94-381) | This paper | N/A |
| pDONR223 *PRKAR1A* ΔNT ΔPS<br>Human *PRKAR1A* (103-381) | This paper | N/A |
| pDONR223 *RHOA*<br>Human *RHOA* | This paper | N/A |
| pDONR223 *RHOA* G14V<br>Human *RHOA* (G14V) | This paper | N/A |
| pDONR223 *RHOA* T19N<br>Human *RHOA* (T19N) | This paper | N/A |
| pDONR223 *Sec62*<br>Mouse *Sec62* | Source Bioscience | Cat# IRAVp968D06104D |
| pDONR223 *Tex264*<br>Mouse *Tex264* | Source Bioscience | Cat# IRAVp968E065D |
| pEGFP<br>GFP only control | This paper | N/A |
| pEGFP DEST C1<br>Gateway destination vector for N terminal tagging of EGFP | Supp Ref 5 | N/A |
| pEGFP *PRKAR1A* (FL)<br>Human *PRKAR1A* fused to N terminal GFP | This paper | N/A |
| pEGFP *PRKAR1A* *mtPS*<br>Human *PRKAR1A* (R96A R97A) fused to N terminal GFP | This paper | N/A |
| pEGFP *PRKAR1A* NT<br>Human *PRKAR1A* (1-93) fused to N terminal GFP | This paper | N/A |
| pEGFP *PRKAR1A* NT+PS<br>Human *PRKAR1A* (1-102) fused to N terminal GFP | This paper | N/A |
| pEGFP *PRKAR1A* ΔC2<br>Human *PRKAR1A* (1-240) fused to N terminal GFP | This paper | N/A |
| pEGFP *PRKAR1A* ΔIDR<br>Human *PRKAR1A* (Δ62-113) fused to N terminal GFP | This paper | N/A |
| pEGFP *PRKAR1A* ΔNT<br>Human *PRKAR1A* (94-381) fused to N terminal GFP | This paper | N/A |
| pEGFP *PRKAR1A* ΔNT ΔPS<br>Human *PRKAR1A* (103-381) fused to N terminal GFP | This paper | N/A |
| pLIX 403 TRE-mApple DEST vector (Dox inducible)<br>Gateway destination lentiviral vector for inducible expression with N-terminal tagging of mApple | This paper, modified from Addgene plasmid #41395 (a gift from David Root) | N/A |
| pLIX 403 TRE-mApple-*Fam134b*<br>Mouse *Fam134b* fused to N-terminal mApple (Dox inducible) | This paper | N/A |
| pLIX 403 TRE-mApple-*PRKAR1A*<br>Human *PRKAR1A* fused to N-terminal mApple (Dox inducible) | This paper | N/A |
| pLX304 *Calcoco1*-TID-V5<br>Mouse *Calcoco1* fused to C-terminal TID-V5 | This paper | N/A |
| pLX304 EGFP N1 DEST<br>Gateway destination lentiviral vector for C-terminal tagging of EGFP | This paper | N/A |
| pLX304 ERM-TID-V5<br>ERM=N-terminal sequence C1(1-29) of rabbit *CYP2C1* fused to TID-V5 | Supp Ref 6 | Addgene Plasmid #107175 |
| pLX304 EV (-)<br>Empty lentiviral vector | This paper | N/A |
| pLX304 *Fam134b* 242*-V5<br>Mouse *Fam134b* (1-242) fused to V5 | This paper | N/A |
| pLX304 *Fam134b* 267*-V5<br>Mouse *Fam134b* (1-267) fused to V5 | This paper | N/A |
| pLX304 *Fam134b* 292*-V5<br>Mouse *Fam134b* (1-292) fused to V5 | This paper | N/A |
| pLX304 *Fam134b* 317*-V5<br>Mouse *Fam134b* (1-317) fused to V5 | This paper | N/A |
| pLX304 *Fam134b* 342*-V5<br>Mouse *Fam134b* (1-342) fused to V5 | This paper | N/A |
| pLX304 *Fam134b* 367*-V5<br>Mouse *Fam134b* (1-367) fused to V5 | This paper | N/A |
| pLX304 *Fam134b* 392*-V5<br>Mouse *Fam134b* (1-392) fused to V5 | This paper | N/A |
| pLX304 *Fam134b* 424*-V5<br>Mouse *Fam134b* (1-424) fused to V5 | This paper | N/A |
| pLX304 *Fam134b* 449*-V5<br>Mouse *Fam134b* (1-449) fused to V5 | This paper | N/A |
| pLX304 *Fam134b* I400R-V5<br>Mouse *Fam134b* (I400R) fused to V5 | This paper | N/A |
| pLX304 *Fam134b* mtLIR-V5<br>Mouse *Fam134b* (436-441A<sub>6</sub>) fused to V5 | This paper | N/A |
| pLX304 *Fam134b* Δ392-408-V5<br>Mouse *Fam134b* (Δ392-408) fused to V5 | This paper | N/A |
| pLX304 *Fam134b* Δ392-424-V5<br>Mouse *Fam134b* (Δ392-424) fused to V5 | This paper | N/A |
| pLX304 *Fam134b* $\Delta 409-424$-V5<br>Mouse *Fam134b* ($\Delta 409-424$) fused to V5 | This paper | N/A |
| pLX304 *Fam134b* $\Delta CT$-V5<br>Mouse *Fam134b* (1-217) fused to V5 | This paper | N/A |
| pLX304 *Fam134b* $\Delta INT$-V5<br>Mouse *Fam134b* ($\Delta 123-191$) fused to V5 | This paper | N/A |
| pLX304 *Fam134b* $\Delta NT$-V5<br>Mouse *Fam134b* fused to V5 | This paper | N/A |
| pLX304 *Fam134b*-EGFP<br>Mouse *Fam134b* fused to C terminal EGFP | This paper | N/A |
| pLX304 *Fam134b*-SRAI<br>Mouse *Fam134b* fused to C terminal SRAI | This paper | N/A |
| pLX304 *Fam134b*-TID-V5<br>Mouse *Fam134b* fused to C terminal TID-V5 | This paper | N/A |
| pLX304 *Fam134b*-V5<br>Mouse *Fam134b* fused to V5 | This paper | N/A |
| pLX304 *Fam134c*-SRAI<br>Mouse *Fam134c* fused to C terminal SRAI | This paper | N/A |
| pLX304 *Fam134c*-TID-V5<br>Mouse *Fam134c* fused to C terminal TID-V5 | This paper | N/A |
| pLX304 *Fam134c*-V5<br>Mouse *Fam134c* fused to V5 | This paper | N/A |
| pLX304 *Luciferase*-V5<br>Firefly *Luciferase* cDNA fused to V5 | This paper | N/A |
| pLX304 mRFP-EGFP-DEST-C1<br>Gateway destination lentiviral vector for N terminal tagging of mRFP-EGFP | This paper | N/A |
| pLX304 mRFP-EGFP-*Fam134b*<br>Mouse *Fam134b* fused to N terminal mRFP-EGFP | This paper | N/A |
| pLX304 mRFP-EGFP-*RHOA*<br>Human *RHOA* fused to N terminal mRFP-EGFP | This paper | N/A |
| pLX304 mRFP-EGFP-*RHOA T19N*<br>Human *RHOA* (T19N) fused to N terminal mRFP-EGFP | This paper | N/A |
| pLX304 mRFP-EGFP-*RHOA G14V*<br>Human *RHOA* (G14V) fused to N terminal mRFP-EGFP | This paper | N/A |
| pLX304 *PRKAR1A* (FL)-V5<br>Human *PRKAR1A* (gRNA insensitive) fused to V5 | This paper | N/A |
| pLX304 *PRKAR1A mtPS* -V5<br>Human *PRKAR1A* (R96A R97A) (gRNA insensitive) fused to V5 | This paper | N/A |
| pLX304 *PRKAR1A* $\Delta IDR$-V5<br>Human *PRKAR1A* ($\Delta 62-113$) (gRNA insensitive) fused to V5 | This paper | N/A |
| pLX304 *RTN4*-EGFP<br>Human *RTN4* fused to C terminal EGFP | This paper | N/A |
| pLX304 *Sec62*-TID-V5<br>Mouse *Sec62* fused to C terminal TID-V5 | This paper | N/A |
| pLX304 *SRAI-C1 DEST*<br>Gateway destination vector | Supp Ref 7 | N/A |
| pLX304 *SRAI-Fam134b*<br>Mouse *Fam134b* fused to N terminal SRAI | This paper | N/A |
| pLX304 *SRAI-Fam134c*<br>Mouse *Fam134c* fused to N terminal SRAI | This paper | N/A |
| pLX304 *SRAI-N1 DEST*<br>Gateway destination vector | Supp Ref 7 | N/A |
| pLX304 *Tex264*-TID-V5<br>Mouse *Tex264* fused to C terminal TID-V5 | This paper | N/A |
| pLX304 TID-C1 DEST<br>Gateway destination vector | This paper | N/A |
| pLX304 TID-N1 DEST<br>Gateway destination vector<br>TID= TurboID (BirA mutant) | This paper based on<br>Supp Ref 6 | N/A |
| pLX304 TID-V5- *Calcoco1*<br>Mouse *Calcoco1* fused to N terminal TID-V5 | This paper | N/A |
| pLX304 TID-V5- *Ccpg1*<br>Mouse *Ccpg1* fused to N terminal TID-V5 | This paper | N/A |
| pLX304 TID-V5- *Fam134b*<br>Mouse *Fam134b* fused to N terminal TID-V5 | This paper | N/A |
| pLX304 TID-V5- *Fam134c*<br>Mouse *Fam134c* fused to N terminal TID-V5 | This paper | N/A |
| pLX304 TID-V5- *Sec62*<br>Mouse *Sec62* fused to N terminal TID-V5 | This paper | N/A |
| pLX304 V5 DEST<br>Gateway destination vector | Supp Ref 8 | Addgene plasmid #25890 |
| pLX304 V5-TID-NES<br>NES = nuclear export signal from the HIV *Rev* protein fused in the N terminus to V5-TID | This paper | N/A |
| pMD2.G | A gift from Didier Trono | Addgene Plasmid #12259 |
| psPAX2 | A gift from Didier Trono | Addgene Plasmid #12260 |
| pSpCas9(BB)-2A-Puro (PX459) V2.0 | Supp Ref 9 | Addgene Plasmid #62988 |
| **Software and algorithms** |  |  |
| AlphaFold v3 | Supp Ref 10 | https://www.alphafoldserver.com |
| Cytoscape v3.10 | Supp Ref 11 | https://cytoscape.org |
| Excel | Microsoft | https://www.microsoft.com/en-us/microsoft-365/excel |
| FoldX 5.0 suite | Supp Ref 12 | https://foldxsuite.crg.eu |
| GraphPad v10.6 | GraphPad Software | https://www.graphpad.com |
| ImageJ 2.9.0 | NIH, Supp Ref 13 | https://imagej.net/ij |
| Imaris 10.0.1 | Oxford instruments | https://imaris.oxinst.com/ |
| IncuCyte S3 | Satorius | https://www.sartorius.com/en/products/live-cell-imaging-analysis/live-cell-analysis-software |
| LAS X software | Leica Microsystems UK | https://www.leica-microsystems.com/products/microscope-software/p/leica-las-x-ls |
| MSFragger v. 22.0 | Supp Ref 14 | https://msfragger.nesvilab.org |
| Nikon NIS-Elements Advanced Research | Nikon | https://www.microscope.healthcare.nikon.com/en_EU/products/software/nis-elements/nis-elements-advanced-research |
| QuPath 0.5.0 | Supp Ref 15 | https://qupath.github.io/ |
| STRING v12.0 | Supp Ref 16 | https://string-db.org |

### Tissue culture

A list of all cell lines can be found in “Experimental models: Cell lines” in Key Resource Table. All lines were cultured in full medium (high glucose Dulbecco’s modified Eagle’s medium (DMEM) supplemented with 10% Fetal Bovine Serum (FBS), 2mM L-Glutamine and 10U/ml Penicillin/Streptomycin) at 37 °C in 5 % CO_2_ unless indicated otherwise. Parental cell lines were HEK293FT human embryonic kidney cell line, from laboratory stocks, and pancreatic ductal adenocarcinoma (PDAC) KPC mouse cell line (C57/BL6J), obtained from Owen Sansom (Beatson Institute for Cancer Research, Glasgow, UK) ^1^. For derivative stable cell pool and clone generation, see section “Viruses and stable cell lines”. Amino acid starvations were performed by washing the cells three times in phosphate-buffered saline (PBS) before culture in fresh Earle’s balanced salt solution (EBSS) for 4 h. Cells were tested every two months to confirm the absence of mycoplasma contamination.

### Plasmids and transfections

The majority of cloning was performed by using the Gateway method following manufacturer’s instructions (i.e., LR and BP gateway reactions, Thermo Fisher Scientific). Truncations and single point mutations were generated by PCR using the KOD polymerase as per manufacturer’s instructions (Sigma-Aldrich, 71842-3). Detailed sequence maps are available from authors upon request. Transient transfections were performed with Lipofectamine 2000 (Thermo Fisher Scientific) following manufacturer’s recommendations with a ratio DNA : Lipofectamine 2000 of 1:3 in Opti-MEM media.

### Viruses and stable cell lines

Lentivirus (from plasmids pLX304, pLIX and lentiCas9) were produced in HEK293FT cells after transient transfection with packaging plasmids as previously described ^7^. Viruses were collected in high-serum full medium (30% FBS) 48 h post-transfection. Media were then filtered using a 0.45 µm filter.

Stable cell lines were generated by transducing PDAC cells with the corresponding lentivirus in the presence of 10 µg/mL polybrene. After 48h, cells were fed with media supplemented with their corresponding antibiotic: for lentiCas9, 1 mg/mL hygromycin B (10 days); for pLX304 plasmids, 10 µg/mL blasticidin (4 days); for pLIX plasmids, 3 µg/mL puromycin (2 days). Prior to experimentation, PDAC cells stably transduced with pLIX mApple constructs were treated with 0.5 µg/mL doxycycline (Bio-Techne) for 24 h to induce the expression of the construct.

### Proximity biotinylation and proteomics

Proximity labelling was performed in PDAC cell pools stably expressing the ER-phagy receptors FAM134B, FAM134C, SEC62, CALCOCO1 and TEX264 with N-terminal or C-terminal tagging of TID-V5. As controls, stable cell lines expressing Luciferase-V5, ERM-TID-V5 (ERM=N-terminal sequence C1(1-29) of rabbit CYP2C1) and V5-TID-NES (NES= nuclear export signal from the HIV Rev) were used ^6^. 3 x 10^6^ cells were plated on 10 cm dishes in triplicate and incubated at 37°C at 5% CO_2_. Alternatively, for immunofluorescence characterisation, 1 x 10^5^ cells were seeded on coverslips. The following day, cells were incubated with Biotin (0.5 mM) for 1 h (FAM134B-TID-V5, FAM13C-TID-V5, SEC62-TID-V5, TEX264-TID-V5, TID-V5-CALCOCO1, TID-V5-FAM134B, TID-V5-FAM134C, TID-V5-SEC62); or 4 h (Luciferase-V5, ERM-TID-V5, V5-

TID-NES, TID-V5-CCPG1, CALCOCO1-TID-V5) at 37°C at 5% CO_2_. For Forskolin (FSK)-treated FAM134B proximity labelling proteomics, the corresponding treatment, i.e., DMSO or FSK (40 µM, 4 h), was added alongside the Biotin (0.5 mM) for 4 h (Luciferase-V5, ERM-TID-V5, V5- TID-NES, TID-V5-FAM134B) or 2 h (FAM134B-TID-V5). Cells were rinsed five times with ice-cold PBS, and then scraped into 1.5 mL tubes with 1 mL ice-cold PBS. Cells were then pelleted by centrifugation at 300 g for 3 min and resuspended in 500 µl of RIPA buffer (50 mM Tris-HCl pH 7.5, 0.2% SDS, 150 mM NaCl, 0.5 % Sodium Deoxycholate, 2 mM activated sodium orthovanadate, 20 mM sodium fluoride, 10 mM sodium pyrophosphate, 1 mM PMSF and Complete Protease Inhibitor (Sigma-Aldrich, 11697498001)). Samples were incubated on ice for 10 min. Lysates were clarified by centrifugation at 13,000 g at 4°C for 10 min and 2% (1/50) of the sample was kept as input. Input samples were boiled at 95°C for 5 min and subjected to SDS-PAGE and immunoblotting. Cleared lysates were incubated with streptavidin-linked sepharose beads (Dynabeads MyOne Streptavidin C1, Thermo Fisher, 10 µl per reaction; previously washed thrice with RIPA lysis buffer), for 2 h at 4 °C with agitation.

For mass spectrometry, samples were then processed for digestion using a KingFisher Duo and Flex systems (Thermo Fisher Scientific). The plates (rows A–H) were arranged according to the order of the washes, starting from row H. Tips were placed in row H, and lysates were placed in row G. Sequential washes (2 min/each) were two RIPA buffer washes, one 1 M KCl wash, one 0.1 M Na₂CO₃ wash, and one 2 M urea in 10 mM Tris-HCl (pH 8) wash. Then, samples were incubated with digestion buffer consisting of 2 M urea, 1 mM dithiothreitol (DTT), and 0.1 μg/μL MS-grade trypsin. Controlled on-bead digestion was performed for 30 min at 37 °C, after which the beads were removed, and the supernatant was digested overnight at 37 °C. Following digestion, samples were alkylated with 20 μL of 5 mg/mL iodoacetamide for 30 min in the dark at room temperature. Samples were then acidified with 10% trifluoroacetic acid (TFA) to a final concentration of 1%. Peptides were desalted using C18 filters in P200 tips, activated with methanol and 0.1% TFA, then eluted with 50% acetonitrile (ACN) containing 0.05% TFA. Eluates were dried using a vacuum centrifuge and reconstituted in 0.1% TFA. 5 µl of the resuspended peptides were analyzed using reversed-phase nano-LC-MS/MS with a nano-Ultimate 3000 LC system and a Lumos Fusion mass spectrometer (Thermo Fisher Scientific). Flow rates were 400 nl/min. Peptides were loaded onto 25 cm Aurora Columns (IonOptiks) utilizing a 67-minute gradient with buffer A (2% acetonitrile, 0.5% acetic acid) and buffer B (80% acetonitrile, 0.5% acetic acid); from 0 to 16 min: 2% buffer B, 16 to 56 min: 3 to 35% buffer B, 56 to 62 min: 99% buffer B; and from 62 to 67 min: 2% buffer B. Full-scan spectra in the Orbitrap were recorded within the m/z range of 350 to 1,400 (resolution: 240,000; AGC: 7.5e5 ions). MS2 was performed in the ion trap, with an isolation window of 0.7, an AGC of 2e4, an HCD collision energy of 28, rapid scan rate, a scan range of 145 to 1450 m/z, a maximum injection time of 50 ms, and an overall cycle time of 1 s. Samples were searched with the appropriate FASTA database and modifications using MS-Fragger v. 22.0.

The protein quantification was obtained from the proteinGroups (Fig. 1) or combined protein (Fig. 6) files obtained after the MSFragger search. After removal of contaminants and single peptides, the file was uploaded to R Studio and processed by utilising a readily available Limma pipeline on GitHub ^17^ for two-group comparisons of the TID-bait with different controls. Identified proteins were put through a cut-off filter using -log *P* value > 1.3 (i.e., *P* <0.05) vs. Luciferase-V5 and TID-NES (or ERM-TID for cytoplasmic CALCOCO1). Remaining proteins were assessed for enrichment vs. ERM-TID (or TID-NES for CALCOCO1) as follows:

For the first set of proximity labelling proteomics (Table S1), high-confidence candidates were chosen with the following criteria - FAM134B-TID-V5, log2(fold change) of > 3.5 and -log *P* value > 1.3 (i.e., P <0.05); TID-V5-FAM134B log2(fold change) of > 4.5 and -log *P* value > 1.25 FAM134C-TID-V5 log2(fold change) of > 2.5 and -log *P* value > 1.3; TID-V5-FAM13C log2(fold change) of > 4 and -log *P* value > 1.3; SEC62-TID-V5 log2(fold change) of > 3.5 and -log *P* value > 1.3; TID-V5-SEC62 log2(fold change) of > 3 and -log *P* value > 1.3; TEX264-TID-V5 log2(fold change) of > 6 and -log *P* value > 1.3; TID-V5-CCPG1 log2(fold change) of > 3 and -log *P* value > 0.9; CALCOCO1-TID-V5 log2(fold change) of > 8 and -log *P* value > 1.3; and TID-V5-CALCOCO1 log2(fold change) of > 6.5 and -log P value > 1.3. Robust *Z*-scores were calculated using the robust *Z*-score method 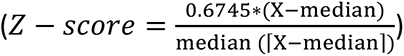 with X being log_2_(fold change).

Robust Z-score values were taken for the selected high-confidence candidates for each receptor bait. With the aim to select the highest confidence candidates independently of the terminus chosen for tagging of the receptors, for ER-phagy receptors tagged to both of their cytoplasmic terminuses (i.e., FAM134B, FAM134C, SEC62 and CALCOCO1), for a given candidate the highest robust Z-score for the candidate was selected. Values were then manually clustered for receptor specificity and presented as a heatmap using GraphPad. Absent or negative Z-score values for a given candidate in other receptor bait datasets were binned as “≤0” for the purposes of heatmap readability.

In direct immunoblot validation of streptavidin pulldowns for FAM134B/C, a higher amount of streptavidin beads was used (15 µl per reaction) and samples were incubated overnight. The following day, samples were washed manually with the same buffers using a magnetic rack at room temperature. In the last wash, beads were transferred to new tubes and beads were then resuspended in 40 µl 2x Laemmli (125 mM Tris HCl pH 6.8, 4% SDS, 20% glycerol, 10% 2-β mercaptoethanol, 0.004% bromphenol blue) supplemented with 2 mM Biotin and 20 mM DTT. Samples were boiled at 95°C for 10 min, and supernatant was subjected to SDS-PAGE and immunoblotting.

### Gene ontology and network analysis

For the second set of proximity labelling proteomics (Table S3), proteins were filtered on a basis of *P* < 0.05 for comparison of FAM134B-TID-V5 versus V5-TID-NES and then ratio of fold enrichment (ΔFC) versus ERM-TID-V5 under FSK treatment with respect to DMSO control was calculated. The greatest magnitude ΔFC were selected for ontological analysis (log2 ΔFC > 0.5, FSK-enriched preys; or < 0.5, DMSO-enriched preys). These selected hits were analysed using STRING 12.0, using default protein mapping and FDR ≤ 0.05. The ontology analysis results for Reactome pathways were rendered and exported as shown in the data figures. N.B. a similar analysis was performed for TID-V5-FAM134B but no significantly enriched Reactome pathways were identified for FSK treatment. In a second “all conditions” analysis, 344 unique hits from the combined proteomic runs (FAM134B tagging at either terminus and treatment with either DMSO or FSK), were identified using a threshold of *P* < 0.01 compared with V5- TID-NES -V5 and log2(fold change) > 1 for comparison of each bait versus ERM-TID. These were analysed using STRING v12.0, as described above.

A deeper investigation of RHO GTPase Reactome pathways was performed by examining the subnetwork of FAM134B-interacting proteins (combined from FSK-enriched and “all conditions” analyses above) belonging to the pathway MMU-9716542 (“Signaling by Rho GTPases, Miro GTPases and RHOBTB3”). Physical subnetworks (from experimental and database sources at interaction score > 0.04) were selected from within both human and mouse orthologue networks in STRING v12.0, with incorporation of additional RHOA, then networks were merged in Cytoscape and rendered to focus on a central continuous physical interaction network around RhoA.

### CRISPR/Cas9 gene targeting

Knockout PDAC cell lines were generated using CRISPR/Cas9 gene targeting. PDAC-Cas9 cells were generated by transduction of PDAC cells with lentivirus expressing SpCas9 and selected with 1 mg/ml hygromycin B for 10 days. Short guide RNAs (gRNAs) targeting the gene of interest (see Table 1) ligated into pCR-Blunt II-TOPO gRNA cloning plasmids were co-transfected using Lipofectamine 2000 (Thermo Fisher Scientific) as per manufacturer’s recommendation in a ratio 3:1 with a plasmid expressing Cas9 and puromycin resistance (pSpCas9(BB)-2A-Puro, Addgene #62988). The following day, transfected cells were selected with media supplemented with 3 μg/mL puromycin for 2 days. Cells were used 6-8 days after transfection (knockout pools). Alternatively, clones for Δ*Prkar1a*, Δ*Fam134b/c* and Δ*Atg7* were generated using limiting dilution cloning in 96-well plates.

**Table 1.**
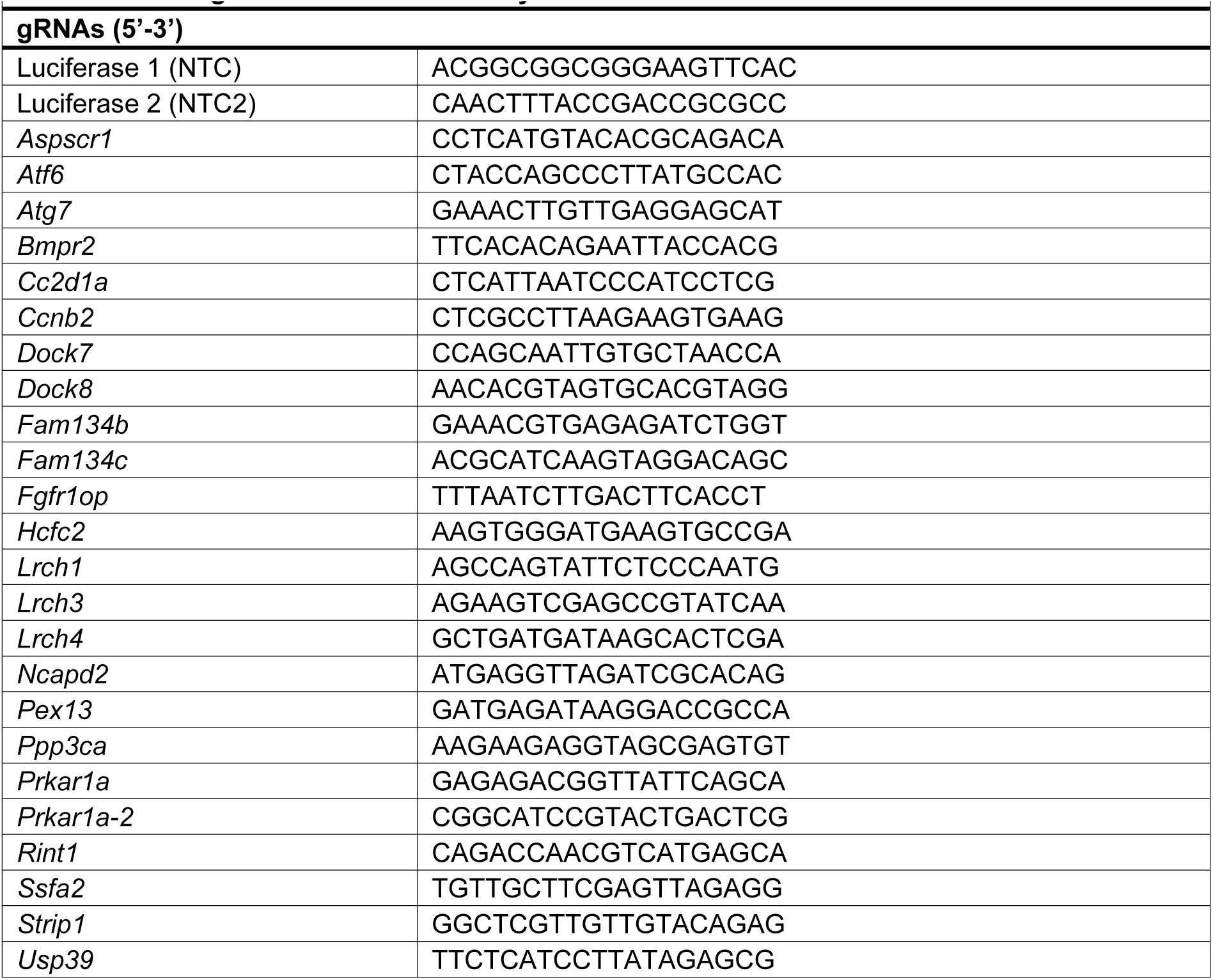
List of gRNAs used in this study.

For reconstitution of PRKAR1A for attempted rescue of transient transfected g*Prkar1a* cells, PRKAR1A cDNA was synthesized mammalian codon optimized on the gRNA-targeted region, so that g*Prkar1a* would not recognize the targeting sequence. Constructs were cloned into pLX304 plasmids for lentiviral production. Cells were transduced with PRKAR1A-V5- expressing lentivirus (i.e., PRKAR1A-V5, PRKAR1A mtPS-V5, PRKAR1A ΔIDR-V5 or V5 empty vector as a control) 1 day before cells were transfected with gNTC or g*Prkar1a*. The day after transfection, media was replaced with 3 µg / mL puromycin and 10 µg / mL blasticidin for 2 days. Cells were used 6-8 days after transfection.

### Quantitative RT-PCR

PDAC cells were plated in 12-well plates 6 days after gRNA was transfected. Total RNA was extracted through columns using RNeasy kit as per manufacturer’s instructions and genomic DNA was digested using DNaseI. RNA concentration was measured using a Nanodrop 2000 Spectrophotometer (Thermo Fisher Scientific). Then, cDNA was synthesised from total RNA using qScript cDNA SuperMix (Quantabio) according to manufacturer’s instructions. qPCR was performed on an Applied Biosystems StepOne Plus Real-Time PCR System using SYBR green select master mix (Applied Biosystems) and using the primers listed in Table 2 (Integrated DNA Technologies) as per manufacturer’s protocol. For the analysis, RNA levels were estimated using the ΔΔCt method and normalized to 18S rRNA.

**Table 2.**
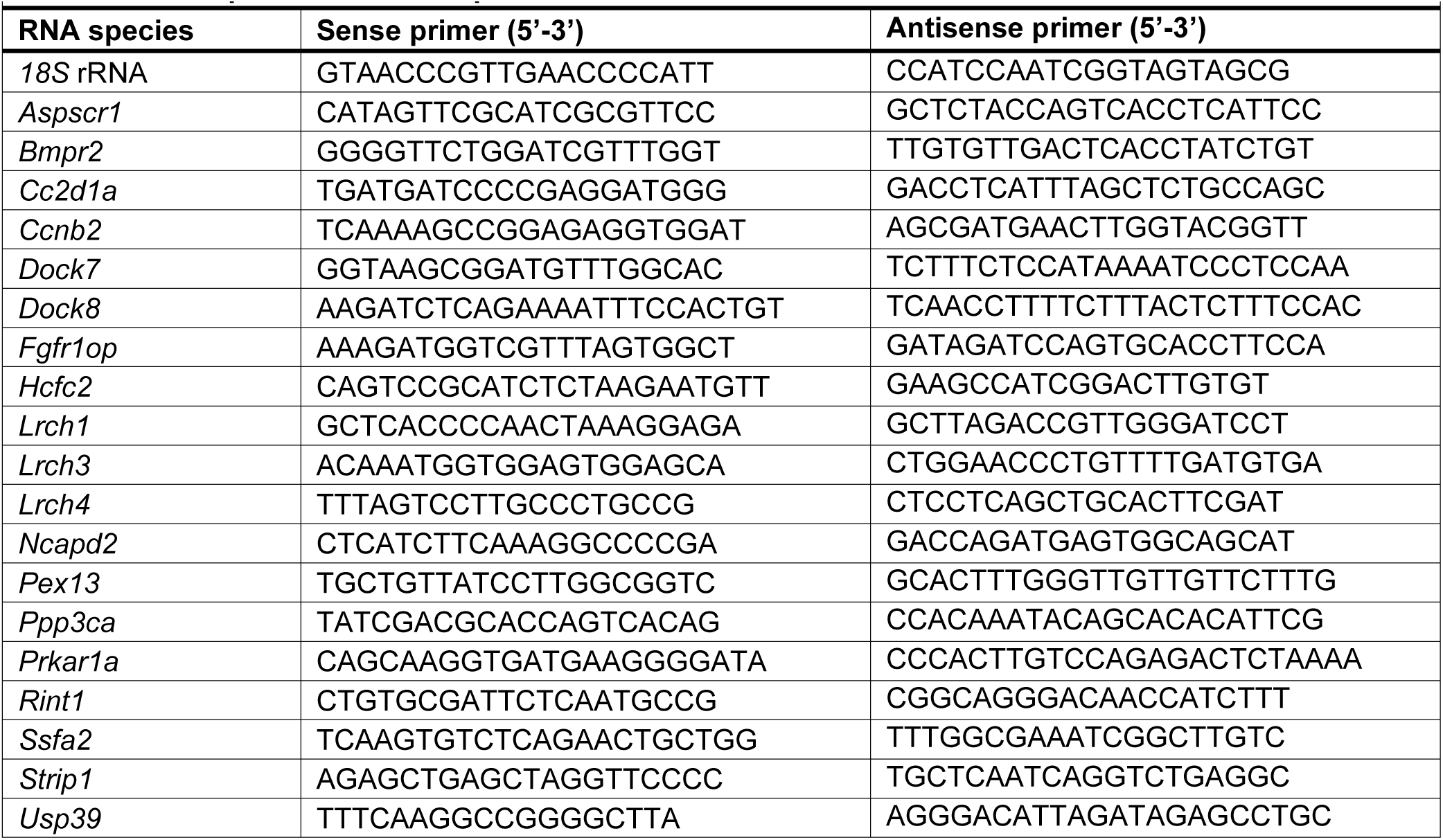
List of primers used for quantitative RT-PCR.

### SRAI flux reporter analysis

For screening, PDAC-Cas9 cells (see “Viruses and stable cell lines”) stably expressing FAM134B/C tagged to the signal retaining autophagy indicator (SRAI) probe at the C-terminus or N terminus were generated, imaged and analysed as previously reported 7. Briefly, PDAC- Cas9 were transduced with lentivirus expressing the SRAI-tagged constructs and pools of expressing cells selected using FACS. Then, cell pools were subjected to CRISPR/Cas9 gene targeting for the gene of interest (see Table 1, section “CRISPR/Cas9 gene targeting”). After 7-8 days, 20,000 cells were plated on glass bottom 96 well plates (IBL). The following day, cells were treated with the corresponding treatments (i.e. Full nutrient or starvation (EBSS, 4 h); DMSO or FSK (10 µM, 2 h)) and then fixed with 4% paraformaldehyde (PFA) in PBS, pH 7.2 at room temperature for 10 min. Cells were then washed twice with PBS and imaged with automated acquisition using a Nikon Eclipse Ti-E inverted microscope (Nikon Instruments UK Ltd., Kingston Upon Thames, UK) with a 40X NA 0.75 objective and a Nikon Nis-Elements JOBS module (script available from GitHub 18). Widefield illumination is provided by a CoolLED PE- 4000 LED light source combined with LED-CFP/YFP/mCherry-3X-A (Pinkel) filter sets. Images were acquired with a Photometrics Prime BSI sCMOS camera. For quantification, TOLLES:YPet index was obtained using a developed ImageJ macro as previously shown 7. Thresholds were empirically chosen using positive (EBSS) and negative controls (BafA1, 50 nM, 4 h). For the screening, robust z-scores were calculated using the robust Z-score method 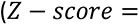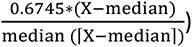 with X being the logarithmic value of the average of every sample within the biological replicate. Then, the mean of the robust Z-scores from two biological replicates with each terminal tagging were presented as a heatmap using GraphPad.

### Structural modelling and FAM134B mutant prediction

AlphaFold v3 server was used for the exploration of interactions between dimeric PRKAR1A (UniProt Q9DBC7) and FAM134B (UniProt Q8VE91). Further exploration with truncated sequences (a.a. 1-92 and 245-480, respectively) to derive more efficient structures for downstream stability evaluation software. The truncated sequences retained the same interaction patterns. The models are subject to AlphaFold Server Output Terms of Use found www.alphafoldserver.com/output-terms. The models were further explored using the FoldX 5.0 suite, which allows energetic evaluation of biomolecular complex stability and interactions ^12^. The structures were relaxed using the ‘RepairPDB’ utility, both for the complex and the extracted individual FAM134B chains. To identify the residues participating in protein-protein interactions, the ‘AnalyseComplex’ was used, which produced a list of interacting residues. Utilizing the FAM134B residue positions identified to interact with the PRKAR1A dimer, ‘BuildModel’ was used to derive stability estimates for all amino acid residue substitutions in the model, deriving values for the difference in the change of Gibbs free energy (ΔΔ*G*) for FAM134B (245-480) chain monomers and the tertiary structure in complex with PRKAR1A (1-92) dimers.

### Immunofluorescence

Cells were grown on 12 mm glass coverslips unless otherwise specified. After the corresponding treatment, cells were fixed by incubating with 4% paraformaldehyde in PBS, pH 7.2 at room temperature for 10 min. Alternatively, for cytoskeletal staining (i.e., RhoA and phosphorylated MLC2), cells were fixed with 4% paraformaldehyde in PHEM buffer, pH 6.9 (60 mM PIPES, 25 mM HEPES, 10 mM EGTA, 2 mM MgCl2) at 4°C for 15 min. Then, cells were permeabilised with either methanol, kept at -20°C for 10 minutes (LC3B) or with 0.25% Triton X-100 in PBS for 15 minutes at room temperature (all other antibodies). After washing twice with PBS, cells permeabilized with Triton-X-100 were blocked for 1 h at room temperature in 5% BSA/PBS/0.02% sodium azide, before adding the primary antibody. Then, cells were incubated with the primary antibody solution (1% BSA/PBS/0.02% sodium azide, or PBS for methanol-fixed cells) for 2 hours at room temperature. Antibody dilutions are listed in the Key Resource Table. Cells were washed three times for 5 min each in PBS, then incubated with secondary antibodies and costained with DAPI (1:2000) for 1 h at room temperature. Finally, cells were washed three times for 5 min each in PBS and mounted on glass slides using Dako fluorescent mounting medium.

### Confocal microscopy

Generally, images were captured with an Olympus FV3000 Confocal Laser Scanning Microscope using Fluoview FV31S-SW (version 2.6.1.243) Software. The microscope consists of an IX83 inverted frame and galvanometer scanhead coupled to a laser bed containing 405, 432, 488, 515, 561 and 640 nm laser lines. Images were taken with an Olympus UPlanSApo 60× 1.35 NA Oil objective. Fluorescence signals were resolved onto the detectors using a series of dichroic mirrors and filters combined with spectral gratings selected as appropriate by the software for the fluorophore. Images were taken using the sequential scan setting to reduce fluorescence bleed through. Images were further analysed in ImageJ 2.9.0 or Imaris 10.0.1 or QuPath v0.5.0.

### Spinning Disk Super-Resolution by Optical Pixel Reassignment (SoRA)

Cells were grown on HiQA Coverslips No. 1.5H, 12mm Ø (100,000 cells). For mApple-FAM134B cell line, doxycycline (0.5µg/ml, 24 h) was supplemented on the media at the time of plating. The following day, cells were then treated with the corresponding treatments (DMSO or forskolin, FSK) and then fixed with 4% paraformaldehyde in PBS. Coverslips were then subjected to immunofluorescence for the corresponding antibody staining (Section “Immunofluorescence”). Super resolution images were obtained using Nikon SoRa (Nikon Europe BV) equipped with 405 nm, 488 nm, 561 nm and 640 nm laser lines. Samples were visualised using a SR Apo TIRF 100x 1.49NA oil lens. The images were captured using PrimeBSi 95B CMOS cameras (Teledyne Photometrics). Data was captured in 3D, for this the z-step size was set to 0.1 μm, and images were acquired, denoised, and deconvolved using Nikon NIS-Elements Advanced Research software (Ver 5.30.05) as per manufacturer’s instructions. 3D surface rendering and 3D volume rendering with normal shading mode was conducted with Imaris 10.0.1 (Oxford Instruments).

### Fluorescence after photobleaching (FRAP)

PDAC stably expressing mApple-PRKAR1A and EGFP-FAM134B were seeded on µ-Slide 8 Well high Glass Bottom plates (Ibidi, 70,000 cells/well). Doxycycline (0.5 µg/ml) was added at the time of plating to induce mApple-PRKAR1A. The following day, media was replaced for complete phenol-free DMEM media and the corresponded treatment was added (DMSO or 40 µM forskolin, FSK) for 2 h. FRAP was performed on a DMi8 Stellaris 8 point scanning confocal microscope using LAS X software (Leica Microsystems UK Ltd, Milton Keynes, UK) using a 63x 1.40 NA objective lens. This microscope is equipped with a white light laser, enabling free tuning of the excitation wavelength between 440-790 nm and the detector is a Hybrid APD/GaAsP. Environmental control for live cell work was provided by an Okolab microscope enclosure with air heater and Bold Line CO_2_ unit (Okolab S.R.L, Ottaviano, NA, Italy). Image stacks were acquired with the corresponding wavelength (488 nm for EGFP- FAM134B or 561 nm for mApple-PRKAR1A) with a frame size of 2048 x 512. Images for both channels were taken prior to FRAP. Then, for each individual channel, small regions of interest (ROIs) of 3 µm for stimulation were drawn over the focal structure of double positive foci or the cytosol. For every stimulation, additional non-bleachable ROIs – outside of the cells and an unbleached cytoplasmic area – of the same size as the bleached ROI, were selected for background subtraction and data normalization. The total FRAP series contained 4 pre-bleach images (1 s interval) followed by 8 cycles of ROI bleaching with 100% laser intensity and 40 s of continuous acquisition to monitor fluorescence recovery. Fluorescence intensity of the ROIs was quantified using LAS X software and images were further analysed in ImageJ 2.9.0. For quantification or bleaching recovery, GraphPad was used to quantify mean area under curves in the first 10 s of recovery (fast recovery) and standard error, with a 0.67 threshold (lowest intensity after photobleaching). Then, a 1-sample two-tailed t-test was performed to compare to no recovery (population mean area = 0).

### Correlative light-electron microscopy (CLEM)

PDAC stably expressing mApple-PRKAR1A and FAM134B-EGFP were plated on 35mm gridded glass-bottomed imaging dishes (Mattek) (300,000 cells). At the time of plating, media was supplemented with doxycycline (0.5 µg/ml, 24 h). The following day, FSK (40 µM) was spiked for 2h. Then cells were fixed with 2% paraformaldehyde (EM grade) with 0.2% glutaraldehyde (EM grade) and 0.1 M Sorensen’s buffer (Electron Microscopy Sciences) for 20 min at room temperature. Then solution was replaced with 1% PFA paraformaldehyde in 0.1 M Sorensen’s buffer. For electron microscopic imaging of sections, following imaging on a Leica SP8 AOBS confocal laser scanning microscope using a 63x oil objective, cells were washed in 0.1 M sodium cacodylate buffer (pH 7.3) then incubated with 1% osmium tetroxide and 1.5% potassium ferricyanide in sodium cacodylate buffer for 1 h in the dark. Following 3x washes in sodium cacodylate buffer, cells were dehydrated in an EtOH series (75%; 90%; 100%), incubated for 1 h in 50% Epon in EtOH, then embedded in Epon overnight at 60°C. Following resin block trimming, 75 nm sections were cut using an Ultracut S ultramicrotome (Leica) and picked up on pioloform slot grids. Sections were stained *en face* in 2% uranyl acetate (10 min), and lead citrate (10 mins; ^19^). 15 nm gold fiducials (Aurion, TomoSol solution) were applied to both surfaces of the sections. The sections were imaged at 120 kV in a Talos L120C (Thermo Scientific) with a ceta 16M camera and tilt series images acquired between −70° to +70° at 1.5° increments.

### Immunoblotting

For direct immunoblotting analysis, cells were plated on 12-well plates and incubated at 37°C in 5 % CO_2_. The following day, after treatment if applicable, cells were initially washed with ice-cold PBS, then lysed in SDS lysis buffer (4 % SDS, 150 mM NaCl, 50 mM Tris, pH 7.5). Sample protein concentration was determined by Nanodrop (A280, Thermo Fisher Scientific). Samples were diluted with Laemmli buffer (0.3125 M Tris HCl, pH 6.8, 10% SDS, 50% glycerol, 25% 2-β mercaptoethanol, 0.02% bromphenol blue) and denatured at 95 °C for 5 min. Denatured samples were then separated by electrophoresis on MOPS 4-12% polyacrylamide Bis-Tris gels or 4-20% polyacrylamide Tris-Glycine gels (for smaller proteins, e.g., LC3B and RhoA), or Novex 4-12% polyacrylamide Tris-Glycine plus midi protein gels (if bigger sample size) following manufacturer’s instructions (Thermo Fisher Scientific). Proteins were transferred to 0.45 µm nitrocellulose membranes and blocked in 5% w/v bovine serum albumin (BSA) in TBST for 1 hour (or 5% non-fat dry milk in TBST for CCPG1 probing). Membranes were then incubated overnight with primary antibody diluted in 2.5 % BSA in TBST (or 5% non-fat dry milk for CCPG1). Antibody dilutions are indicated in the Key Resource Table. Membranes were washed thrice with TBST for 10 min and incubated with the corresponding HRP-linked antibody. Membranes were then washed again three times prior incubation with ECL chemiluminescence reagents and visualised on an Amersham ImageQuant 800 Western blot imaging system (Cytiva) or alternatively using a film developer. Quantitative analyses were performed using ImageJ 2.9.0 (Gel analyzer plugin, NIH, Bethesda, USA).

### Immunoprecipitation

Input samples for immunoprecipitation (IP) were obtained from indicated PDAC cell lines seeded in 6 cm tissue culture dishes. For testing GFP-PRKAR1A mutants, 1x10^6^ PDAC WT cells stably expressing FAM134B-V5 or V5 were seeded in 6 cm tissue culture dishes and transfected the following day with the corresponding GFP-PRKAR1A constructs (3 µg) using lipofectamine. For FLAG-Immunoprecipitation, 1.5 x 10^6^ HEK293FT cells were plated on 6 cm tissue culture dishes and co-transfected the following day with FAM134-V5 tagged constructs (1.5 µg) and FLAG-tagged constructs or FLAG empty vector as a control (1.5 µg). 24 h after plating or transfection, IP was performed. If cross-linking was performed (as indicated in figure legends), 3,3′-Dithiodipropionic acid di(N-hydroxysuccinimide ester) (DSP)-crosslinking was used as described in ^20^. DSP was prepared fresh on the day of the IP. Media was removed from cells and washed twice with pre-warmed PBS (37°C), then cells were incubated with 1 mM DSP (diluted in PBS) for 30 min at 37°C at 5% CO_2_. Then, DSP solution was discarded and cells were washed with PBS once before incubating with STOP solution (20 mM Tris-HCl pH 7.5 in PBS) at room temperature for 15min. Cell extracts were obtained by washing twice with ice-cold PBS before lysis in 400 µl of IGEPAL cell lysis buffer (50 mM Tris-HCl pH 7.5, 150 mM NaCl, 0.5% IGEPAL, 2 mM activated sodium orthovanadate, 20 mM sodium fluoride, 10 mM sodium pyrophosphate, 1 mM PMSF and Complete Protease Inhibitor (Sigma-Aldrich, 11697498001)). Cells were incubated for 10 min with shaking and then lysates were transferred and thoroughly mixed in 1.5ml tubes and incubated on ice for another 10 min. Lysates were clarified by centrifugation at 13,000 g at 4°C for 10 min and 2% of the sample was kept as input reference.

For endogenous FAM134B immunoprecipitation, samples were then incubated at 4°C with rotation with 2 µg of α-FAM134B (Cell Signalling Technologies) or α-IgG rabbit as control for 3h. After this, samples were incubated for another 2 h with 20 µl protein A/G magnetic beads, previously washed thrice with IGEPAL lysis buffer. Alternatively, for V5 or FLAG immunoprecipitation, cleared lysates were incubated at 4°C with rotation with 10 µl of V5 magnetic beads (Chromotek) or 15 µl of FLAG agarose beads (Chromotek, HEK293FT lysates), previously washed thrice with IGEPAL lysis buffer. After incubation, beads were washed with IGEPAL lysis buffer three times using a magnetic rack – or by centrifugation at 2,500 g for 5 min for agarose beads – then washed once with 50 mM Tris-HCl, pH 7.5, 150 mM NaCl. After the final wash, supernatant was aspirated and 40 µl of 2× Laemmli buffer was added to the beads. Samples were boiled at 95°C for 5 min, and after separation of magnetic beads, supernatant was subjected to SDS-PAGE and immunoblotting.

### Recombinant protein and affinity pulldowns

Mouse FAM134B CT and human PRKAR1A NT+PS were cloned into pDEST 15 (for N-terminal tagging of GST) or pDEST 17 (For N-terminal tagging of His), respectively, and transformed into Rosetta II (DE3) cells. Cultures grown in L-broth were induced with 1 mM IPTG for 2.5 h. The culture was centrifuged at 1,500 g for 10 min at 4 °C, and the pellet resuspended in homogenization buffer (50 mM Tris-HCl, pH 7.5, 150 mM NaCl, 1 mM EDTA, 0.1 mM PMSF and Complete Protease Inhibitor (Sigma-Aldrich, 11697498001). Cells were sonicated five times in 15 s intervals at 4 °C and, for complete lysis, cells were subsequently incubated with 1% IGEPAL for 1 h at 4 °C with rotation. Cleared lysates were obtained by centrifugation at 12,000 *g* for 20 min at 4 °C and protein expression was confirmed by SDS-PAGE followed by Coomassie staining or immunoblotting. For affinity precipitation (AP), 20 µl/reaction Glutathione Sepharose 4B beads were washed thrice in IGEPAL buffer (50 mM Tris-HCl pH 7.5, 150 mM NaCl, 0.5% IGEPAL and Complete Protease Inhibitor) and incubated with GST-fusion protein containing lysate (i.e., GST or GST-FAM134B CT) for 2h with rotation at 4°C. GST-fusion protein-bound beads were washed thrice in PBS with Complete Protease Inhibitors and then bacterial His-PRKAR1A NT+PS lysate was incubated with beads in AP buffer (10 mM Tris-HCl pH 7.5, 100 mg/ml PEG4000, with Complete Protease Inhibitors) (1 ml total volume) for 3 h with rotation at 4°C. Then, beads were washed thrice with AP buffer and then resuspended in 40 µl 2x Laemmli buffer. Samples were boiled at 95°C for 5 min, and supernatant was subjected to immunoblotting.

### Rhotekin-RBD active RhoA pull-down

3 x 10^6^ PDAC-Cas9 cells, either *ΔFam134b+*c transduced with V5 empty vector virus or reconstituted with FAM134B-V5, were seeded in 10 cm tissue culture dishes and incubated at 37°C at 5% CO_2_. The following day, cells were treated with DMSO or FSK. After treatment, cells were washed twice with ice-cold PBS and then lysed in 700 µl of RhoA lysis buffer (25 mM HEPES, pH 7.5, 150 mM NaCl, 1% IGEPAL, 10 mM MgCl2, 1 mM EDTA, 2% Glycerol, 0.1 mM PMSF and Complete Protease Inhibitor (Sigma-Aldrich, 11697498001)). Cells were incubated for 10 min and scraped into 1.5ml tubes and incubated on ice for another 10 min. Lysates were clarified by centrifugation at 13,000 *g* at 4°C for 10 min and 2% of the sample was kept as input. Then, 40 µl of GST-Rhotekin-RBD beads (Cell Biolabs) were added to the lysates for 1.5 h and incubated at 4°C with rotation. Beads were washed with RhoA lysis buffer three times by centrifugation at 14,000 *g* for 10 s. After the last wash, supernatant was aspirated and 40 µl of 2× Laemmli buffer was added to the beads. Samples were boiled at 95°C for 5 min, and supernatant was subjected to SDS-PAGE and immunoblotting.

### Scratch wound healing assays

Indicated PDAC cell lines were plated on 48-well plates for 24 h wound closure assays (200,000 cells) or on 12mm coverslips (450,000 cells) for F-actin staining and incubated at 37°C at 5% CO_2_. The following day, using a pipette tip, a linear scratch in the middle of the well – or coverslip – was created. Media was then removed and cells were washed once with low-serum media (1% FBS). Cells were then treated with DMSO or FSK in low-serum media. For 24 h wound closure assays, 1h after treatment was added, cells were imaged with time-lapse automated acquisition with 30min intervals using a Nikon Eclipse Ti-E inverted microscope (Nikon Instruments UK Ltd., Kingston Upon Thames, UK) with a 4X 0.13 NA objective. Phase contrast images were acquired with a Photometrics Prime BSI sCMOS camera (Teledyne Photometrics, 3440 E. Britannia Drive, Suite 100, Tucson, AZ 85706). Cells were maintained at 37°C using a custom-made environmental enclosure (Institute of Genetics and Cancer, University of Edinburgh, UK). A 5% CO_2_ atmosphere was maintained by a Okolabs chamber (Okolabs Pozzuoli NA, Italy). Wound closure was calculated using the time 0 and time 22h from our time lapse with a readily available pipeline on GitHub to calculate the wound area 21, 22. For F-actin staining, coverslips were fixed with 4% paraformaldehyde in PBS 7h after treatment/scratch and cells were then permeabilized with 0.25% Triton X-100 (15 min) and blocked for 1 h at room temperature in 5% BSA/PBS/0.02% sodium azide, before incubating with Phalloidin-FITC (Thermo Fisher Scientific) for 1 h at room temperature. Quantification of single cells in wound was done with QuPath by manually counting the number of F-actin-positive single cells disseminating from the leading edge of the wound.

### 3D spheroid collagen invasion assay

PDAC, PDAC *ΔPRKAR1A*, PDAC *ΔFAM134B/C* were seeded in 96-well ultra-low attachment plates (Corning) (4,000 cells/well). Plate was centrifuged at 125 g for 10 min at room temperature and then cultured at 37°C at 5% CO_2_ for 3 days to allow spheroid formation. On day 3, 3.38 mg/ml Collagen type I (Corning) was added on top to the pre-chilled plate containing the spheroids, the plate was then centrifuged 200 g for 3min at 4°C as per manufacturer’s instructions (IncuCyte S3 3D Spheroid Invasion Assay, Sartorius). After that, plate was incubated at 37 °C in 5 % CO_2_ to allow polymerization for 30 min before topping up with full medium. Then, spheroids were placed in IncuCyte S3 (Sartorius) with a phase contrast and brightfield single spheroid scan type using a 4x lens and S3/SX1 G/R optical module to follow invasion for 48 h (4 h intervals). The day after collagen addition, treatment (DMSO or 40 µM FSK – see below for adrenaline-treated spheroids) was added in full medium to 1 x final concentration. Collective invasion index was calculated in Incucyte software with Spheroid invasion analysis with a measurement of largest Invading Cells Brightfield Object Area (µm/A²). After 24 h, spheroids were fixed in by adding 16% paraformaldehyde (Thermo Fisher Scientific) to a final concentration of 4%. Then, spheroids were permeabilized with 0.25% Triton X-100 in PBS for 15min and stained with Phalloidin-FITC for 1 h. After PBS washes, spheroids were mounted onto slides with DAKO mounting media using a micro spoon-shaped spatula. Fluorescence images were acquired using a 10x lens in a BX51 widefield fluorescence Microscope (Olympus) with a DP71 digital microscope camera (Olympus) using Cell F software (Olympus). For analysis, the number of F-actin-positive single cells disseminating from the spheroid and invading into the collagen was counted using ImageJ. For adrenaline-treated spheroids, cells were kept with low serum media from the time of plating (1% FBS). Then, 3 days after plating, together with 3.38 mg/ml Collagen type I, treatment was added (adrenaline 50 µM (freshly made) or DMSO). Treatment was also reapplied the following day. Spheroids were fixed 48 h later as described above.

### Image quantifications

ImageJ was used for: line traces using the plot profile tool; manual counting of protrusion number per cell on peripherial cells; total P-MLC2 cytosolic fluorescence intensity using intensity measurement over a threshold and removing nuclear staining using DAPI as a mask; spheroid single cell invasion using analyse measure particles tool for phalloidin-FITC positive cells outside the spheroid core. Imaris was used for foci number quantifications using the spots tool and batch processing and colocalization using the spot function “Colocalize spots” (Matlab). QuPath was used for manual quantification of the number of F-actin-positive single cells disseminating from the leading edge of the wound over the total number of cells in the periphery of the leading edge.

### Statistics

Statistical tests were performed as described in figure legends using GraphPad Prism 10 (GraphPad Software). Data distribution was assumed to be normal, but this was not formally tested. Results are presented as mean ± s.e.m. or mean ± s.d., as detailed in figure legends. *n* and independent or technical replicate status is indicated in the figure legends. Where values within an independent replicate are normalised to a shown control condition, a 1-sample t-test was employed for comparisons with that control. Multiple testing correction for *P* was performed as described in individual figure legends.

## References

1. Wu, H., Carvalho, P. & Voeltz, G.K. Here, there, and everywhere: The importance of ER membrane contact sites. Science 361 (2018).

2. Gubas, A. & Dikic, I. ER remodeling via ER-phagy. Mol Cell 82, 1492–1500 (2022).

3. Reggiori, F. & Molinari, M. ER-phagy: mechanisms, regulation, and diseases connected to the lysosomal clearance of the endoplasmic reticulum. Physiol Rev 102, 1393–1448 (2022).

4. Mochida, K. & Nakatogawa, H. ER-phagy: selective autophagy of the endoplasmic reticulum. EMBO Rep 23, e55192 (2022).

5. Reggio, A. et al. Role of FAM134 paralogues in endoplasmic reticulum remodeling, ER-phagy, and Collagen quality control. EMBO Rep 22, e52289 (2021).

6. Khaminets, A. et al. Regulation of endoplasmic reticulum turnover by selective autophagy. Nature 522, 354–358 (2015).

7. Fumagalli, F. et al. Translocon component Sec62 acts in endoplasmic reticulum turnover during stress recovery. Nat Cell Biol 18, 1173–1184 (2016).

8. Grumati, P. et al. Full length RTN3 regulates turnover of tubular endoplasmic reticulum via selective autophagy. Elife 6 (2017).

9. Smith, M.D. et al. CCPG1 Is a Non-canonical Autophagy Cargo Receptor Essential for ER- Phagy and Pancreatic ER Proteostasis. Dev Cell 44, 217–232 e211 (2018).

10. An, H. et al. TEX264 Is an Endoplasmic Reticulum-Resident ATG8-Interacting Protein Critical for ER Remodeling during Nutrient Stress. Mol Cell 74, 891–908 e810 (2019).

11. Chino, H., Hatta, T., Natsume, T. & Mizushima, N. Intrinsically Disordered Protein TEX264 Mediates ER-phagy. Mol Cell 74, 909–921 e906 (2019).

12. Stephani, M. et al. A cross-kingdom conserved ER-phagy receptor maintains endoplasmic reticulum homeostasis during stress. Elife 9 (2020).

13. Nthiga, T.M. et al. CALCOCO1 acts with VAMP-associated proteins to mediate ER-phagy. EMBO J 39, e103649 (2020).

14. Forrester, A. et al. A selective ER-phagy exerts procollagen quality control via a Calnexin- FAM134B complex. EMBO J 38 (2019).

15. Fregno, I. et al. ER-to-lysosome-associated degradation of proteasome-resistant ATZ polymers occurs via receptor-mediated vesicular transport. Embo j 37 (2018).

16. Salomo Coll, C. et al. ER-phagy and proteostasis defects prime pancreatic epithelial state changes in KRAS-mediated oncogenesis. Dev Cell (2025).

17. Ishii, S. et al. CCPG1 recognizes endoplasmic reticulum luminal proteins for selective ER- phagy. Molecular Biology of the Cell 34 (2023).

18. Herhaus, L. et al. IRGQ-mediated autophagy in MHC class I quality control promotes tumor immune evasion. Cell (2024).

19. Parashar, S. et al. Endoplasmic reticulum tubules limit the size of misfolded protein condensates. Elife 10 (2021).

20. Cunningham, C.N. et al. Cells Deploy a Two-Pronged Strategy to Rectify Misfolded Proinsulin Aggregates. Mol Cell 75, 442–456.e444 (2019).

21. Ji, C.H. et al. The N-Degron Pathway Mediates ER-phagy. Mol Cell 75, 1058–1072.e1059 (2019).

22. Hayashi, Y. et al. TOLLIP acts as a cargo adaptor to promote lysosomal degradation of aberrant ER membrane proteins. Embo j 42, e114272 (2023).

23. Schultz, M.L. et al. Coordinate regulation of mutant NPC1 degradation by selective ER autophagy and MARCH6-dependent ERAD. Nat Commun 9, 3671 (2018).

24. Loi, M., Raimondi, A., Morone, D. & Molinari, M. ESCRT-III-driven piecemeal micro-ER-phagy remodels the ER during recovery from ER stress. Nat Commun 10, 5058 (2019).

25. Omari, S. et al. Noncanonical autophagy at ER exit sites regulates procollagen turnover. Proc Natl Acad Sci U S A 115, E10099–e10108 (2018).

26. Schuck, S., Gallagher, C.M. & Walter, P. ER-phagy mediates selective degradation of endoplasmic reticulum independently of the core autophagy machinery. Journal of Cell Science 127, 4078–4088 (2014).

27. Kim, D.Y. et al. A selective ER-phagy exerts neuroprotective effects via modulation of α- synuclein clearance in parkinsonian models. Proc Natl Acad Sci U S A 120, e2221929120 (2023).

28. Mou, W., Tang, Y., Huang, Y., Wu, Z. & Cui, Y. Upregulation of neuronal ER-phagy improves organismal fitness and alleviates APP toxicity. Cell Rep 43, 114255 (2024).

29. Buonomo, V. et al. Two FAM134B isoforms differentially regulate ER dynamics during myogenesis. Embo j 44, 1039–1073 (2025).

30. Iavarone, F. et al. Fam134c and Fam134b shape axonal endoplasmic reticulum architecture in vivo. EMBO Rep 25, 3651–3677 (2024).

31. González, A. et al. Ubiquitination regulates ER-phagy and remodelling of endoplasmic reticulum. Nature 618, 394–401 (2023).

32. Berkane, R. et al. The function of ER-phagy receptors is regulated through phosphorylation- dependent ubiquitination pathways. Nat Commun 14, 8364 (2023).

33. De Leonibus, C. et al. Sestrin2 drives ER-phagy in response to protein misfolding. Dev Cell 59, 2035–2052 e2010 (2024).

34. Di Lorenzo, G. et al. Phosphorylation of FAM134C by CK2 controls starvation-induced ER- phagy. Sci Adv 8, eabo1215 (2022).

35. Cinque, L. et al. MiT/TFE factors control ER-phagy via transcriptional regulation of FAM134B. EMBO J 39, e105696 (2020).

36. Liang, J.R. et al. A Genome-wide ER-phagy Screen Highlights Key Roles of Mitochondrial Metabolism and ER-Resident UFMylation. Cell 180, 1160–1177 e1120 (2020).

37. Jiang, X. et al. FAM134B oligomerization drives endoplasmic reticulum membrane scission for ER-phagy. EMBO J 39, e102608 (2020).

38. Foronda, H. et al. Heteromeric clusters of ubiquitinated ER-shaping proteins drive ER-phagy. Nature 618, 402–410 (2023).

39. Rudinskiy, M., Galli, C., Raimondi, A. & Molinari, M. The intrinsically disordered regions of organellophagy receptors are interchangeable and control organelle fragmentation, ER- phagy and mitophagy flux. Nat Cell Biol (2025).

40. Poveda-Cuevas, S.A. et al. Intrinsically disordered region amplifies membrane remodeling to augment selective ER-phagy. Proc Natl Acad Sci U S A 121, e2408071121 (2024).

41. Holehouse, A.S. & Kragelund, B.B. The molecular basis for cellular function of intrinsically disordered protein regions. Nat Rev Mol Cell Biol 25, 187–211 (2024).

42. Kumar, D. et al. RTN4B interacting protein FAM134C promotes ER membrane curvature and has a functional role in autophagy. Mol Biol Cell 32, 1158–1170 (2021).

43. Wang, B. et al. Liquid-liquid phase separation in human health and diseases. Signal Transduct Target Ther 6, 290 (2021).

44. Su, Q., Mehta, S. & Zhang, J. Liquid-liquid phase separation: Orchestrating cell signaling through time and space. Mol Cell 81, 4137–4146 (2021).

45. Yang, Z. et al. Autophagy adaptors mediate Parkin-dependent mitophagy by forming sheet- like liquid condensates. EMBO J 43, 5613–5634 (2024).

46. Licheva, M. et al. Phase separation of initiation hubs on cargo is a trigger switch for selective autophagy. Nat Cell Biol 27, 283–297 (2025).

47. Shao, Y. et al. Biomolecular condensates of ATG18 reshape ER for autophagy in plants. Dev Cell (2025).

48. Fujioka, Y. et al. Phase separation promotes Atg8 lipidation and vesicle condensation for autophagy progression. Nat Struct Mol Biol (2025).

49. Zheng, Q. et al. Calcium transients on the ER surface trigger liquid-liquid phase separation of FIP200 to specify autophagosome initiation sites. Cell 185, 4082–4098.e4022 (2022).

50. Hoffmann, M. E. et al. TBC1D2B undergoes phase separation and mediates autophagy initiation. J Cell Biochem 125, e30481 (2024).

51. Zaffagnini, G. et al. p62 filaments capture and present ubiquitinated cargos for autophagy. EMBO J 37 (2018).

52. Gallagher, E.R. & Holzbaur, E.L.F. The selective autophagy adaptor p62/SQSTM1 forms phase condensates regulated by HSP27 that facilitate the clearance of damaged lysosomes via lysophagy. Cell Rep 42, 112037 (2023).

53. Sun, D., Wu, R., Zheng, J., Li, P. & Yu, L. Polyubiquitin chain-induced p62 phase separation drives autophagic cargo segregation. Cell Res 28, 405–415 (2018).

54. Zhang, J.Z. et al. Phase Separation of a PKA Regulatory Subunit Controls cAMP Compartmentation and Oncogenic Signaling. Cell 182, 1531–1544 e1515 (2020).

55. Hardy, J.C. et al. Molecular determinants and signaling effects of PKA RIalpha phase separation. Mol Cell 84, 1570–1584 e1577 (2024).

56. Svec, K.V. & Howe, A.K. Protein Kinase A in cellular migration-Niche signaling of a ubiquitous kinase. Front Mol Biosci 9, 953093 (2022).

57. Sassone-Corsi, P. The Cyclic AMP Pathway. Cold Spring Harbor Perspectives in Biology 4, a011148–a011148 (2012).

58. Branon, T.C. et al. Efficient proximity labeling in living cells and organisms with TurboID. Nat Biotechnol 36, 880–887 (2018).

59. Jimenez-Moreno, N., Salomo-Coll, C., Murphy, L.C. & Wilkinson, S. Signal-Retaining Autophagy Indicator as a Quantitative Imaging Method for ER-Phagy. Cells 12 (2023).

60. Katayama, H. et al. Visualizing and Modulating Mitophagy for Therapeutic Studies of Neurodegeneration. Cell 181, 1176–1187 e1116 (2020).

61. Chijiwa, T. et al. Inhibition of forskolin-induced neurite outgrowth and protein phosphorylation by a newly synthesized selective inhibitor of cyclic AMP-dependent protein kinase, N-[2-(p-bromocinnamylamino)ethyl]-5-isoquinolinesulfonamide (H-89), of PC12D pheochromocytoma cells. J Biol Chem 265, 5267–5272 (1990).

62. Akaki, K., Mino, T. & Takeuchi, O. DSP-crosslinking and Immunoprecipitation to Isolate Weak Protein Complex. Bio Protoc 12 (2022).

63. Taylor, S.S. et al. Dynamics of signaling by PKA. Biochim Biophys Acta 1754, 25–37 (2005).

64. Deng, Z. et al. Selective autophagy of AKAP11 activates cAMP/PKA to fuel mitochondrial metabolism and tumor cell growth. Proc Natl Acad Sci U S A 118 (2021).

65. Segura-Roman, A. et al. Autophagosomes anchor an AKAP11-dependent regulatory checkpoint that shapes neuronal PKA signaling. EMBO J 44, 3150–3179 (2025).

66. Bhaskara, R.M. et al. Curvature induction and membrane remodeling by FAM134B reticulon homology domain assist selective ER-phagy. Nat Commun 10, 2370 (2019).

67. Zhao, J., Li, Z. & Li, J. The crystal structure of the FAM134B–GABARAP complex provides mechanistic insights into the selective binding of FAM134 to the GABARAP subfamily. FEBS Open Bio 12, 320–331 (2022).

68. Ganser, L.R. & Myong, S. Methods to Study Phase-Separated Condensates and the Underlying Molecular Interactions. Trends Biochem Sci 45, 1004–1005 (2020).

69. Ribbeck, K. & Gorlich, D. The permeability barrier of nuclear pore complexes appears to operate via hydrophobic exclusion. EMBO J 21, 2664–2671 (2002).

70. Croft, D. et al. Reactome: a database of reactions, pathways and biological processes. Nucleic Acids Res 39, D691–697 (2011).

71. Sahai, E. & Marshall, C.J. RHO-GTPases and cancer. Nat Rev Cancer 2, 133–142 (2002).

72. Vicente-Manzanares, M., Ma, X., Adelstein, R.S. & Horwitz, A.R. Non-muscle myosin II takes centre stage in cell adhesion and migration. Nat Rev Mol Cell Biol 10, 778–790 (2009).

73. Self, A.J. & Hall, A. Purification of recombinant Rho/Rac/G25K from Escherichia coli. Methods Enzymol 256, 3–10 (1995).

74. Ren, X.D., Kiosses, W.B. & Schwartz, M.A. Regulation of the small GTP-binding protein Rho by cell adhesion and the cytoskeleton. EMBO J 18, 578–585 (1999).

75. Uehata, M. et al. Calcium sensitization of smooth muscle mediated by a Rho-associated protein kinase in hypertension. Nature 389, 990–994 (1997).

76. Zimmerman, N.P. et al. Cyclic AMP regulates the migration and invasion potential of human pancreatic cancer cells. Mol Carcinog 54, 203–215 (2015).

77. Pandya, P., Orgaz, J.L. & Sanz-Moreno, V. Modes of invasion during tumour dissemination. Mol Oncol 11, 5–27 (2017).

78. Gong, B., Johnston, J.D., Thiemicke, A., de Marco, A. & Meyer, T. Endoplasmic reticulum- plasma membrane contact gradients direct cell migration. Nature 631, 415–423 (2024).

79. Rawal, S. et al. Edge curvature drives endoplasmic reticulum reorganization and dictates epithelial migration mode. Nat Cell Biol 27, 1660–1675 (2025).

80. Tikhomirova, M.S., Kadosh, A., Saukko-Paavola, A.J., Shemesh, T. & Klemm, R.W. A role for endoplasmic reticulum dynamics in the cellular distribution of microtubules. Proc Natl Acad Sci U S A 119, e2104309119 (2022).

81. Hoyer, M.J. et al. Combinatorial selective ER-phagy remodels the ER during neurogenesis. Nat Cell Biol 26, 378–392 (2024).

82. Zhou, X. et al. Integrated proteomics reveals autophagy landscape and an autophagy receptor controlling PKA-RI complex homeostasis in neurons. Nat Commun 15, 3113 (2024).

83. Ordureau, A. et al. Temporal proteomics during neurogenesis reveals large-scale proteome and organelle remodeling via selective autophagy. Mol Cell 81, 5082–5098 e5011 (2021).

84. Li, B. et al. Cholesterol sensing by the SCAP-FAM134B complex regulates ER-phagy and STING innate immunity. Nat Cell Biol 27, 1739–1756 (2025).

85. Kajiho, H. & Sakisaka, T. Degradation of STIM1 through FAM134B-mediated ER-phagy is potentially involved in cell proliferation. J Biol Chem 300, 107674 (2024).

86. Sheehan, B.K., Orefice, N.S., Peng, Y., Shapiro, S. L. & Puglielli, L. ATG9A regulates proteostasis through reticulophagy receptors FAM134B and SEC62 and folding chaperones CALR and HSPB1. iScience 24, 102315 (2021).

87. Peng, S.Z. et al. Phase separation of Nur77 mediates celastrol-induced mitophagy by promoting the liquidity of p62/SQSTM1 condensates. Nat Commun 12, 5989 (2021).

88. Pool, E.H. et al. Aberrant phase separation of two PKA RIbeta neurological disorder mutants leads to mechanistically distinct signaling deficits. Cell Rep 44, 115797 (2025).

89. Lee, H.N. et al. Phase separation of a PKA type I regulatory subunit regulates beta-cell function through cAMP compartmentalization. PLoS Biol 23, e3003262 (2025).

90. Mavrakis, M., Lippincott-Schwartz, J., Stratakis, C.A. & Bossis, I. mTOR kinase and the regulatory subunit of protein kinase A (PRKAR1A) spatially and functionally interact during autophagosome maturation. Autophagy 3, 151–153 (2007).

91. Overhoff, M. et al. Autophagy regulates neuronal excitability by controlling cAMP/protein kinase A signaling at the synapse. EMBO J 41, e110963 (2022).

92. Belaid, A. et al. Autophagy plays a critical role in the degradation of active RHOA, the control of cell cytokinesis, and genomic stability. Cancer Res 73, 4311–4322 (2013).

## References for Extended Methods

1. Li, A. et al. Fascin is regulated by slug, promotes progression of pancreatic cancer in mice, and is associated with patient outcomes. Gastroenterology 146, 1386–1396 e1381-1317 (2014).

2. Stringer, B.W. et al. A reference collection of patient-derived cell line and xenograft models of proneural, classical and mesenchymal glioblastoma. Sci Rep 9, 4902 (2019).

3. Mali, P. et al. RNA-guided human genome engineering via Cas9. Science 339, 823–826 (2013).

4. Newman, A.C. et al. TBK1 kinase addiction in lung cancer cells is mediated via autophagy of Tax1bp1/Ndp52 and non-canonical NF-kappaB signalling. PLoS One 7, e50672 (2012).

5. Smith, M.D. et al. CCPG1 Is a Non-canonical Autophagy Cargo Receptor Essential for ER-Phagy and Pancreatic ER Proteostasis. Dev Cell 44, 217–232 e211 (2018).

6. Branon, T.C. et al. Efficient proximity labeling in living cells and organisms with TurboID. Nat Biotechnol 36, 880–887 (2018).

7. Jimenez-Moreno, N., Salomo-Coll, C., Murphy, L.C. & Wilkinson, S. Signal-Retaining Autophagy Indicator as a Quantitative Imaging Method for ER-Phagy. Cells 12 (2023).

8. Yang, X. et al. A public genome-scale lentiviral expression library of human ORFs. Nat Methods 8, 659–661 (2011).

9. Ran, F.A. et al. Genome engineering using the CRISPR-Cas9 system. Nat Protoc 8, 2281–2308 (2013).

10. Abramson, J. et al. Accurate structure prediction of biomolecular interactions with AlphaFold 3. Nature 630, 493–500 (2024).

11. Shannon, P. et al. Cytoscape: a software environment for integrated models of biomolecular interaction networks. Genome Res 13, 2498–2504 (2003).

12. Delgado, J., Radusky, L.G., Cianferoni, D. & Serrano, L. FoldX 5.0: working with RNA, small molecules and a new graphical interface. Bioinformatics 35, 4168–4169 (2019).

13. Schneider, C.A., Rasband, W.S. & Eliceiri, K.W. NIH Image to ImageJ: 25 years of image analysis. Nat Methods 9, 671–675 (2012).

14. Kong, A.T., Leprevost, F.V., Avtonomov, D.M., Mellacheruvu, D. & Nesvizhskii, A.I. MSFragger: ultrafast and comprehensive peptide identification in mass spectrometry-based proteomics. Nat Methods 14, 513–520 (2017).

15. Bankhead, P. et al. QuPath: Open source software for digital pathology image analysis. Sci Rep 7, 16878 (2017).

16. Szklarczyk, D. et al. The STRING database in 2023: protein-protein association networks and functional enrichment analyses for any sequenced genome of interest. Nucleic Acids Res 51, D638–D646 (2023).

17. Aftab, W. (GitHub, GitHub; 2022).

18. Murphy, L.C., Pearson, M., Jimenez Moreno, N. & Wilkinson, S. (GitHub, GitHub; 2023).

19. Reynolds, E.S. The use of lead citrate at high pH as an electron-opaque stain in electron microscopy. J Cell Biol 17, 208–212 (1963).

20. Akaki, K., Mino, T. & Takeuchi, O. DSP-crosslinking and Immunoprecipitation to Isolate Weak Protein Complex. Bio Protoc 12 (2022).

21. Suarez-Arnedo, A. et al. An image J plugin for the high throughput image analysis of in vitro scratch wound healing assays. PLoS One 15, e0232565 (2020).

22. Arnedo, A. (Github, Github; 2020).

