## Supplementary material for "ER-liquid condensate contacts sequester FAM134B/C and RhoA to govern cell morphology": Raw immunoblots

Raw (uncropped) immunoblot data

### Figure 1C

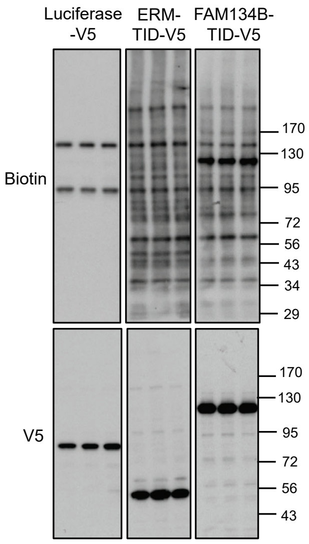


### Figure S1B (left panel)

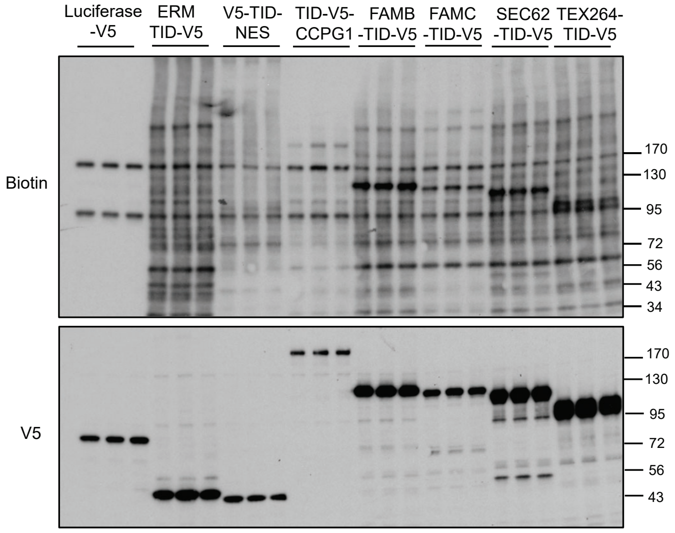


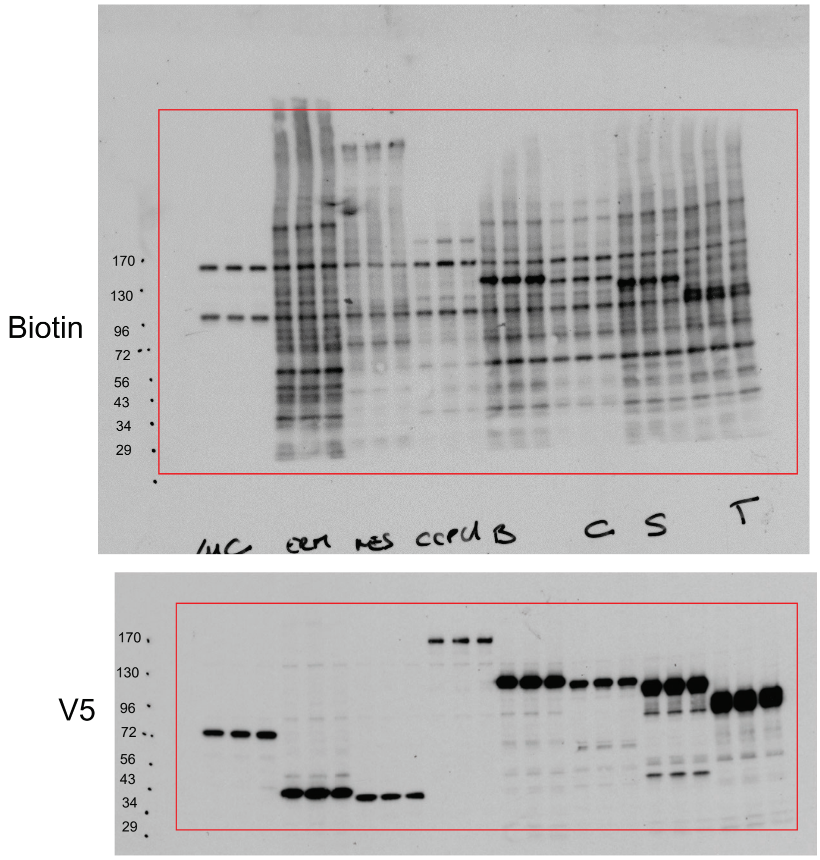

### Figure S1A

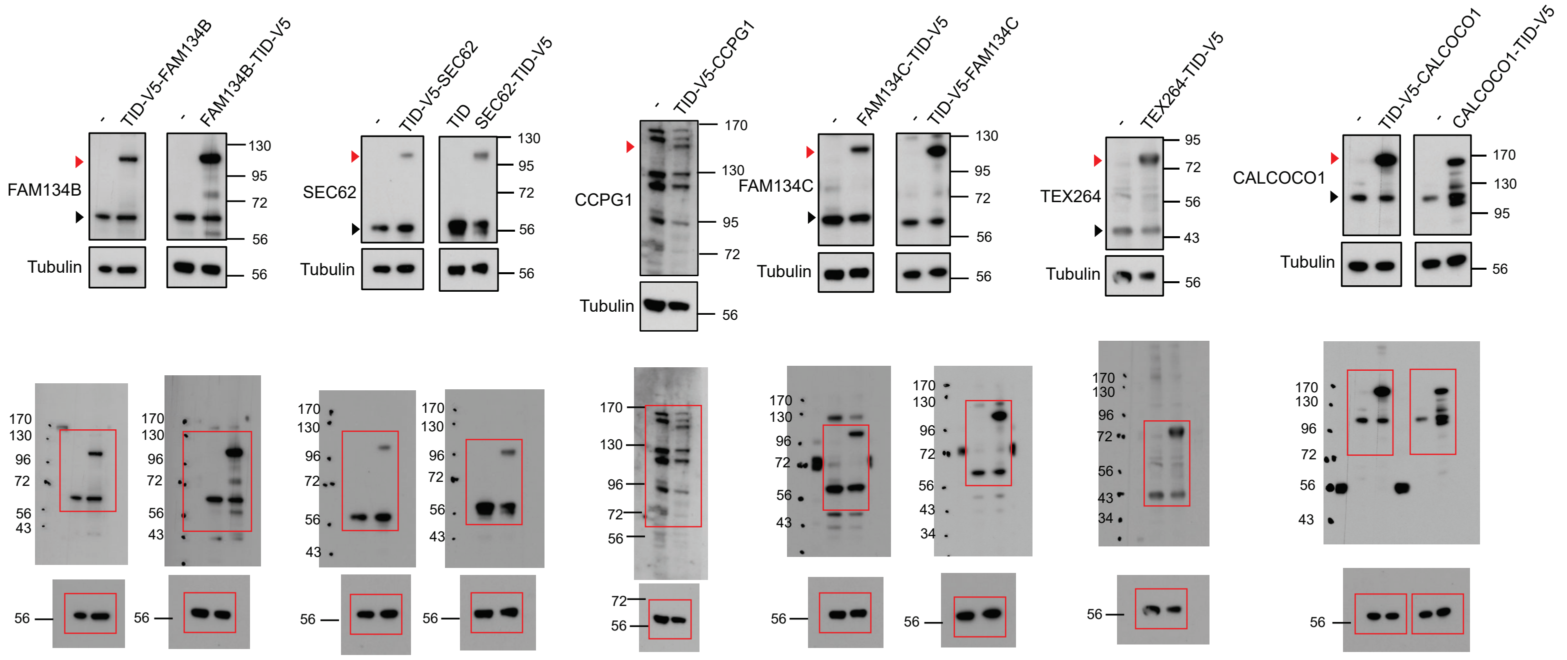


### Figure S1B (right panel)

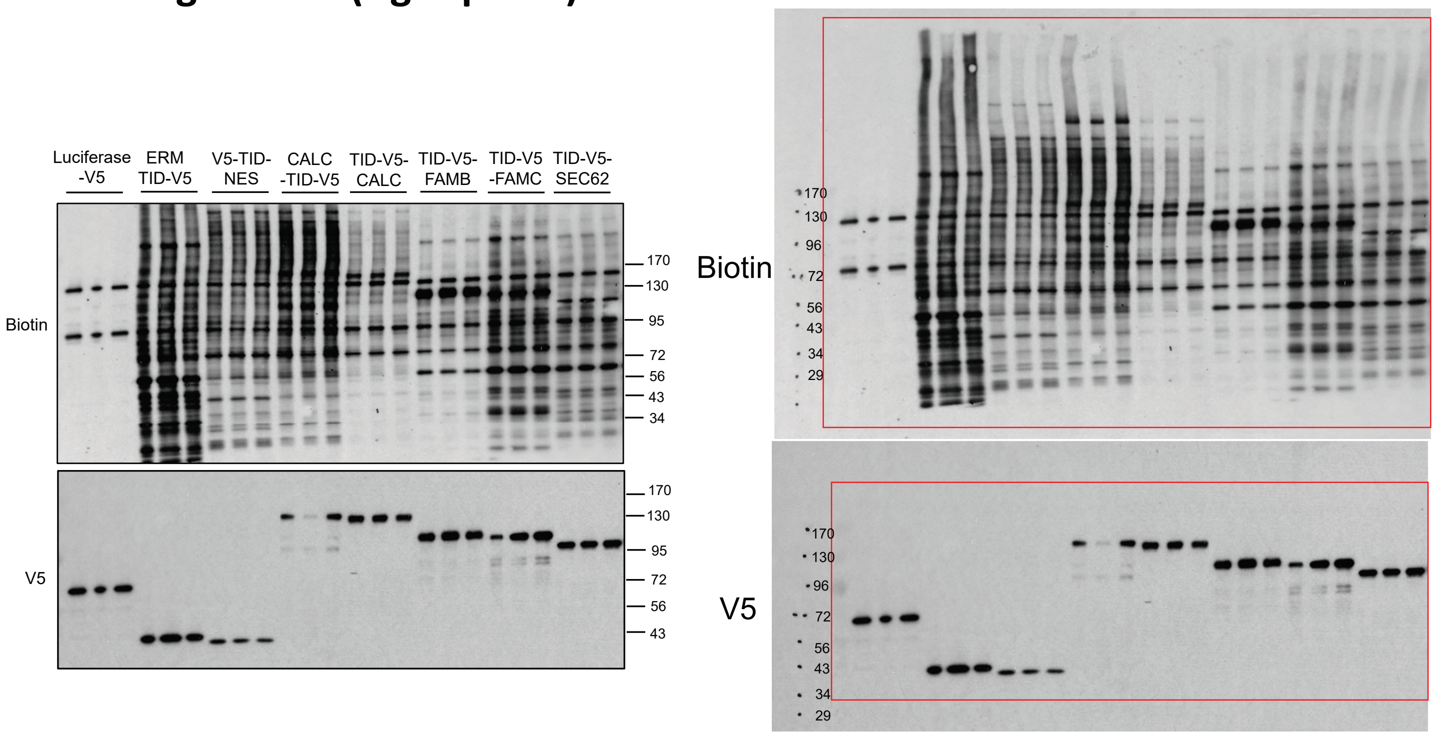

#### Figure 2E

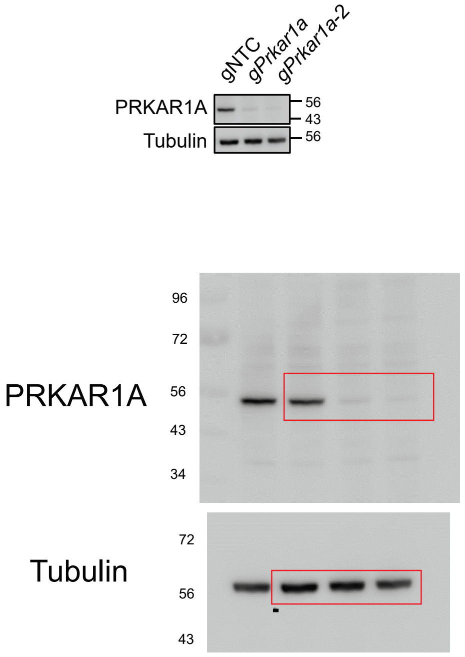


#### Figure 2G

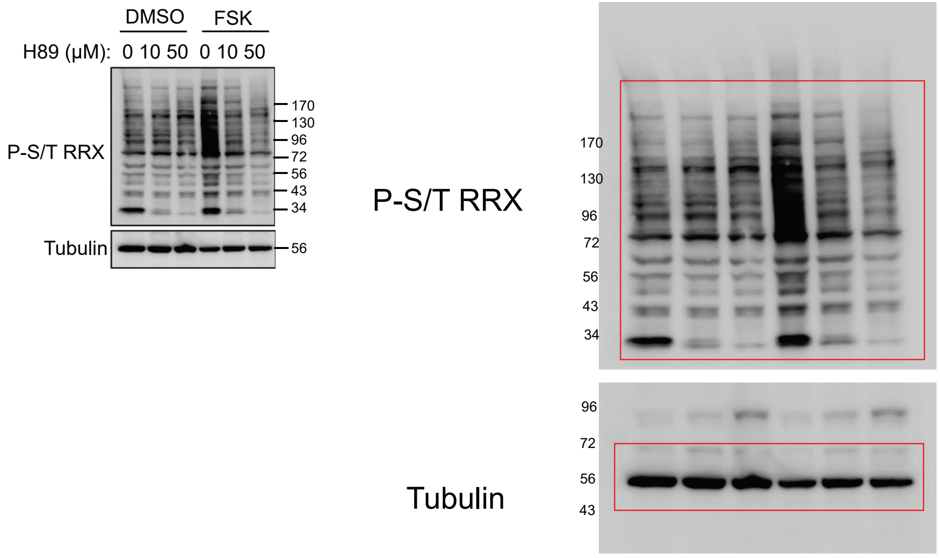

### Figure S2B

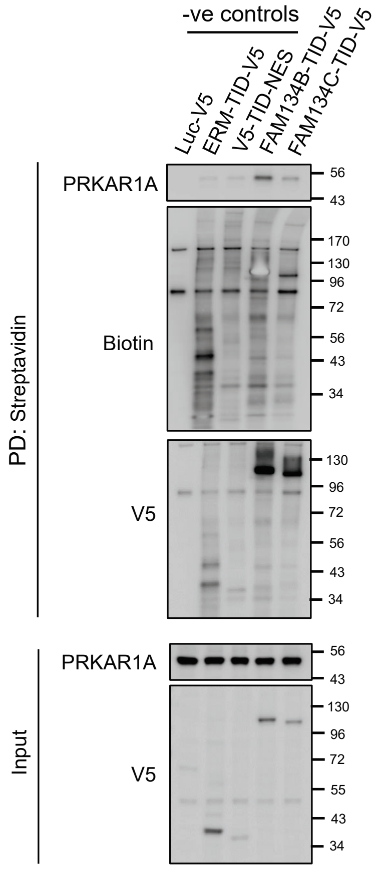


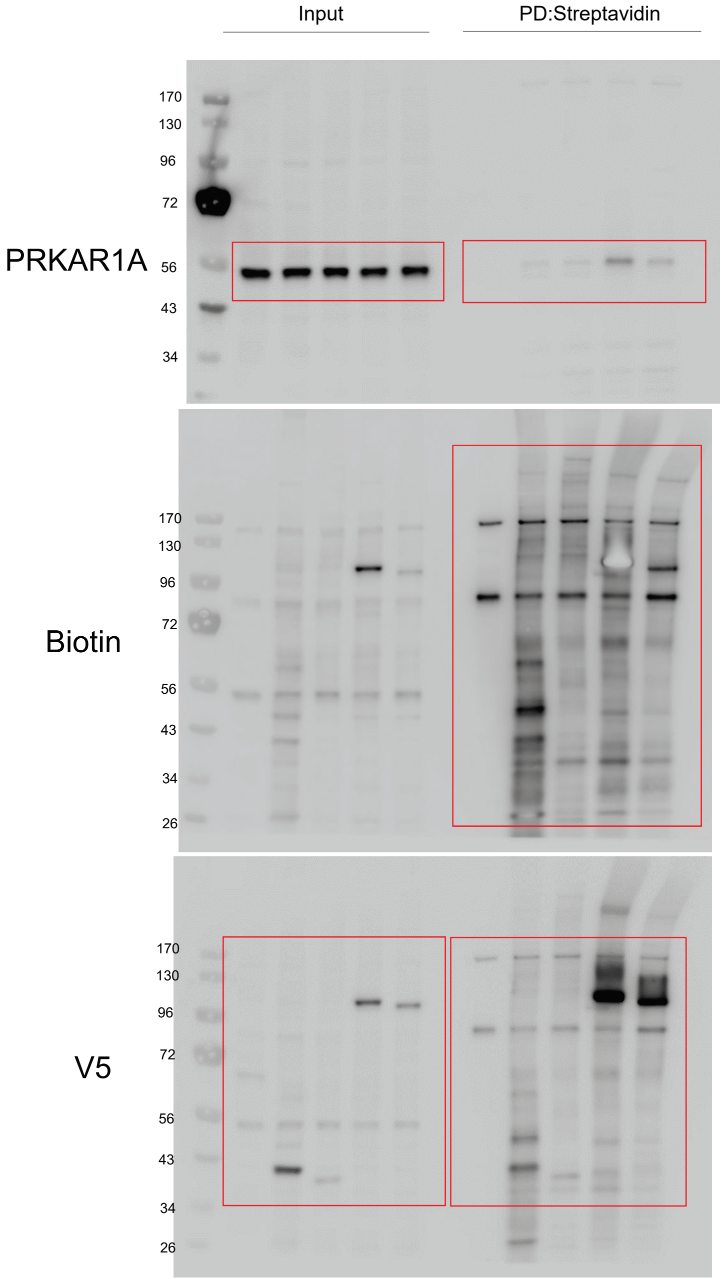


### Figure S2C

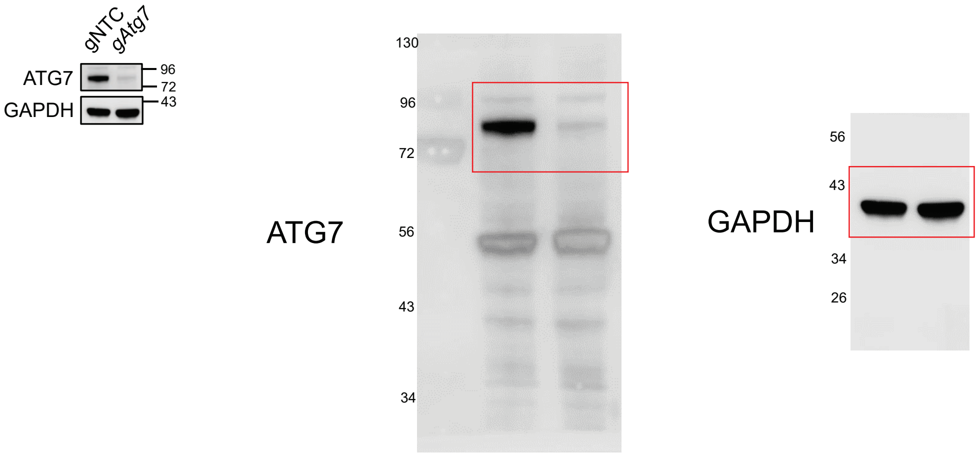

### Figure 3A

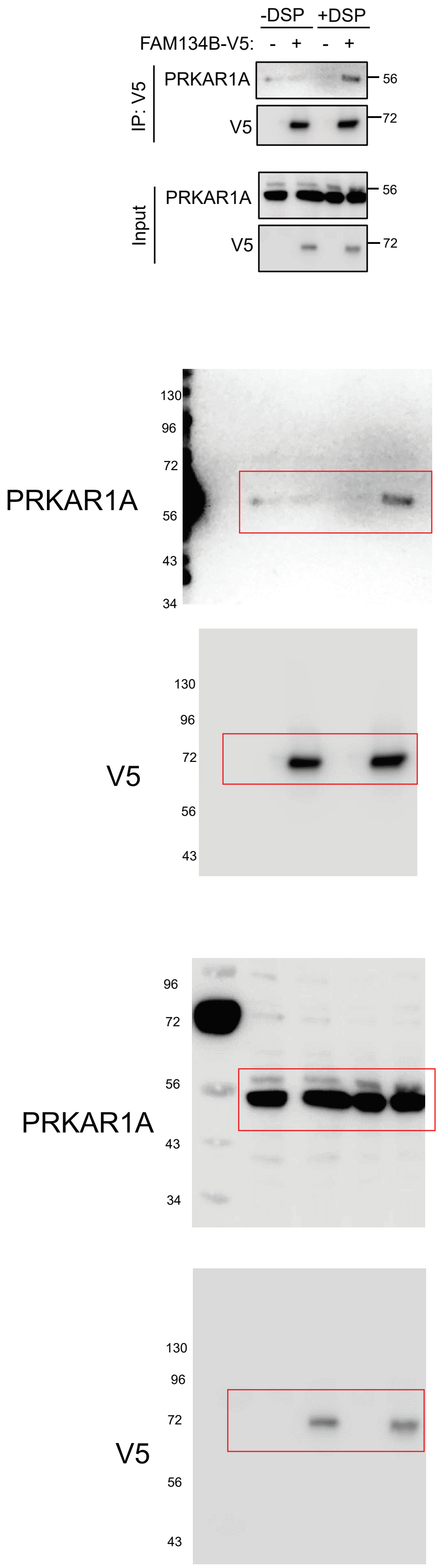


### Figure 3B

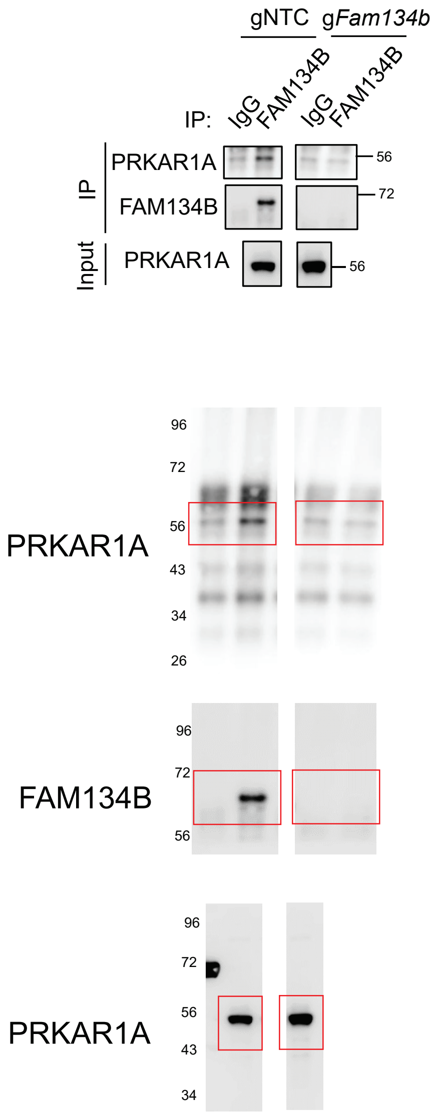

### Figure 3D

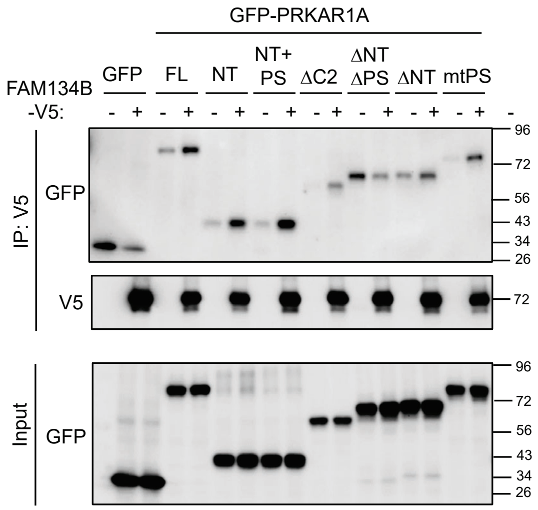

GFP-PRKAR1A
FAM134B -V5:
GFP
FL
NT
NT+ PS
ΔC2
ΔNT ΔPS
ΔNT
mtPS
- + - + - + - + - + - + - + - + -
IP: V5
GFP
V5
Input
GFP
96
72
56
43
34
26


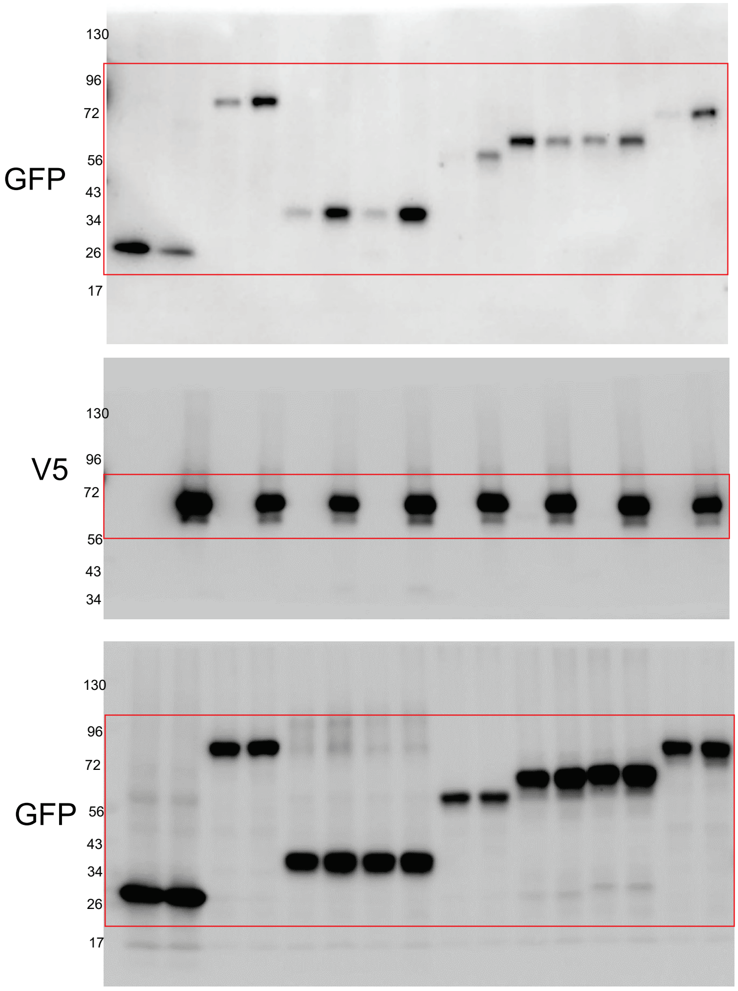

GFP
130
96
72
56
43
34
26
17
V5
130
96
72
56
43
34
GFP
130
96
72
56
43
34
26
17

### Figure 3F

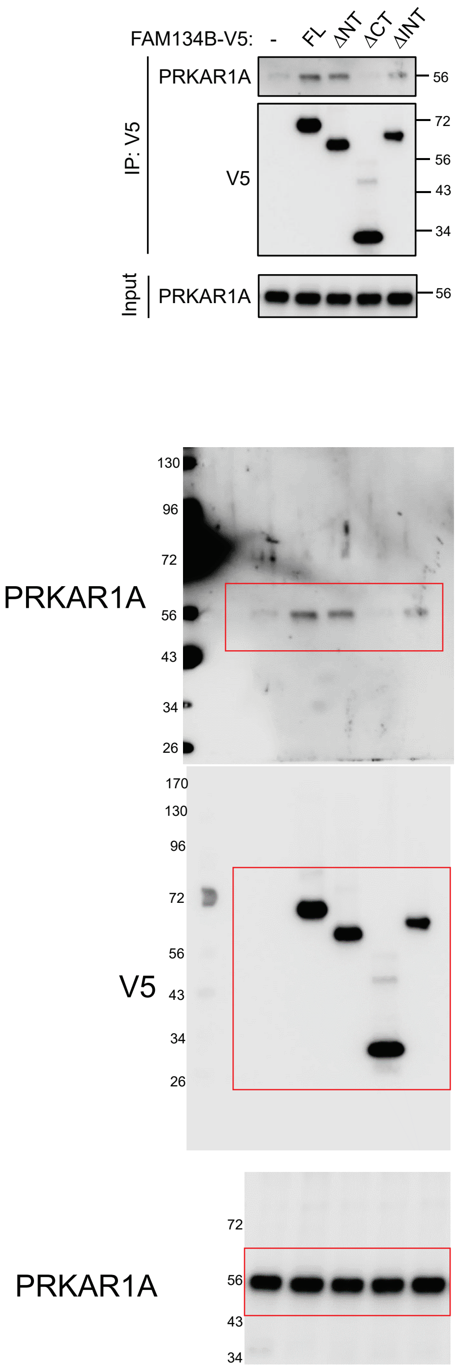


### Figure 3G

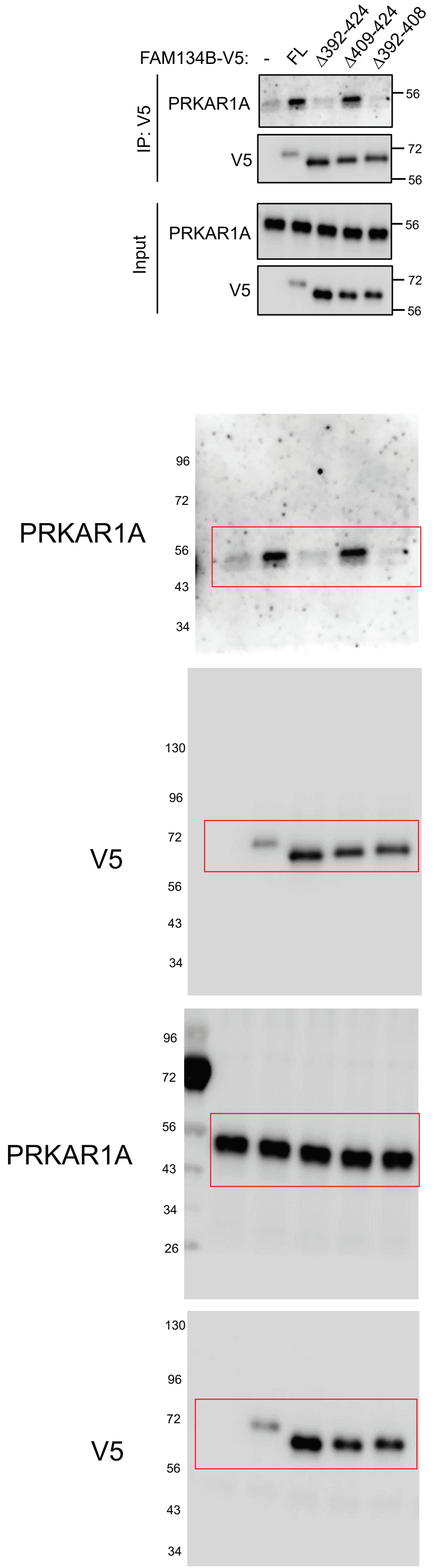

### Figure 3I

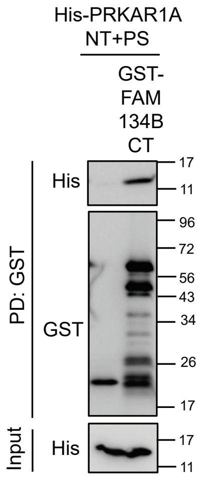


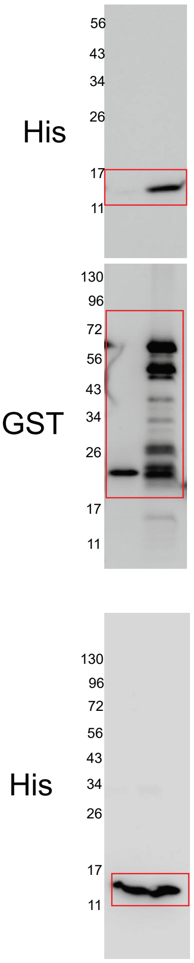


### Figure 3L

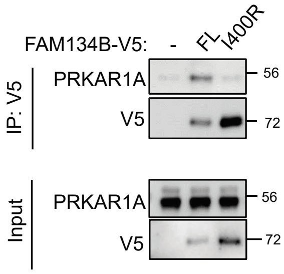


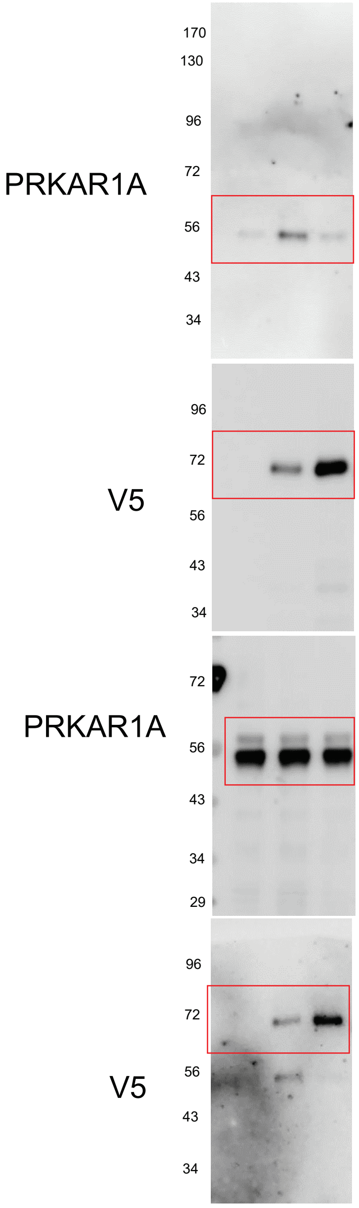

### Figure S3A

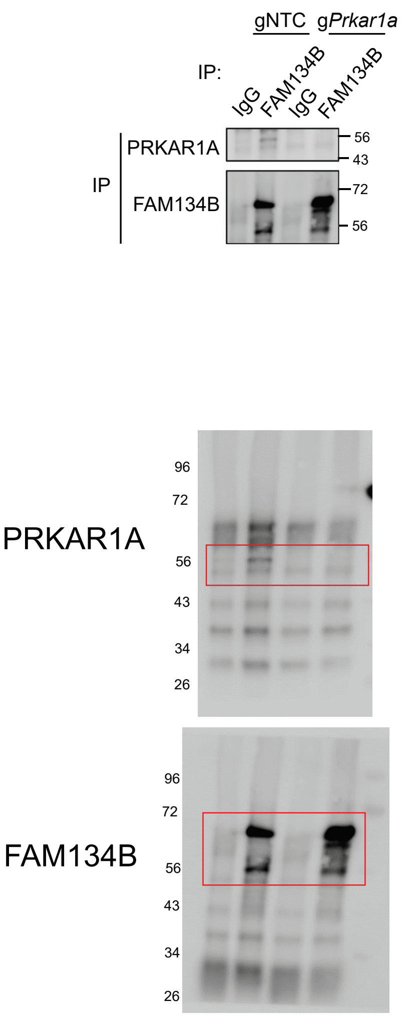


### Figure S3B

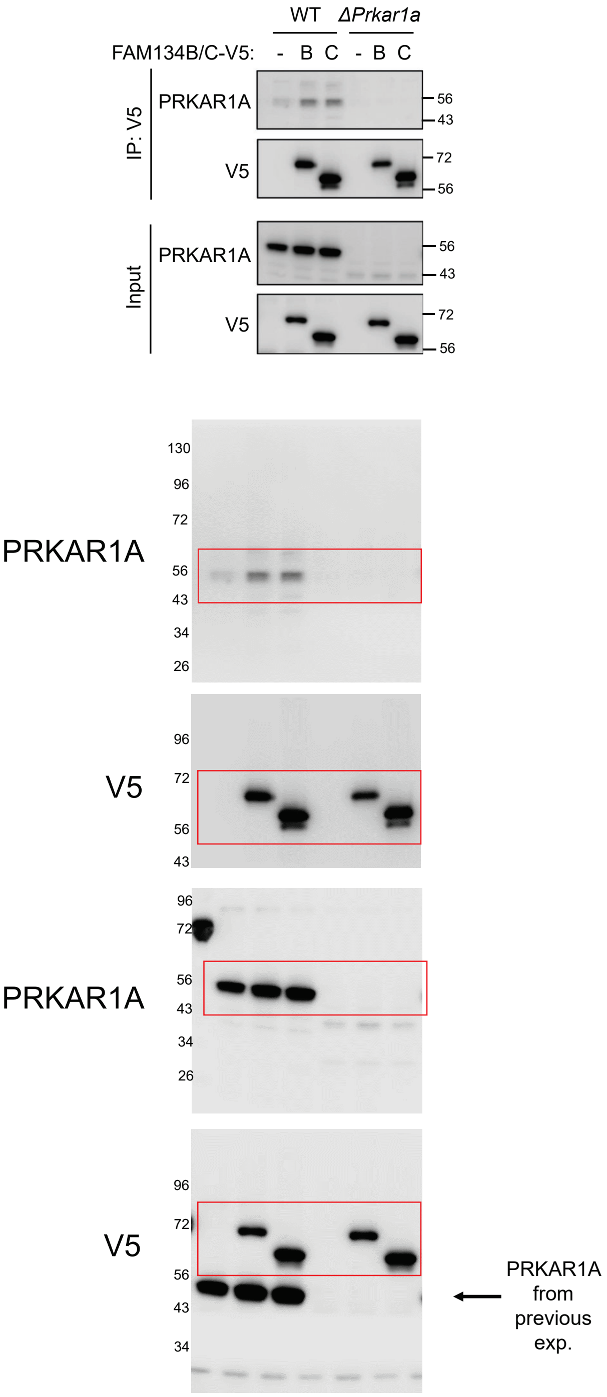

### Figure S3C

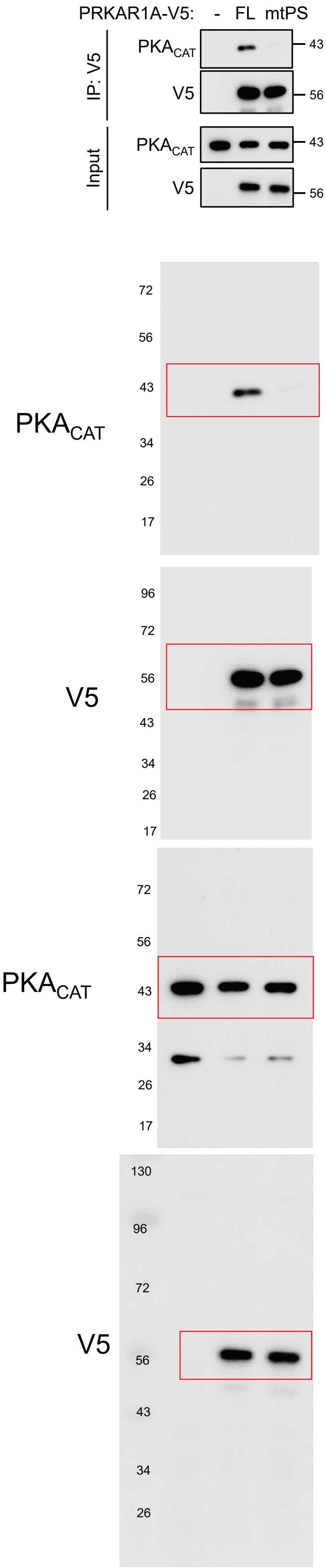


### Figure S3D

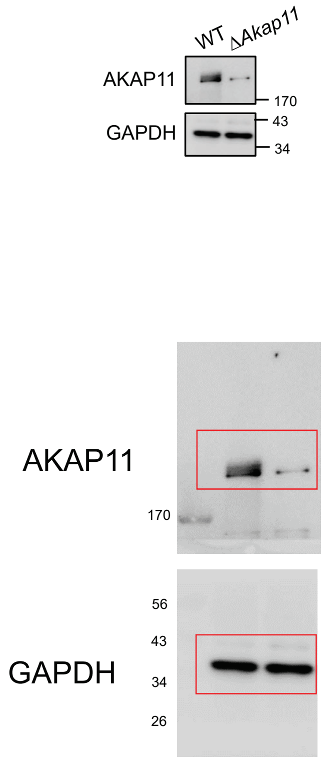

### Figure S3E

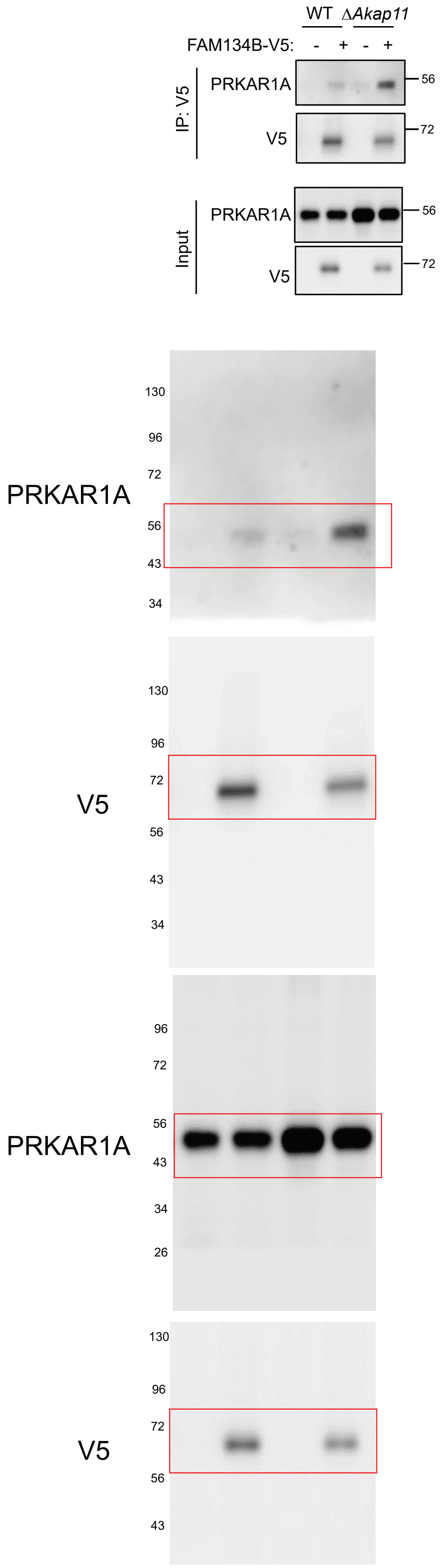


### Figure S3G

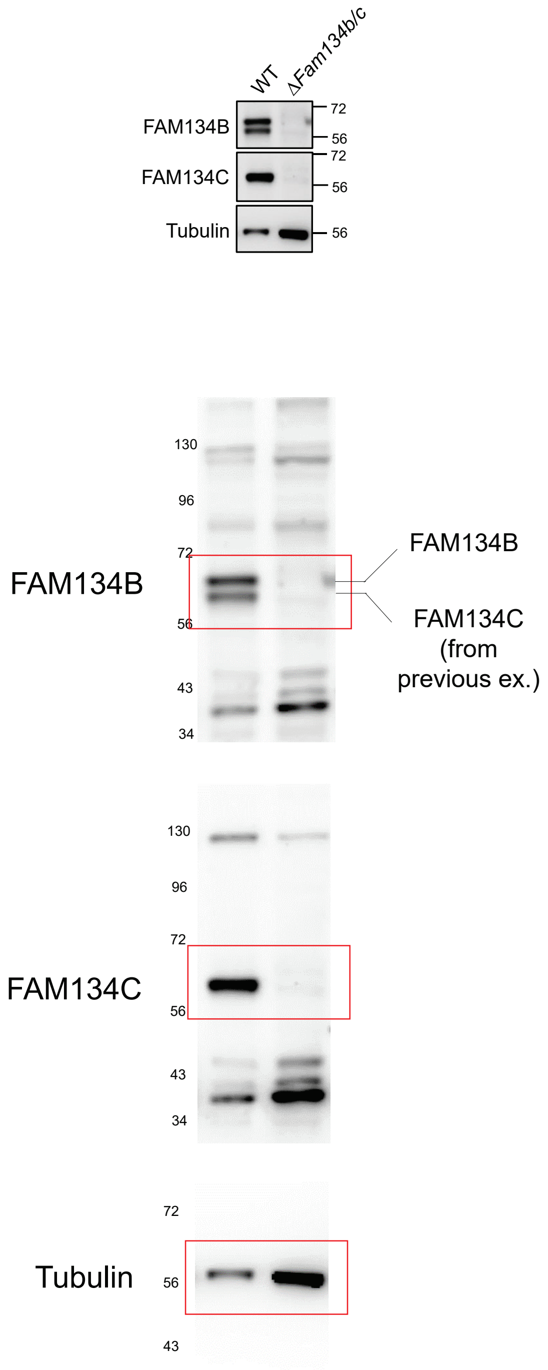

### Figure S3H

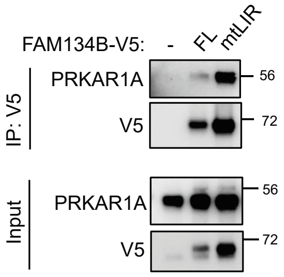
FAM134B-V5:
-
FL
mtLIR
IP: V5
PRKAR1A
V5
Input
PRKAR1A
V5
56
72
56
72

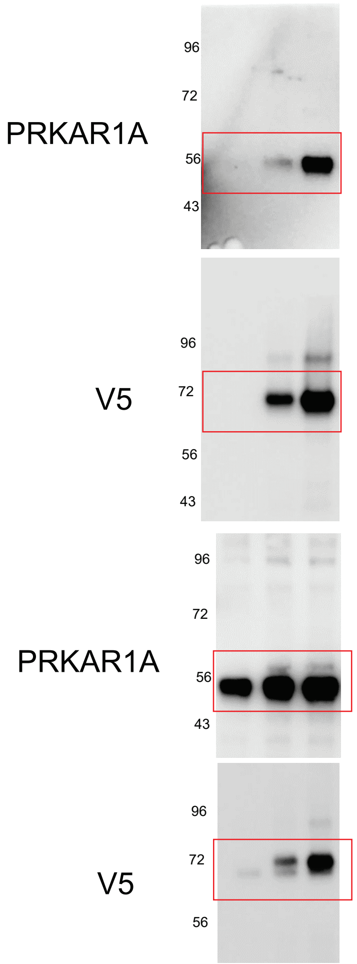
PRKAR1A
96
72
56
43
V5
96
72
56
43
PRKAR1A
96
72
56
43
V5
96
72
56

### Figure S3I

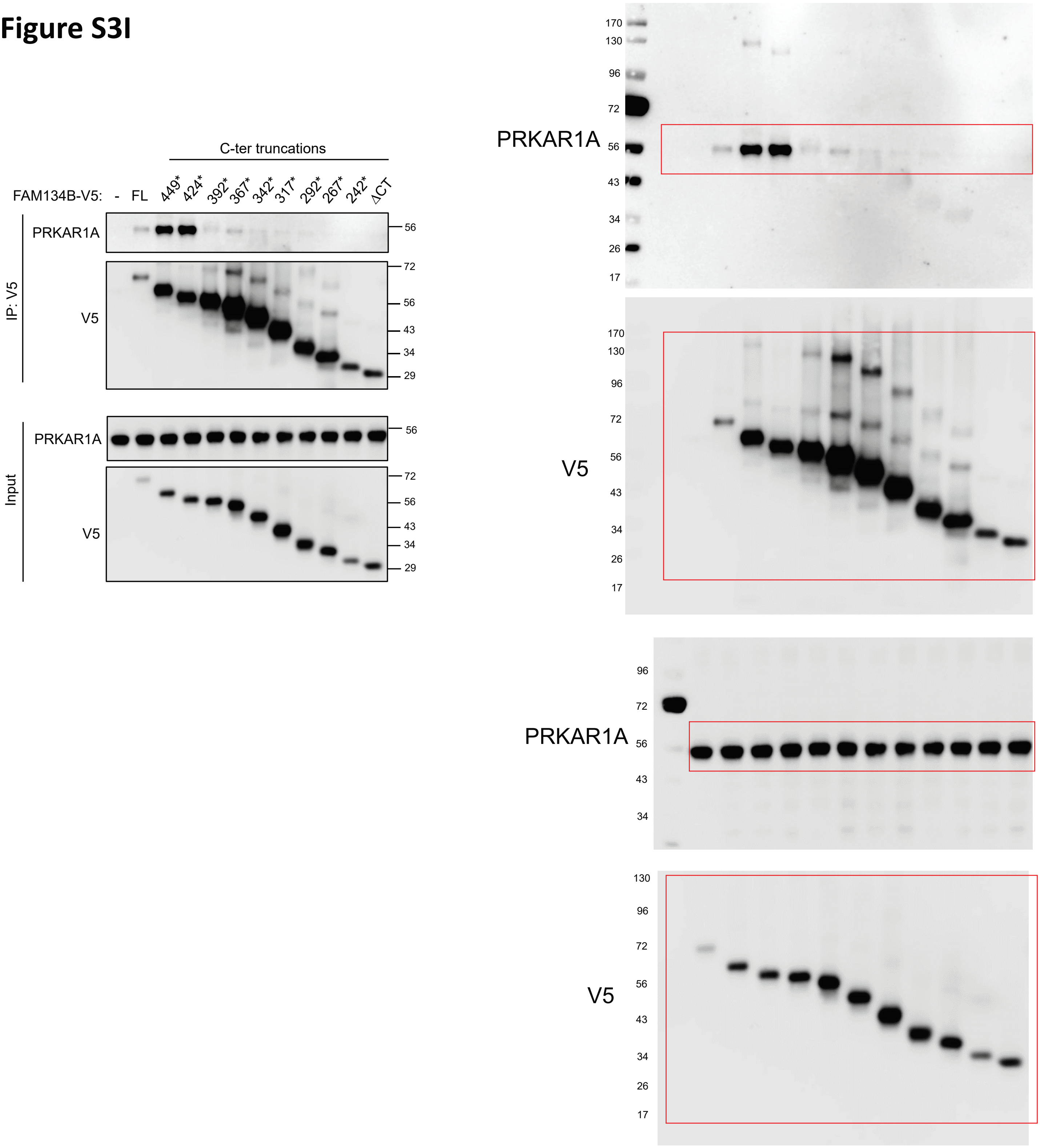
C-ter truncations
FAM134B-V5: - FL 449* 424* 392* 367* 342* 317* 292* 267* 242* ΔCT
IP: V5
PRKAR1A
V5
Input
PRKAR1A
V5
56
72
56
43
34
29
170
130
96
72
56
43
34
26
17
PRKAR1A
V5
PRKAR1A
V5

### Figure 5H

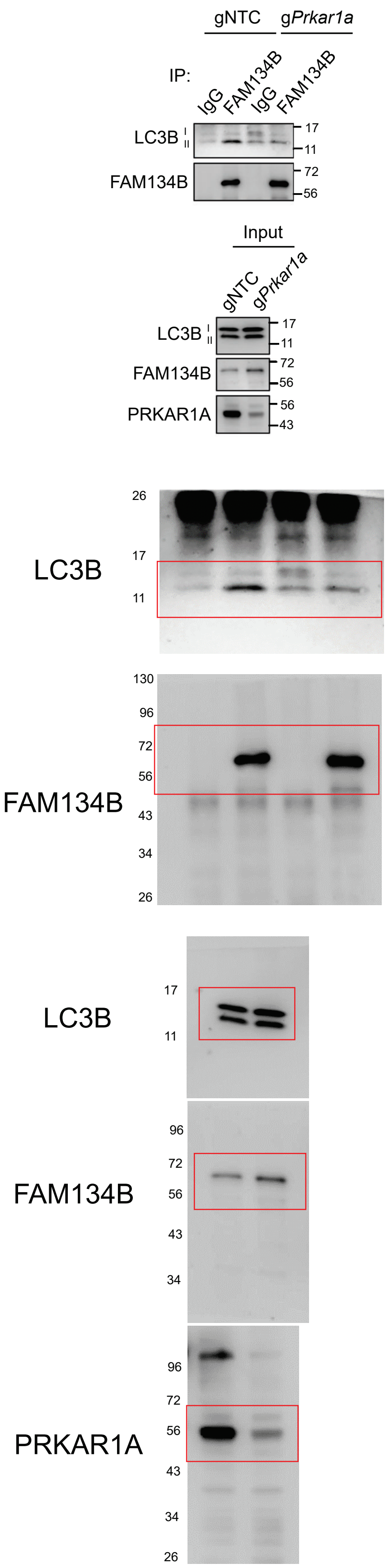


### Figure 5J

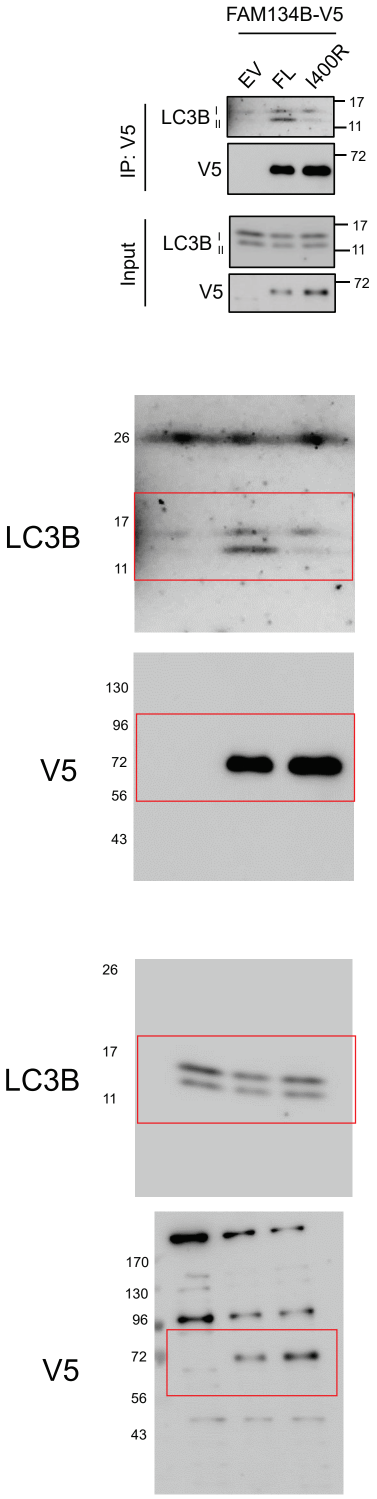

### Figure S5C

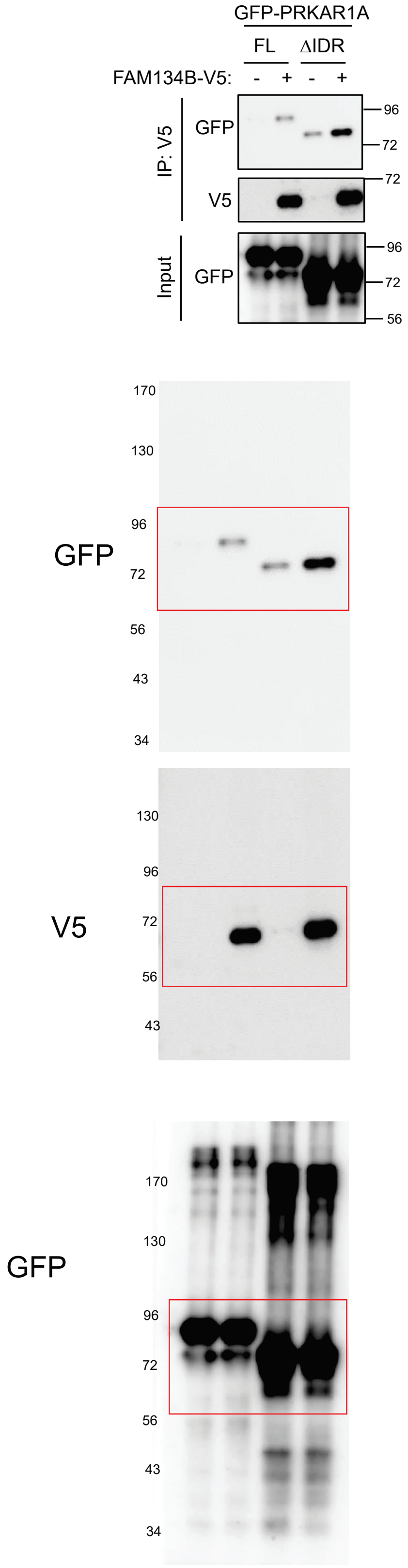


### Figure S5D

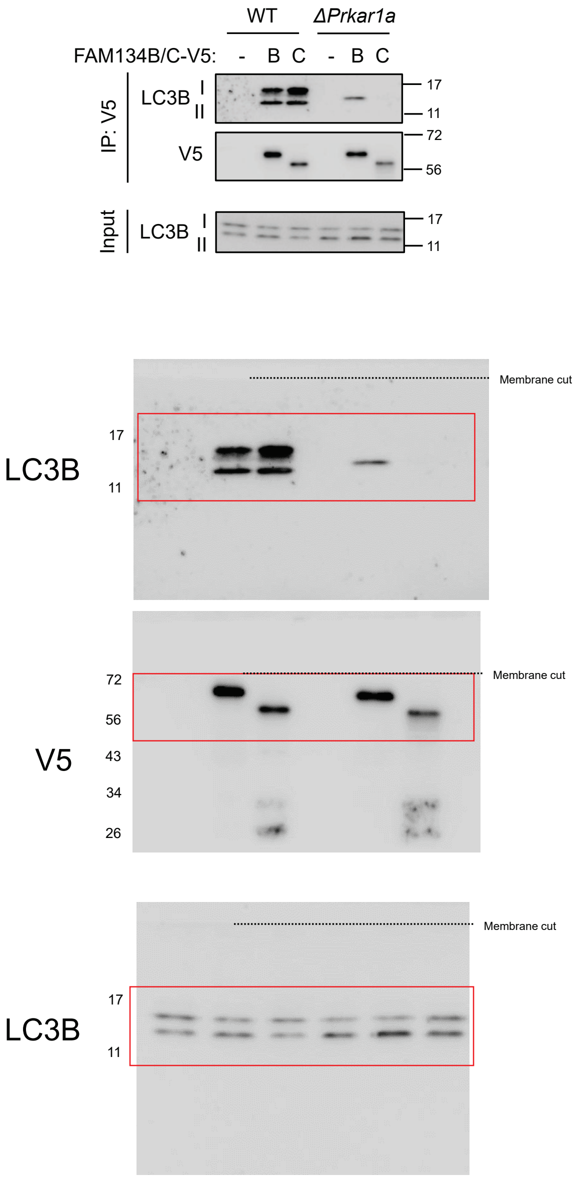

### Figure 6E

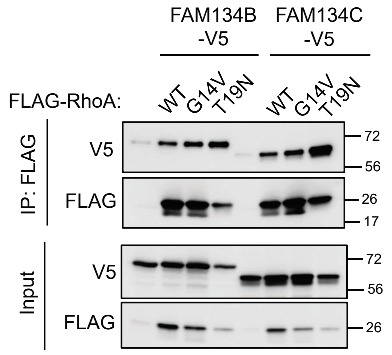

FAM134B -V5
FAM134C -V5
FLAG-RhoA:
WT
G14V
T19N
WT
G14V
T19N
IP: FLAG
V5
FLAG
Input
V5
FLAG
72
56
26
17
72
56
26


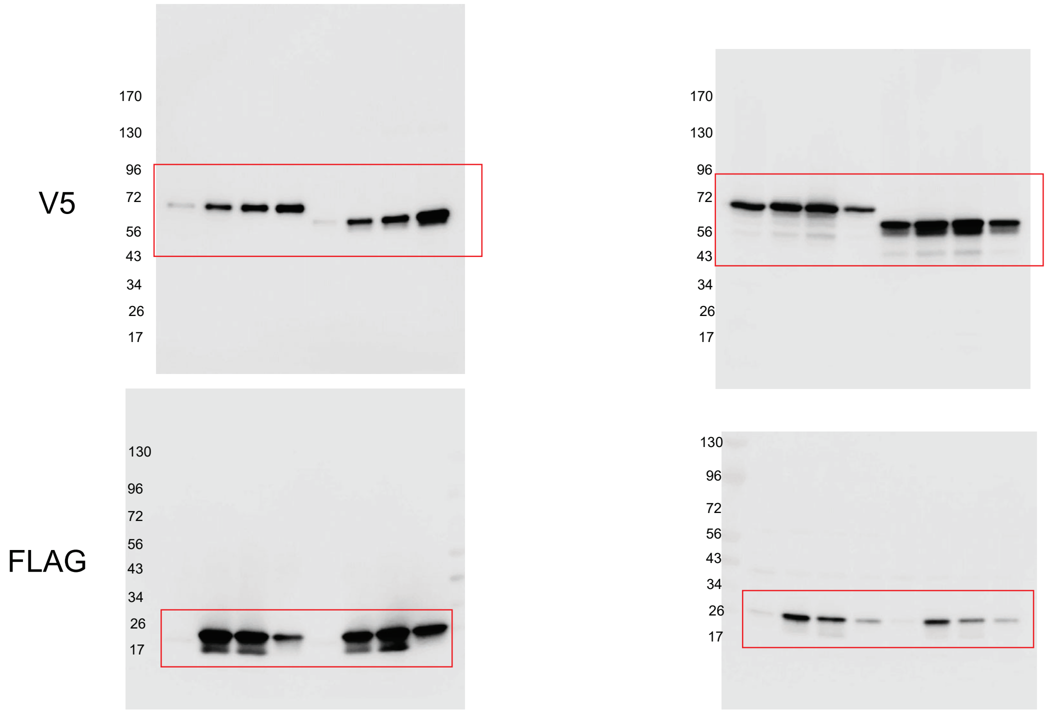

V5
FLAG
170
130
96
72
56
43
34
26
17

### Figure 6K

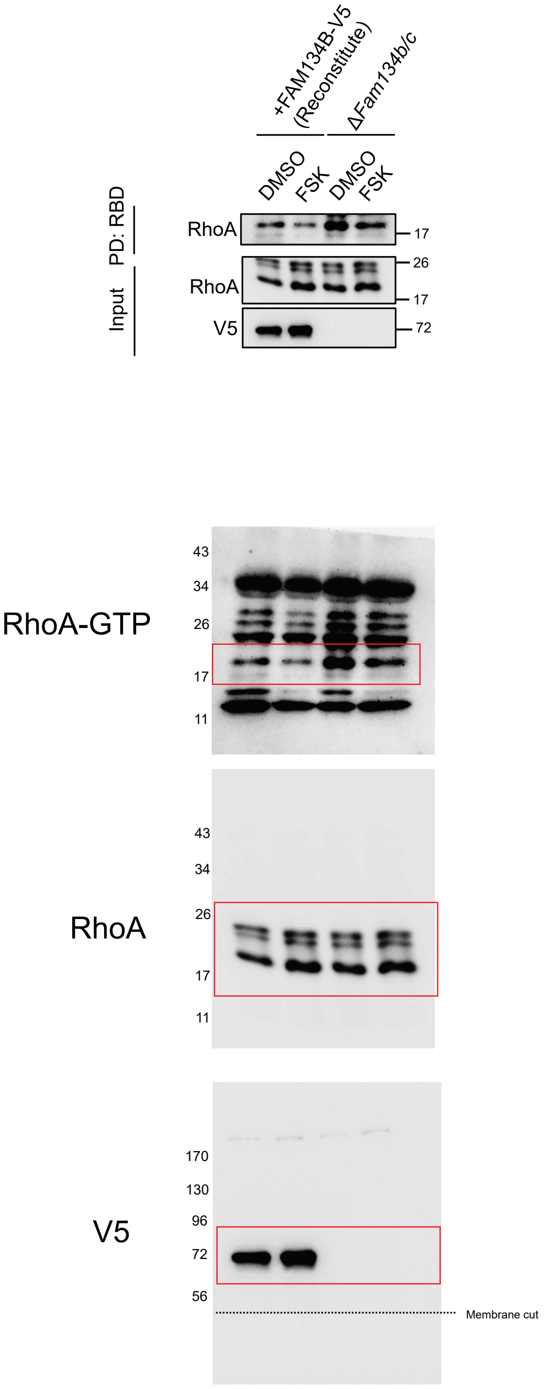


### Figure S6A

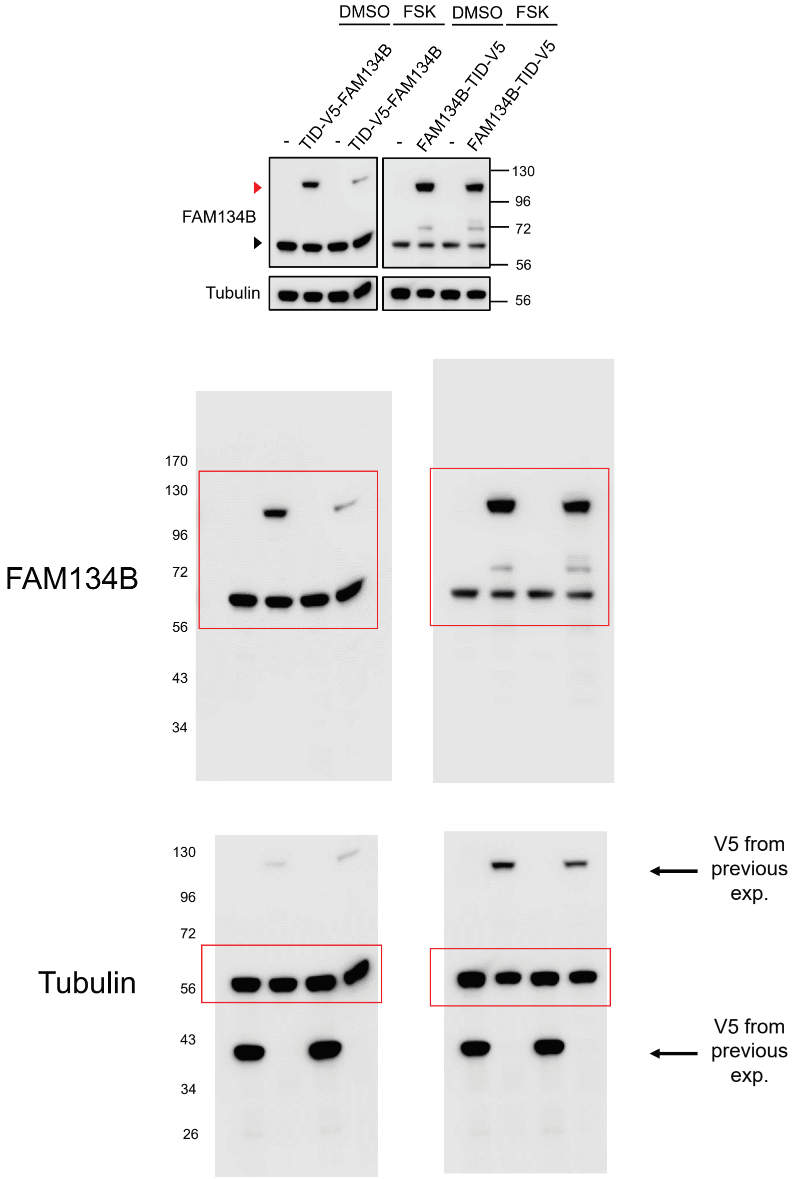

### Figure S6B

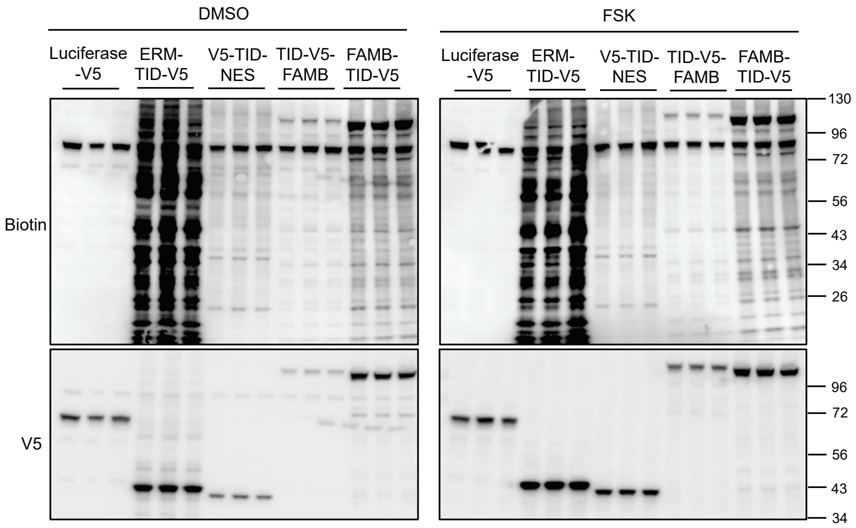
DMSO
FSK
Luciferase -V5
ERM-TID-V5
V5-TID-NES
TID-V5-FAMB
FAMB-TID-V5
Luciferase -V5
ERM-TID-V5
V5-TID-NES
TID-V5-FAMB
FAMB-TID-V5
Biotin
V5
130
96
72
56
43
34
26
96
72
56
43
34

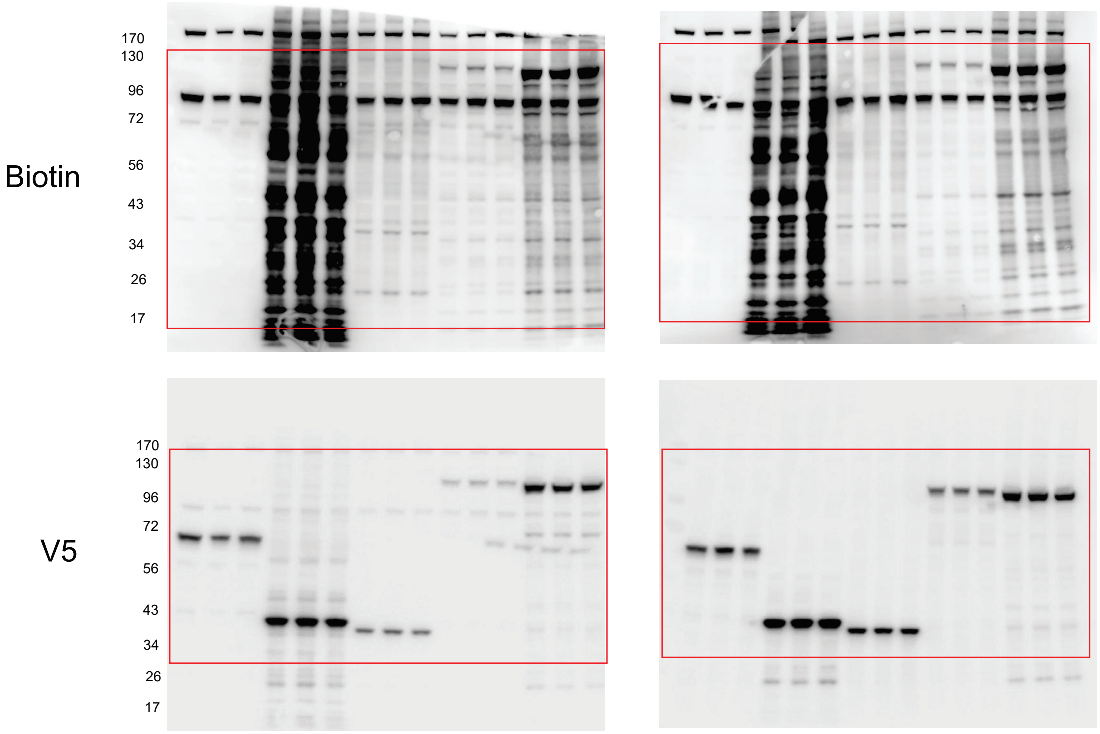
Biotin
170
130
96
72
56
43
34
26
17
V5
170
130
96
72
56
43
34
26
17

### Figure S6I

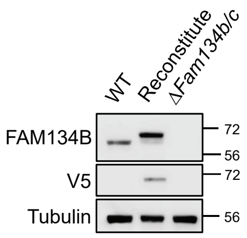
WT
Reconstitute
ΔFam134b/c
FAM134B
V5
Tubulin
72
56
72
56

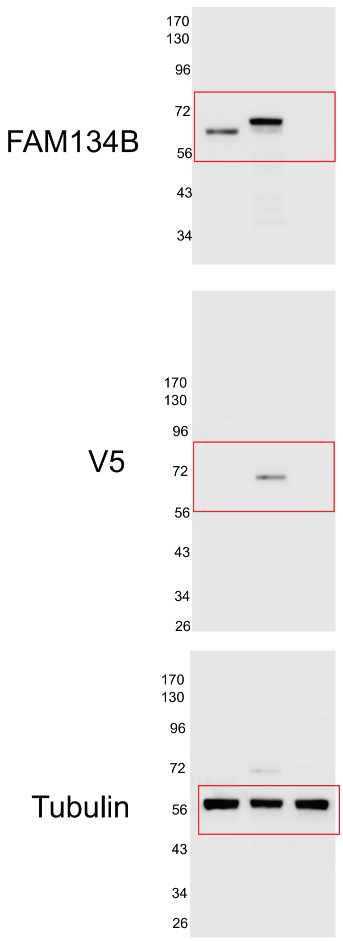
FAM134B
170
130
96
72
56
43
34
V5
170
130
96
72
56
43
34
26
Tubulin
170
130
96
72
56
43
34
26

### Figure S7A

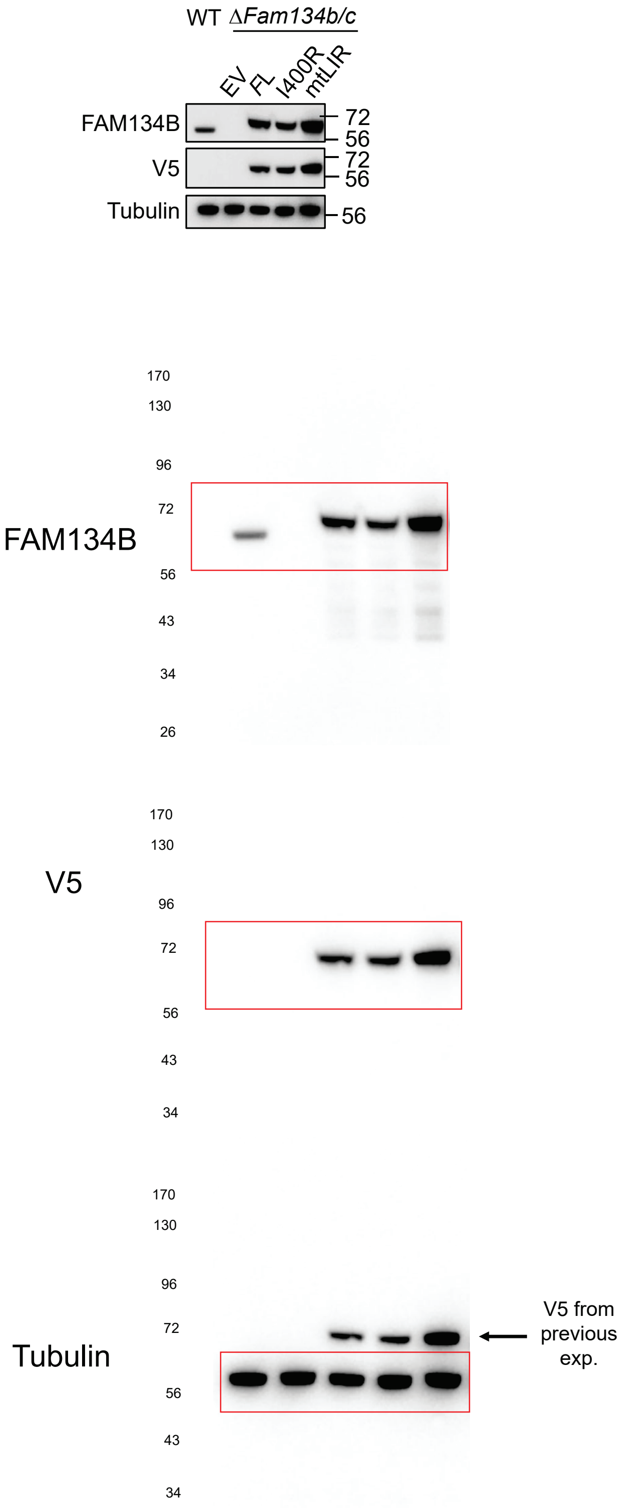


### Figure S7E

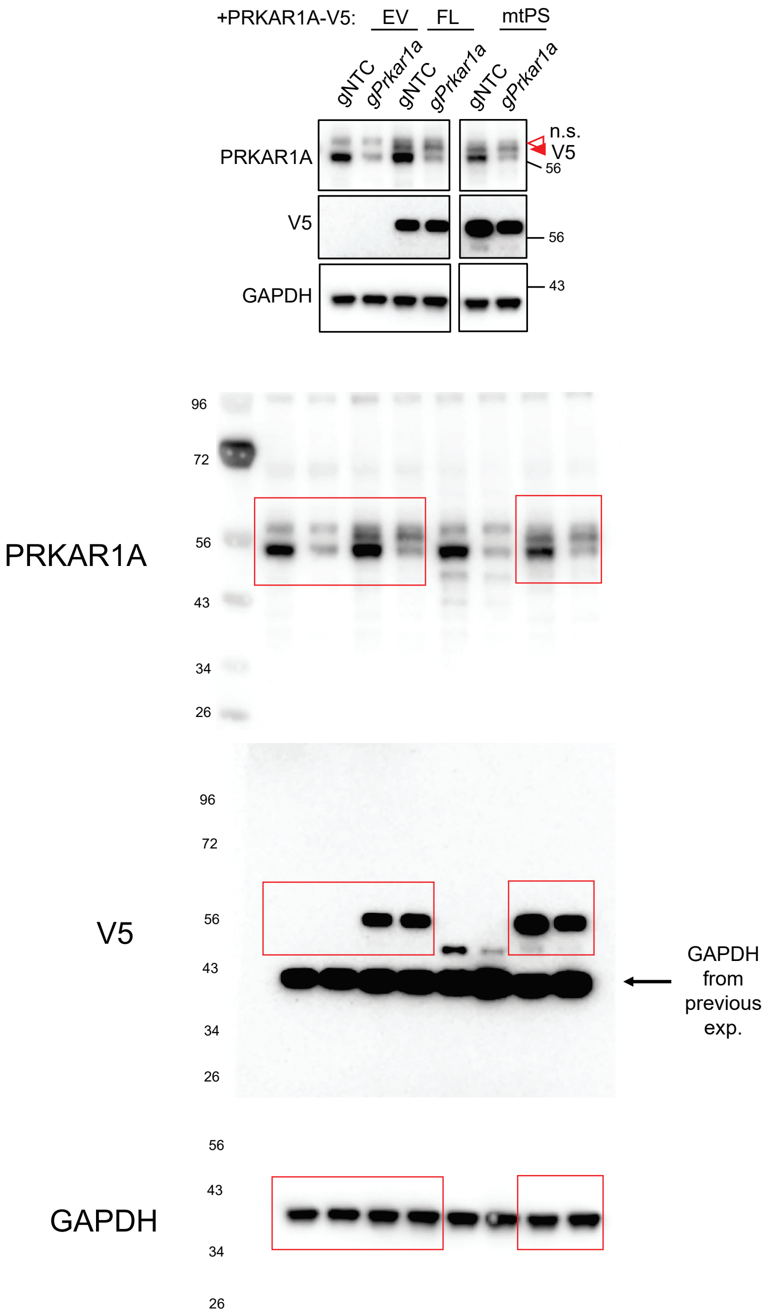

### Figure S7G

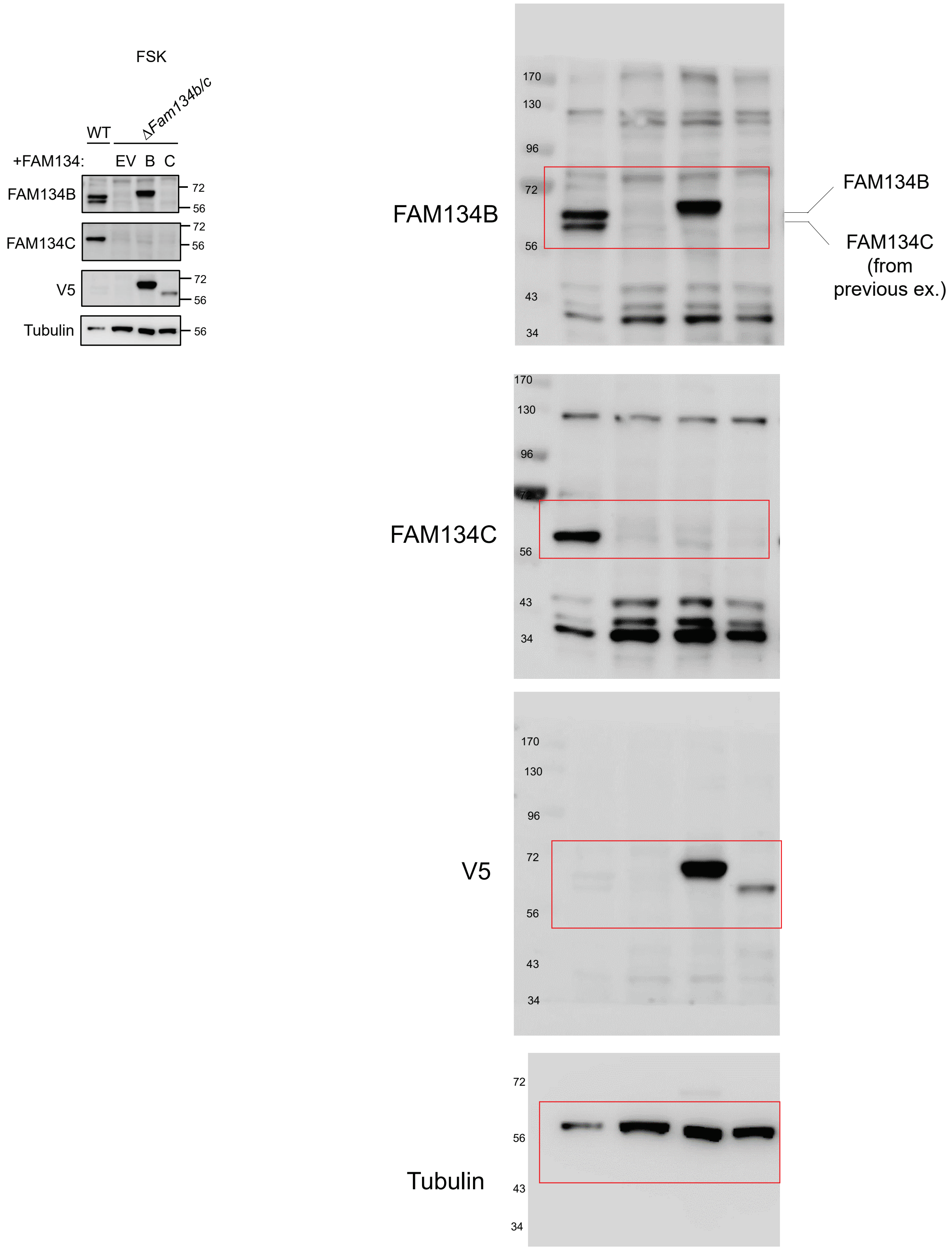

FSK
WT
ΔFam134b/c
+FAM134:
EV B C
FAM134B
FAM134C
V5
Tubulin
72
56
FAM134B
170
130
96
43
34
FAM134B
FAM134C
(from
previous ex.)
FAM134C
V5
Tubulin
