## Supplementary figures for "ER-liquid condensate contacts sequester FAM134B/C and RhoA to govern cell morphology"

#### Contents of this File

Figures S1-S7

Legends for Tables S1-S3

Legends for Movies S1-S12

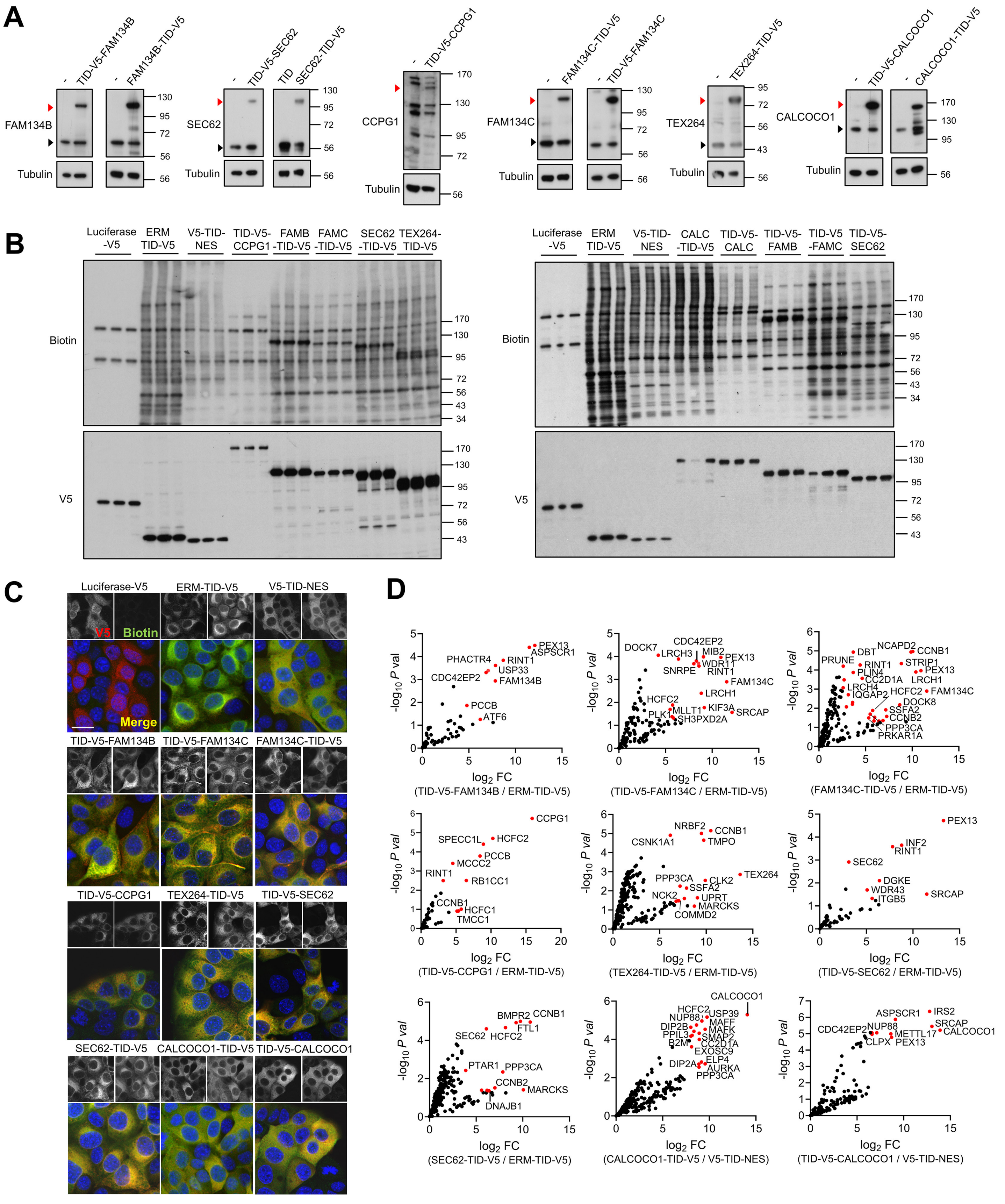


**Figure S1, related to Figure 1**

### Figure S1, related to Figure 1

**(A)** Immunoblot of PDAC cells stably-expressing TID-V5 tagged ER-phagy receptors with indicated antibodies, facilitating comparison of endogenous receptor levels (non-expressing controls indicated by -). Red arrowheads indicate stably-expressed fusion protein; black arrowheads indicate endogenous protein (N.B. CCPG1 is endogenously expressed at very low levels in cultured PDAC cells).

**(B-C)** Differential biotinylated protein abundance in the indicated PDAC stable expression cell pools after biotin supplementation, as detected by (**B**) HRP-Streptavidin blot (*n* = 3 replicates) or (**C**) Streptavidin-FITC/anti-V5 costaining and confocal immunofluorescence microscopy (representative of *n* = 2 independent experiments). Cells were treated with 0.5 mM Biotin for varying times to reduce variability in total biotinylation levels: 1 h (FAM134B-TID-V5, FAM13C-TID-V5, SEC62-TID-V5, TEX264-TID-V5, TID-V5-CALCOCO1, TID-V5-FAM134B, TID-V5-FAM134C, TID-V5-SEC62); or 4 h (Luciferase-V5, ERM-TID-V5, V5-TID-NES, TID-V5-CCPG1, CALCOCO1-TID-V5). In (**B**) FAM134B abbreviated FAMB; FAM134C abbreviated FAMC; CALCOCO1 abbreviated CALC. Scale bar, 20 µm.

**(D)** Full set of Volcano plots corresponding to Fig. 1E (*n* = 3 replicates, FC = mean fold change, *P* val = *P* value, LIMMA analysis). Most enriched high-confidence hits are highlighted in red (criteria for each bait described in detail in Materials and Methods).

MW markers, kDa.

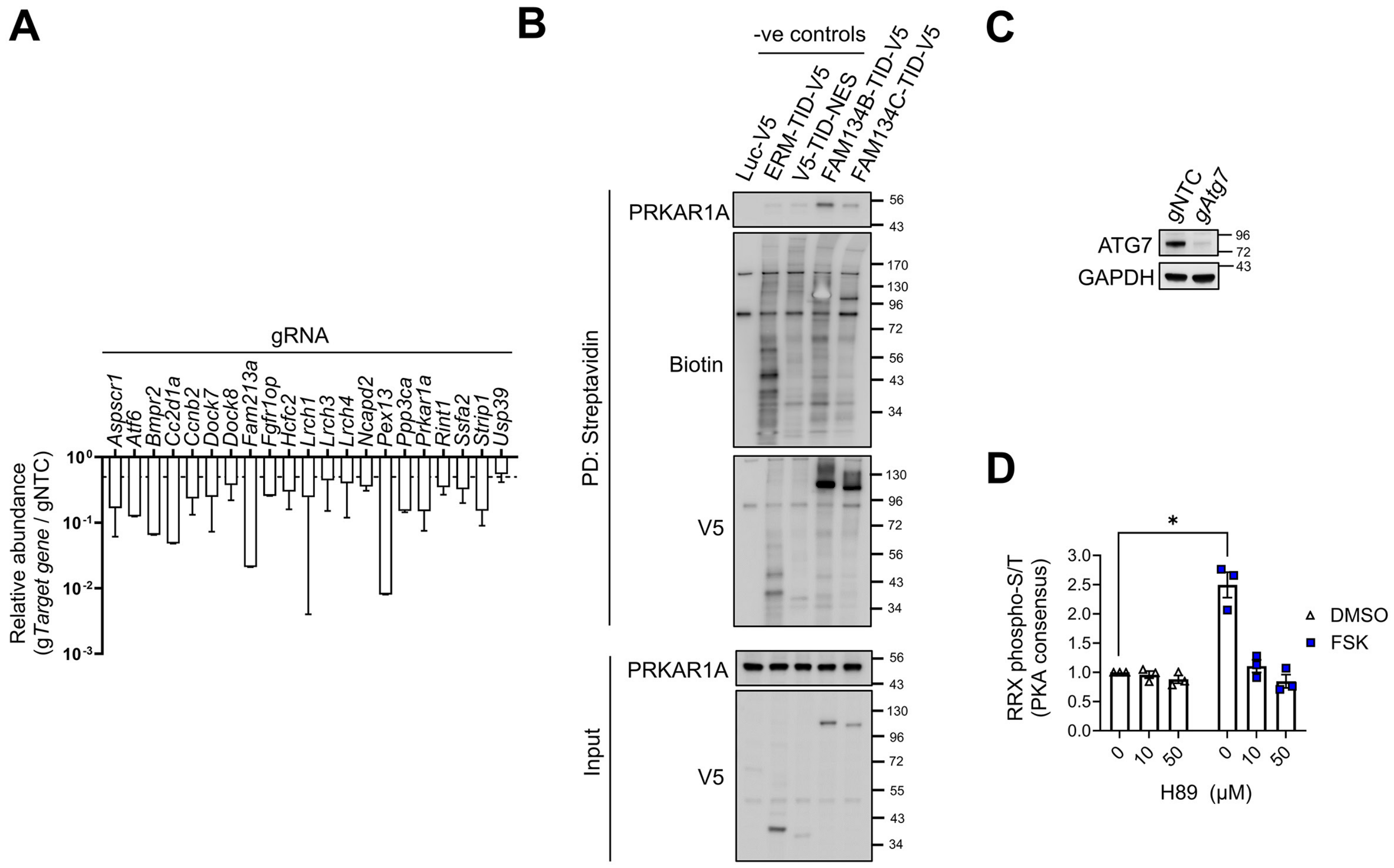


### Figure S2 related to Figure 2

**(A)** gRNA validation via qRT-PCR after PDAC-Cas9 cells were transfected with the corresponding gRNAs for 6 days (*18S*-normalised values divided by gNTC, non-targeting control gRNA, *n* = 2 independent replicates ± s.d.).

**(B)** Streptavidin pull-down (PD) from PDAC cells stably-expressing the indicated constructs and treated with Biotin (0.5 mM) for 1 h (FAM134B/C-TID-V5 fusions) or 4 h (controls), PD and 1/50 input blotted for detection of biotin and V5 as per Fig. 1C and for PRKAR1A (representative of *n* = 2 independent experiments). MW markers, kDa.

**(C)** Immunoblot of ATG7 levels in PDAC-Cas9 cells transfected with gNTC or g*Atg7* gRNAs (*n* = 3 independent replicates, representative blot shown).

**(D)** Quantification of Fig. 2G. Samples were normalised to DMSO (no H89) (*n* = 3 independent replicates, ± s.e.m., 1-sample 2-tailed t-test DMSO vs. forskolin, FSK, * = $P \leq 0.05$).

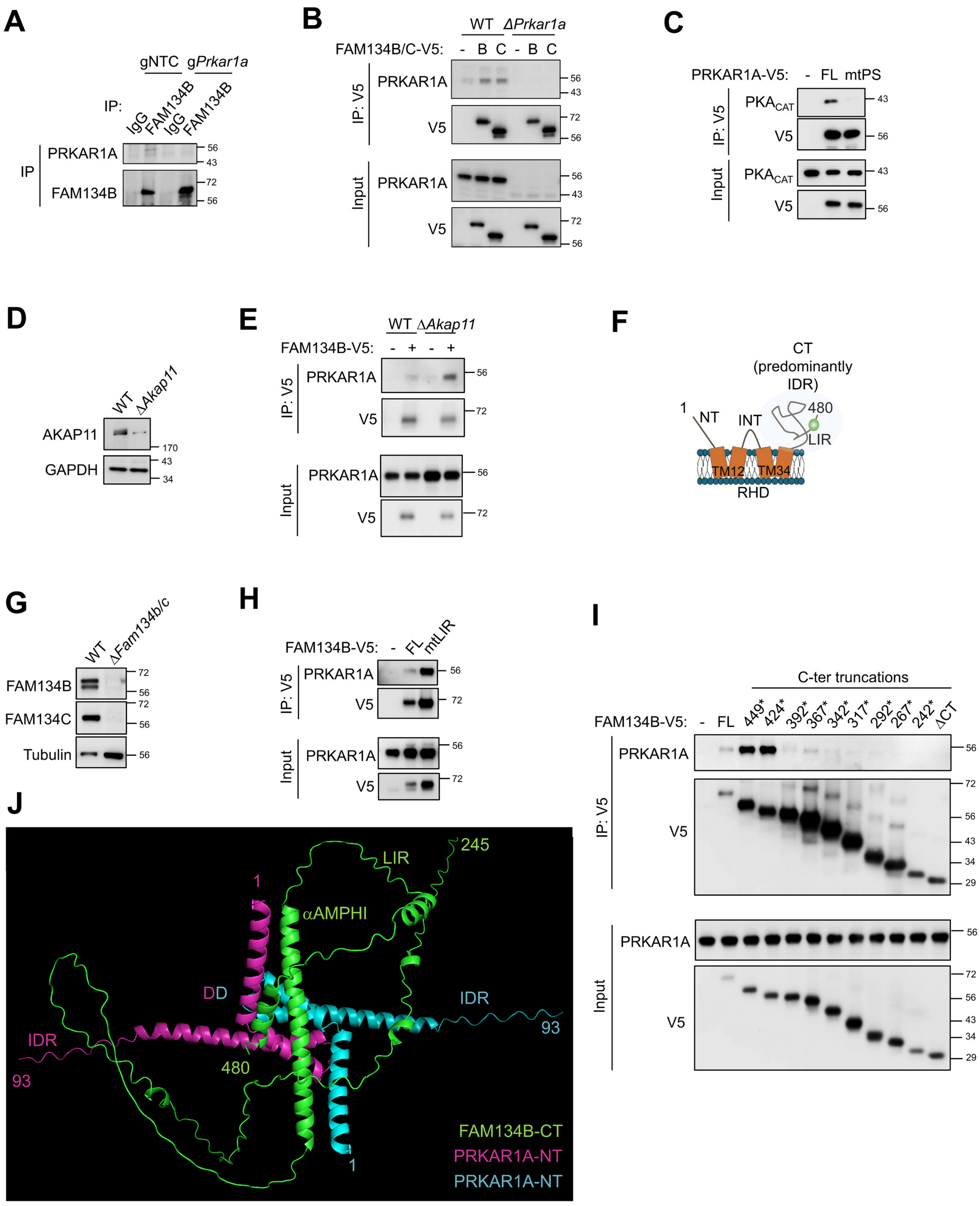


**Figure S3 related to Figure 3**

### Figure S3 related to Figure 3

**(A)** Validation of PRKAR1A signal in endogenous FAM134B immunoprecipitation (IP), performed as per Fig. 3B, from PDAC-Cas9 cells transfected with NTC (non-targeting control) or *Prkar1a* gRNA (*n* = 3 independent replicates, representative blot shown).

**(B)** Immunoprecipitation (IP) of FAM134B-V5 (B) or FAM134C-V5 (C) from stably-expressing parental PDAC (WT) or Δ*Prkar1a* clone, or cognate empty vector transduced controls (-), IP and 1/50 input immunoblotted for endogenous PRKAR1A and V5 (*n* = 2 independent replicates, representative blot shown).

**(C)** Immunoprecipitation (IP) of PRKAR1A-V5 (FL vs. mtPS mutant) from stably-expressing PDAC cells, or empty vector controls (-), IP and 1/50 input immunoblotted for endogenous $PKA_{CAT}$ (*n* = 2 independent replicates, representative blot shown). mtPS fails to interact with $PKA_{CAT}$ as expected.

**(D)** Immunoblot validation of knockout in Δ*Akap11* clone vs. parental (WT) PDAC-Cas9 cells (*n* = 3 independent replicates, representative blot shown).

**(E)** Cross-linking immunoprecipitation (IP) of FAM134B-V5 from stably-expressing PDAC WT or Δ*Akap11* clone, or cognate empty vector controls (-) and input 1/50, immunoblotted for endogenous PRKAR1A (*n* = 2 independent replicates, representative blot shown).

**(F)** Schematic of full length FAM134B protein (murine numbering). Two membrane inserted segments (orange, belonging to the reticulon homology domain, RHD) are separated by the cytoplasmic INT region with N- and C-terminal cytoplasmic domains either side. The C-terminal domain (CT) is predominantly intrinsically disordered and encompasses the LIR (LC3 interacting region) motif.

**(G)** Immunoblot validation of knockout of FAM134B and FAM134C in Δ*Fam134b/c* vs. parental (WT) PDAC-Cas9 cells (*n* = 3 independent replicates, representative blot shown).

**(H)** Cross-linking IP of FAM134B-V5 from stably-reconstituted Δ*Fam134b/c* PDAC cells (FL vs. mtLIR) or empty vector control (-), IP and 1/50 input immunoblotted for endogenous PRKAR1A and V5 (*n* = 3 independent replicates, representative blot shown).

**(I)** Cross-linking IP of FAM134B-V5 from stably-reconstituted Δ*Fam134b/c* PDAC cells (FL vs. indicated C-terminal truncations; * = fused with C-terminal V5 tag after indicated amino acid) or empty vector control (-), IP and 1/50 input immunoblotted for endogenous PRKAR1A and V5 (*n* = 2 independent replicates, representative blot shown).

**(J)** Alphafold 3 prediction of mouse FAM134B CT (green, a.a. 245-480) complexed with dimer of mouse PRKAR1A NT (magenta and cyan, each monomer a.a. 1-93) (IDR, part of intrinsically disordered region; LIR, LC3 interacting region; DD, dimerization domain; CT, C-terminal domain of FAM134B; NT, N-terminal region of PRKAR1A).

MW markers = kDa.

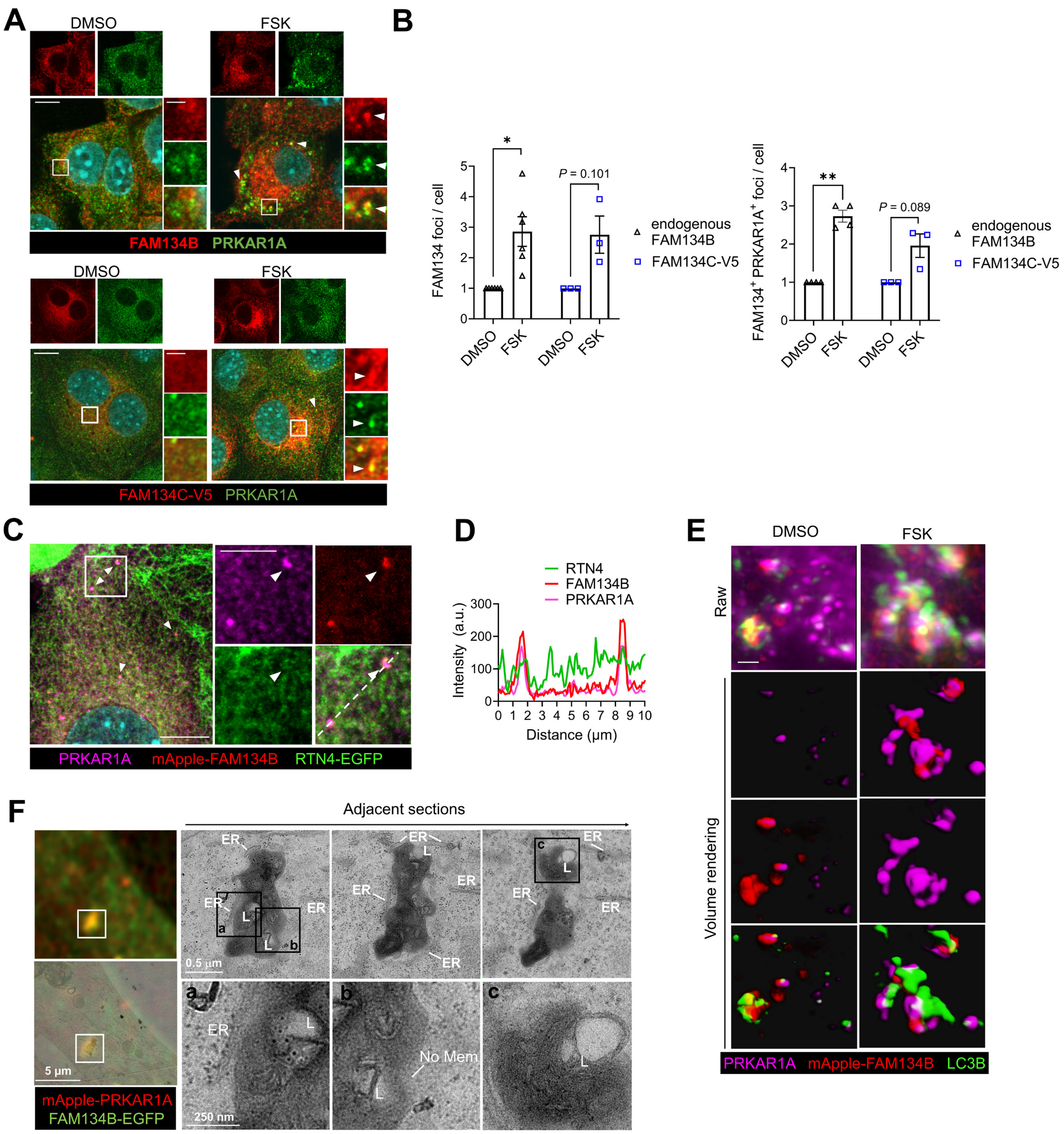


**Figure S4 related to Figure 4**

**(A-B)** Confocal immunofluorescence microscopy of PDAC cells (top panels) or PDAC cells stably-expressing FAM134C-V5 (bottom panels), treated with DMSO or forskolin, FSK (40 µM, 4 h), stained for endogenous FAM134B (top) or V5 (bottom) and PRKAR1A (all). Representative images in (**A**). Arrowheads indicate co-localising foci. Zoom panels are delineated by white boxes. Scale bars, 10 µm (main) and 2 µm (zoom). Quantifications are shown in (**B**) for endogenous FAM134B or FAM134C-V5 foci (left) or the number of these foci colocalised with PRKAR1A (right) (*n* = 3 - 6 independent replicates, > 100 cells/condition, mean ± s.e.m., 1-sample two-tailed t-tests DMSO vs. FSK, * = $P \leq 0.05$, ** = $P < 0.01$).

**(C-D)** Confocal immunofluorescence microscopy of PDAC cells stably-expressing mApple-FAM134B (red) and RTN4-EGFP (ER marker, green) treated with FSK (40 µM, 4 h), co-stained for PRKAR1A (magenta). Representative confocal fluorescence image in (**C**) and fluorescence intensity trace in (**D**) (*n* = 2 independent replicates). Zoom panel is delineated by the white box. White arrowheads indicate FAM134B-PRKAR1A foci colocalising with the RTN4 network. a.u. = arbitrary units. Scale bars, 10 µm (main), 5 µm (zoom).

**(E)** Additional example of SoRA super-resolution microscopy shown in Fig. 4E. PDAC cells stably-expressing mApple-FAM134B (magenta) treated with DMSO or FSK (40 µM, 4 h), co-stained for LC3B (cyan) and PRKAR1A (green). Left, representative raw image. Right, representative 3D reconstruction of the image with normal shading by Imaris volume rendering, each column shows different fluorescence channel combinations (*n* = 2, independent replicates). Scale bar, 1 µm.

**(F)** Correlative light electron microscopy (CLEM) of mApple-PRKAR1A and FAM134B-EGFP co-localised foci after FSK treatment (40 µM, 2 h) (see also example in Fig. 4F). Left panels, representative fluorescence image and overlay with transmission electron microscopy (EM) image. Right panels, representative EM of the zoomed image, and adjacent sections. Boxes indicate zoom panels. Scale bars, 5 µm (fluorescence), 0.5 µm (EM) and 250 nm (EM zoom). ER = endoplasmic reticulum; L = lysosome; M = mitochondrion; No mem = non-membrane delimited.

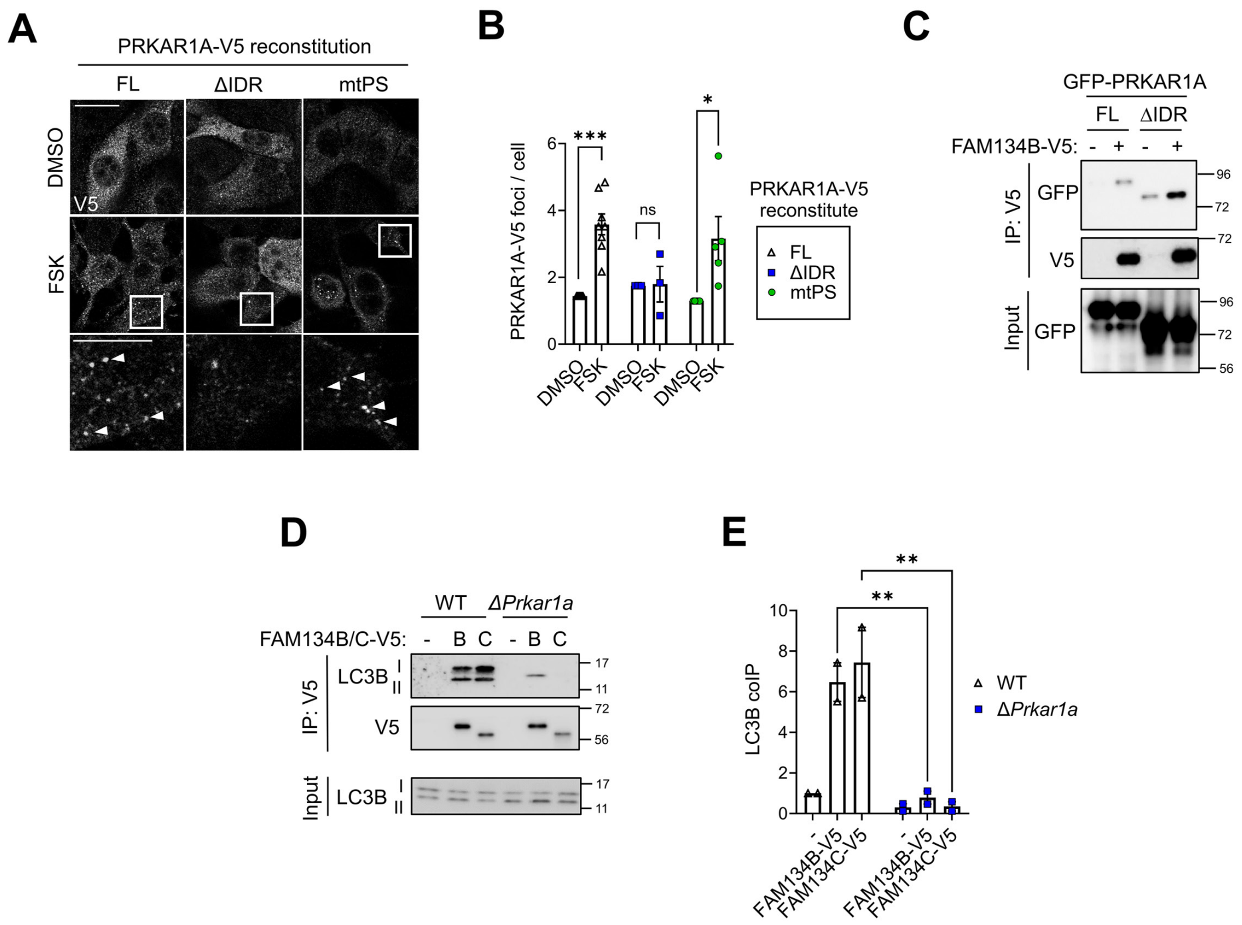


### Figure S5 related to Figure 5

**(A-B)** PDAC-Cas9 cells from Fig. 5F, representative confocal immunofluorescence images of PRKAR1A-V5 shown in (**A**). White boxes delineate zoom panels. Arrowheads in zoom indicate foci of reconstituted PRKAR1A-V5. Scale bars, 20 µm (main), 10 µm (zoom). Quantification in (**B**) of foci of reconstituted PRKAR1A-V5 (FL, wild-type, full-length; mtPS = point mutations R96A R97A preventing $PKA_{CAT}$ binding; ΔIDR, no intrinsically-disordered region required for liquid-like condensation). Each PRKAR1A variant with FSK is normalised to cognate DMSO sample (*n* ≥ 3 independent replicates, > 70 cells/condition, mean ± s.e.m., 1-sample 2-tailed t-tests, * = *P* ≤ 0.05, ns = *P* > 0.05).

**(C)** Cross-linking immunoprecipitation (IP) of FAM134B-V5 from stably-expressing PDAC cells (+) or empty vector controls (-), after transfection with indicated variants of GFP-PRKAR1A (*n* = 2 independent replicates, representative immunoblot shown).

**(D-E)** IP of FAM134B-V5 or FAM134C-V5 from stably-expressing WT PDAC or Δ*Prkar1a* PDAC cells or empty vector controls (-), immunoblotted for LC3B (B, FAM134B; C, FAM134C). Representative immunoblot in (**D**) and quantification in (**E**) of ratio of LC3B-II (lipidated LC3B) immunoprecipitated to input, normalised to control condition (WT + empty vector [-]) (*n* = 2 independent replicates, mean ± s.e.m., 2-way ANOVA and indicated Holm-Šídák multiple comparisons, ** = *P* < 0.01).

MW markers = kDa.

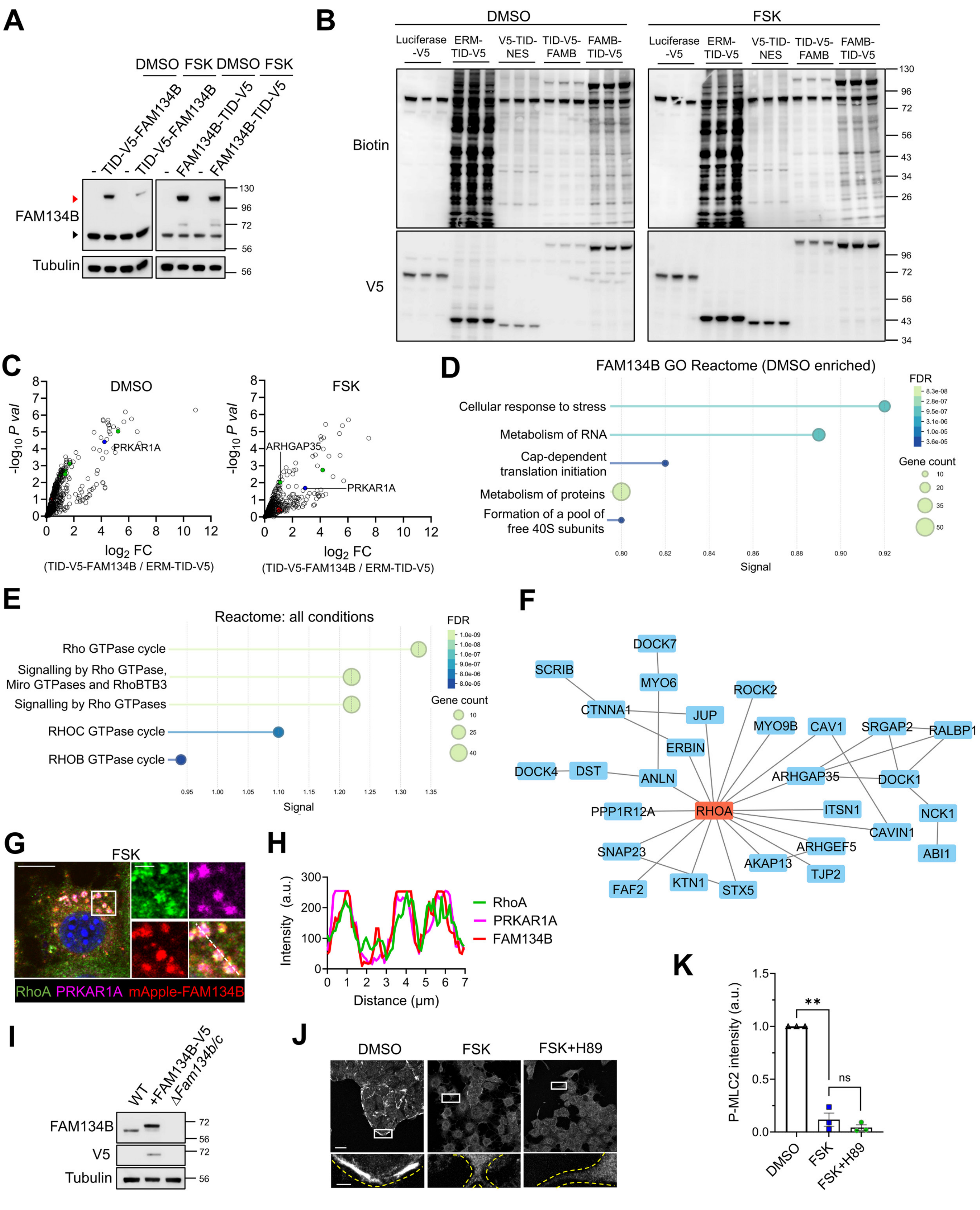


**Figure S6 related to Figure 6**

### Figure S6 related to Figure 6

In **(A-F)** PDAC cells stably-expressing FAM134B-TurboID-V5 or TurboID-V5-FAM134B (or controls ERM-TurboID-V5, V5-TurboID-NES or Luciferase-V5) were treated with DMSO or forskolin, FSK (40 µM, 4 h). Biotinylation was induced with Biotin (0.5 mM) for 4 h or 2 h (FAM134B-TID-V5 only). TID = TurboID.

**(A)** Representative immunoblots probing for FAM134B facilitating comparison of endogenous receptor levels (non-expressing controls indicated by -). Red arrowheads indicate stably-expressed FAM134B fusion; black arrowheads indicate endogenous FAM134B.

**(B)** Immunoblot for biotinylated protein abundance in the indicated stably-expressing cell pools after biotin supplementation, as detected by HRP-Streptavidin immunoblot. FAMB = FAM134B.

**(C)** Volcano plots for enriched biotinylated proteins after proximity proteomics for TID-V5-FAM134B. Shown are proteins remaining after *P*-value cut-off versus TID-NES controls, plotted for enrichment vs. ERM-TID ($n$ = 3 independent replicates, FC = mean fold change, *P* val = *P* value, LIMMA analysis). See also Table S3. Red highlights FSK-enriched prey proteins from within enriched Reactomes (Fig. 6B). Green highlights additional proteins from within these Reactomes present in analysis of all conditions (insensitive to FSK status, Fig. S6E) ($n$ = 3 technical replicates).

**(D)** Reactome Pathway enrichment analysis of prey hits enriched in DMSO-stimulated FAM134B-TID-V5 proximity labelling proteomics versus FSK control (see also Fig. 6A-B). FDR = false discovery rate.

**(E)** Reactome Pathway enrichment analysis of prey hits enriched in all conditions versus negative controls (ERM-TurboID-V5 and V5-TurboID-NES, see Methods). FDR = false discovery rate.

**(F)** Physical interaction network of FAM134B-proximal proteins in red and green in Volcano plots in Fig. 6A (STRING analysis, see Methods), trimmed to show the central network around RhoA.

**(G-H)** Immunofluorescence confocal microscopy of PDAC cells stably-expressing mApple-FAM134B (red) treated with forskolin, FSK (40 µM, 4 h), co-stained for RhoA (green) and PRKAR1A (magenta). Representative image in (**G**). White box indicates zoom panel. Scale bars, 10 µm (main), 2 µm (zoom). Fluorescence intensity trace along the broken line in (**H**) ($n$ = 3 replicates). a.u. = arbitrary units.

**(I)** Representative immunoblot for samples from Fig. 6K for level of FAM134B-V5 expression in reconstituted Δ*Fam134b/c* PDAC cells (+FAM134B-V5) relative to endogenous FAM134B in WT (wild-type) parental cells ($n$ = 3 independent replicates).

**(J-K)** Immunofluorescence confocal microscopy of PDAC cells treated with DMSO or FSK (40 µM, 2 h) +/- the PKA inhibitor H89 (20 µM), stained for phospho-S18/T19-myosin light chain 2 (P-MLC2). Representative images in (**J**), white box delineates zoom panel, yellow lines indicate cell boundaries. Scale bars, 20 µm (main), 5 µm (zoom). Quantifications in (**K**) of extranuclear P-MLC2 intensity in cells, normalised to DMSO, a.u. = arbitrary units ($n$ = 3 independent replicates, > 200 cells/condition, mean ± s.e.m., 1-sample 2-tailed t-test DMSO vs. FSK, 2 tailed t-test FSK vs. FSK+H89, ** = $P < 0.01$, ns = $P > 0.05$).

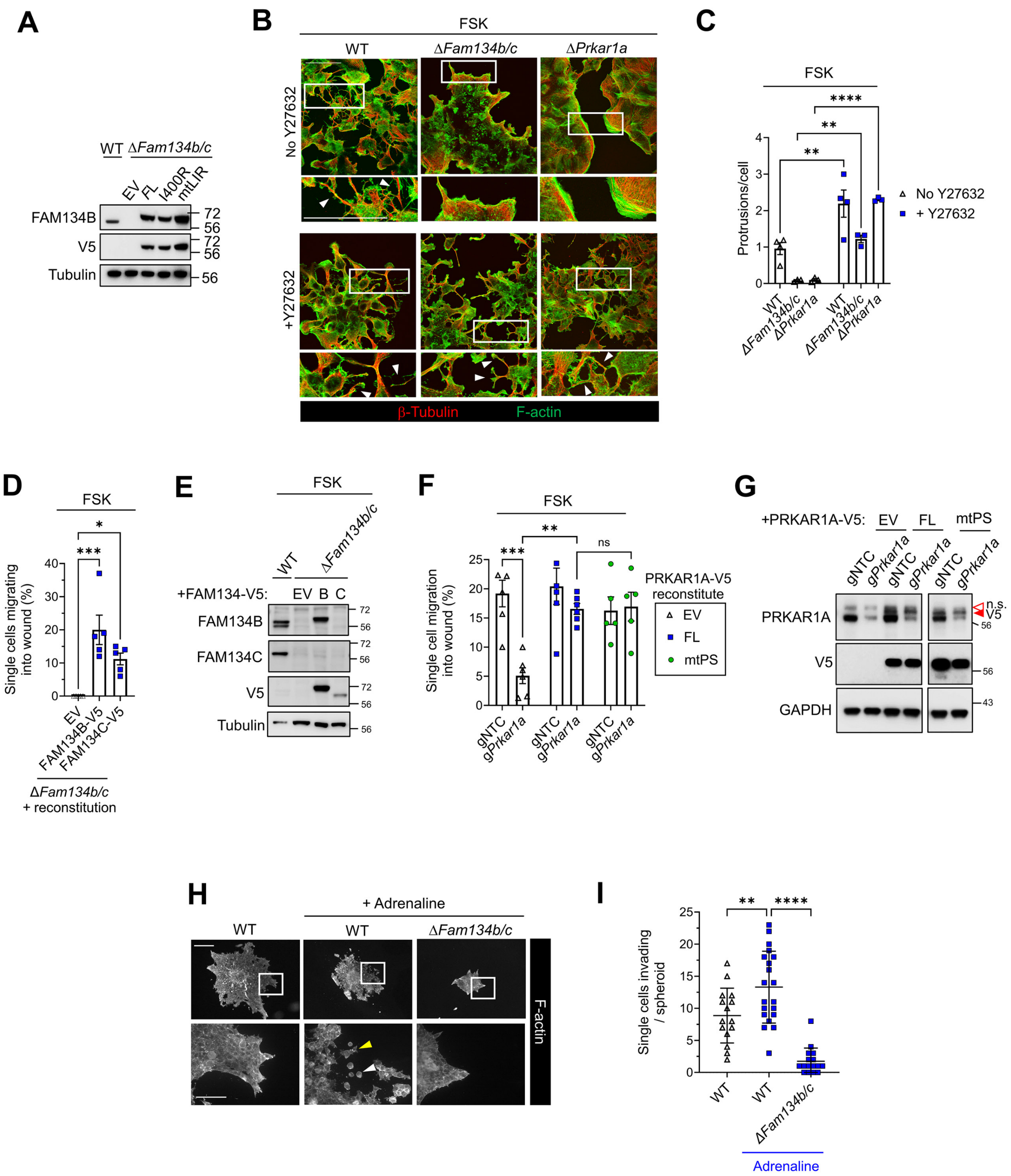


Figure S7 related to Figure 7

**Figure S7 related to Figure 7**

**(A)** FAM134B-V5 expression relative to endogenous levels (WT, wild-type cells) in Δ*Fam134b/c* PDAC cells reconstituted with indicated variants of FAM134B-V5. Representative immunoblot shown (*n* = 2 independent replicates).

**(B-C)** Confocal immunofluorescence microscopy of PDAC WT, Δ*Prkar1a* or *ΔFam134b/c* cells treated with forskolin (FSK) (40 µM, 2 h) after pre-treatment with either DMSO vehicle (no Y27632) or 20 µM ROCK inhibitor (+Y27632) (20 µM). Representative images shown in (**B**). Cells co-stained for F-actin (phalloidin, green) and β-Tubulin (red). Arrowheads indicate cellular protrusions. White boxes delineate zoom panels. Scale bars, 50 µm. Quantifications in (**C**) for protrusion formation driven by FSK (*n* = ≥ 3 independent replicates, mean ± s.e.m, 2-way ANOVA and indicated Holm-Šídák multiple comparisons, **** = p ≤ 0.0001, ** = p ≤ 0.01).

**(D-E)** 2D scratch wound healing in Δ*Fam134b/c* cells reconstituted with either FAM134B-V5 or FAM134C-V5, or empty vector control (EV), and treated with FSK (40 µM, 7 h). Quantification in (**D**) of single cells migrating into the wound (without collective invasion of neighbours) (*n* = 5 independent replicates, mean ± s.e.m., 1-way ANOVA and indicated Holm-Šídák multiple comparisons, *** = p < 0.001, * = p ≤ 0.05). Control immunoblot in (**E**) for V5, FAM134B and FAM134C shows endogenous or below endogenous levels, respectively, in the reconstitutions (WT = wild-type parental cells) (representative blot, *n* = 2 independent replicates).

**(F-G)** 2D scratch wound healing assay in PDAC-Cas9 cells expressing indicated gRNA-insensitive PRKAR1A-V5 variants (mtPS = $PKA_{CAT}$ binding-deficient), or empty vector control (EV), transfected with gRNA targeting endogenous *Prkar1a* (or NTC, non-targeting control) for 7 days before wounding and treatment with FSK (40 µM, 7 h). Quantification of single cells migrating into the wound (without collective invasion of neighbours) shown in (**F**) (*n* = ≥ 5 independent replicates, mean ± s.e.m., 2-way ANOVA and indicated Holm-Šídák multiple comparisons, *** = p < 0.001, ** = p < 0.01, ns = p > 0.05). Control immunoblot for V5 and PRKAR1A in (**G**) shows modest ectopic PRKAR1A reconstitution at levels not exceeding endogenous (representative blot, *n* = 2 independent replicates). n.s. = non-specific band. V5 = band corresponding to PRKAR1A-V5.

**(H-I)** 3D spheroid collagen matrix invasion assay (72 h) with PDAC parental (WT) or Δ*Fam134b/c* cells, untreated or treated with Adrenaline (50 µM, 72 h). Representative widefield fluorescence images of spheroids stained for F-actin (Phalloidin) shown in (**H**). White boxes delineate zoom panels. Arrowheads indicate invasive single cells (yellow, mesenchymal morphology; white, amoeboid morphology). Scale bars, 200 µm (main), 100 µm (zooms). Single cell invasion is quantified in (**I**) (*n* ≥ 15 individual spheroids from 3 independent replicates, mean ± s.d., 1-way ANOVA and indicated Holm-Šídák multiple comparisons, **** = p ≤ 0.0001, ** = p ≤ 0.01).

MW markers = kDa.

### Supplementary Table Legends

**Table S1. Proximity labelling proteomics mass spectrometry data related to Figure 1F.**

Reporting imputed LFQ values, fold-change, p-values and robust Z-scores.

**Table S2. Signal-retaining autophagy indicator (SRAI) data related to Figure 2B.**

Reporting imputed robust Z-scores.

**Table S3. Proximity labelling proteomics mass spectrometry data related to Figure 6A-B and S6A-F.**

Reporting imputed LFQ values, fold-change and p-values.

### Supplementary Movie Legends

**Movie S1. Representative FRAP of mApple-PRKAR1A signal from FAM134B$^+$ PRKAR1A$^+$ foci, related to Figure 4C.**

Photobleached area is marked by a white circle. Time lapse confocal fluorescence microscopy shows the area before (pre-bleach, 0 – 3 s)) and after photobleaching (post-bleach, 10 s onwards – recovery of fluorescence). 1 s intervals. Scale bar, 5 µm.

**Movie S2. Representative FRAP of FAM134B-EGFP signal from FAM134B$^+$ PRKAR1A$^+$ foci, related to Figure 4C.**

Photobleached area is marked by a white circle. Time lapse confocal fluorescence microscopy shows the area before (pre-bleach, 0 - 3 s) and after photobleaching (post-bleach, 10 s onwards – recovery of fluorescence). 1 s intervals. Scale bar, 5 µm.

**Movie S3. Representative surface rendering of mApple-FAM134B$^+$ LC3B$^+$ PRKAR1A$^+$ foci, related to Figure 4E.**

Representative SoRa super-resolution microscopy z-stack animation of surface rendering image in Fig. 4E. Briefly, PDAC cells expressing mApple-FAM134B (magenta) were treated with FSK (40 µM, 4 h) and co-stained with LC3B (cyan) and PRKAR1A (yellow). Scale bar, 0.5 µm.

**Movie S4. Representative surface rendering of LAMP1$^+$ PRKAR1A$^+$ foci, related to Figure 4G.**

Representative SoRa super-resolution microscopy z-stack animation of surface rendering image in Fig. 4G. Briefly, PDAC cells were treated with FSK (40 µM, 30 min) and co-stained with LAMP1 (red) and PRKAR1A (yellow). Scale bar, 0.2 µm.

**Movie S5. Wound healing assay for DMSO-treated PDAC WT cells, related to Figure 7C.** Representative phase contrast time-lapse video-microscopy, wound closure of scratched PDAC WT monolayer treated with DMSO for 22.5 h. Scale bar, 100 µm.

**Movie S6. Wound healing assay for FSK-treated WT PDAC cells, related to Figure 7C.**
Representative phase contrast time-lapse video-microscopy, wound closure of scratched PDAC WT monolayer treated with FSK (40 µM) for 22.5 h. Scale bar, 100 µm.

**Movie S7. Wound healing assay for FSK-treated PDAC Δ*Fam134b/c* cells, related to Figure 7C.** Representative phase contrast time-lapse video-microscopy, closure of scratched PDAC Δ*Fam134b/c* monolayer treated with FSK (40 µM) for 22.5 h. Scale bar, 100 µm.

**Movie S8. Wound healing assay for FSK-treated PDAC Δ*Prkar1a* cells, related to Figure 7C.** Representative phase contrast time-lapse video-microscopy, wound closure of scratched PDAC Δ*Prkar1a* monolayer treated with FSK (40 µM) for 22.5 h. Scale bar, 100 µm.

**Movie S9. Spheroid invasion assay of DMSO-treated PDAC WT cells, related to Figure 7H-J.** Representative time-lapse video-microscopy (Incucyte Brightfield) at 4 h intervals over 40 h of PDAC WT embedded in a collagen matrix and treated with DMSO (final 20 h). Scale bar, 1 mm.

**Movie S10. Spheroid invasion assay of FSK-treated PDAC WT cells, related to Figure 7H-J.** Representative time-lapse video-microscopy (Incucyte Brightfield) at 4 h intervals over 40 h of PDAC WT embedded in a collagen matrix and treated with FSK (40 µM, final 20 h). Scale bar, 1 mm.

**Movie S11. Spheroid invasion assay of FSK-treated PDAC Δ*Fam134b/c* cells, related to Figure 7H-J.** Representative time-lapse video-microscopy (Incucyte Brightfield) at 4 h intervals over 40 h of PDAC Δ*Fam134b/c* embedded in a collagen matrix and treated with FSK (40 µM, final 20 h). Scale bar, 1 mm.

**Movie S12. Spheroid invasion assay of FSK-treated PDAC Δ*Prkar1a* cells, related to Figure 7H-J.** Representative time-lapse video-microscopy (Incucyte Brightfield) at 4 h intervals over 40 h of PDAC Δ*Prkar1a* embedded in a collagen matrix and treated with FSK (40 µM, final 20 h). Scale bar, 1 mm.
